# Air-driven aldehyde synthesis in engineered bacteria via gene deletion and aryl-alcohol oxidase profiling

**DOI:** 10.64898/2026.07.12.738062

**Authors:** Roman M. Dickey, Joshua Bryan, Vishal Somasundaram, Shelby R. Anderson, Nathan Phan, Aditya M. Kunjapur

## Abstract

Engineered bacterial routes for oxidation of non-native alcohols face three challenges: Nicotinamide-dependent enzymes are coupled to cellular redox metabolism, nicotinamide-independent aryl-alcohol oxidases (AAOs) usually express poorly in bacteria, and aldehyde products are rapidly modified by host enzymes. Here, we address these limitations by engineering aldehyde-retaining *Escherichia coli* for discovery and application of soluble bacterial AAOs. Screening 51 candidates revealed a high-expression sequence cluster containing enzymes that are active on diverse aromatic and furan-based alcohols. Pairing the top-performing AAO with designer pathways in aldehyde-retaining cells enabled modular C-N and C-C bond forming cascades starting from supplied alcohols. By making both the oxidase and its product compatible with the host, this work advances air-driven oxidation of diverse alcohols as a programmable entry point to aldehyde-derived chemistry in engineered bacteria.

## Introduction

Alcohol oxidation is a common and enabling transformation in the synthesis of fine chemicals and pharmaceutical intermediates^1–5^, but its implementation is often poorly aligned with sustainability goals. For example, in a recent analysis of >400,000 medicinal chemistry reactions, alcohol oxidations were singled out among transformations that frequently rely on hazardous solvents, with dichloromethane and hypervalent iodine reagents used in 54% of surveyed alcohol oxidation examples^6^. Biocatalysis can improve the selectivity and sustainability of alcohol oxidation, and cellular biocatalyst formats, including live cells and lysates, are especially attractive because they eliminate enzyme purification and can connect oxidation to downstream multi-reaction sequences using a single strain^7–11^. However, the current state-of-the-art enzymes for alcohol oxidation are alcohol dehydrogenases (ADHs)^12,13^, whose reversible reactions rely on NAD(P)^+^/NAD(P)H pools shared with central metabolism. Even in the presence of co-factor regenerating enzymes, these systems remain coupled to an extensive network of nicotinamide co-factor consuming reactions as well as aldehyde consuming reactions, including aldehyde over-oxidation to the carboxylic acid that occurs in the presence of excess oxidized nicotinamide co-factor^8^.

Aryl-alcohol oxidases (AAOs) offer a complementary solution given their use of enzyme-bound flavin and molecular oxygen available in air to oxidize alcohols irreversibly to aldehydes without stoichiometric consumption of nicotinamide cofactors^14,15^ (**Fig. 1**). AAOs also offer theoretical advantages for aldehyde synthesis over reductive alternatives that start from carboxylic acids, such as carboxylic acid reductases (CARs)^16,17^, which require both ATP and NADPH per turnover, or aldehyde dehydrogenases (ALDHs)^18,19^, which catalyze reversible, NAD(P)H-consuming reactions. Despite their conceptual appeal, AAOs have experienced limited adoption in low-cost, bacterial cell-based biocatalyst formats due to the characteristics of known AAOs and the instability of the aldehyde product. First, nearly all characterized AAOs are secreted fungal glycoproteins that express poorly in bacterial hosts^20^. Rare exceptions that are compatible with heterologous expression in *Escherichia coli* have been found either through further exploration of fungal homologs or by examining sequences of bacterial origin that share similarity^21,22^. Among bacterial sequences, one of the first confirmed AAOs was purified from the dye decolorizing bacterium *Sphingobacterium* sp. ATM at native levels of expression, without investigation of heterologous expression^23^. In 2024, the Ferreira group reported the discovery of 2 novel bacterial AAOs that solubly express in *E. coli*, from *Sphingobacterium daejeonense* (*Sd*AAO) and *Streptomyces hiroshimensis* (*Sh*AAO)^24^. Based on their solved crystal structures, these compact and monomeric AAOs seem to possess highly accessible active sites with loop regions in distinct positions of the channel and complementary substrate preferences to fungal AAOs. However, when the substrate scope of *Sh*AAO was investigated using 60 primary and secondary alcohols, it exhibited low conversion for many tested (hetero)benzylic alcohols, with >50% conversion only observed for 2 of the 19 reactants from this category^25^. These observations suggested that a broader and more systematic bioprospecting effort could increase understanding of sequence features that increase compatibility with heterologous expression and/or improve activity on (hetero)benzylic alcohols.

**Figure 1.**
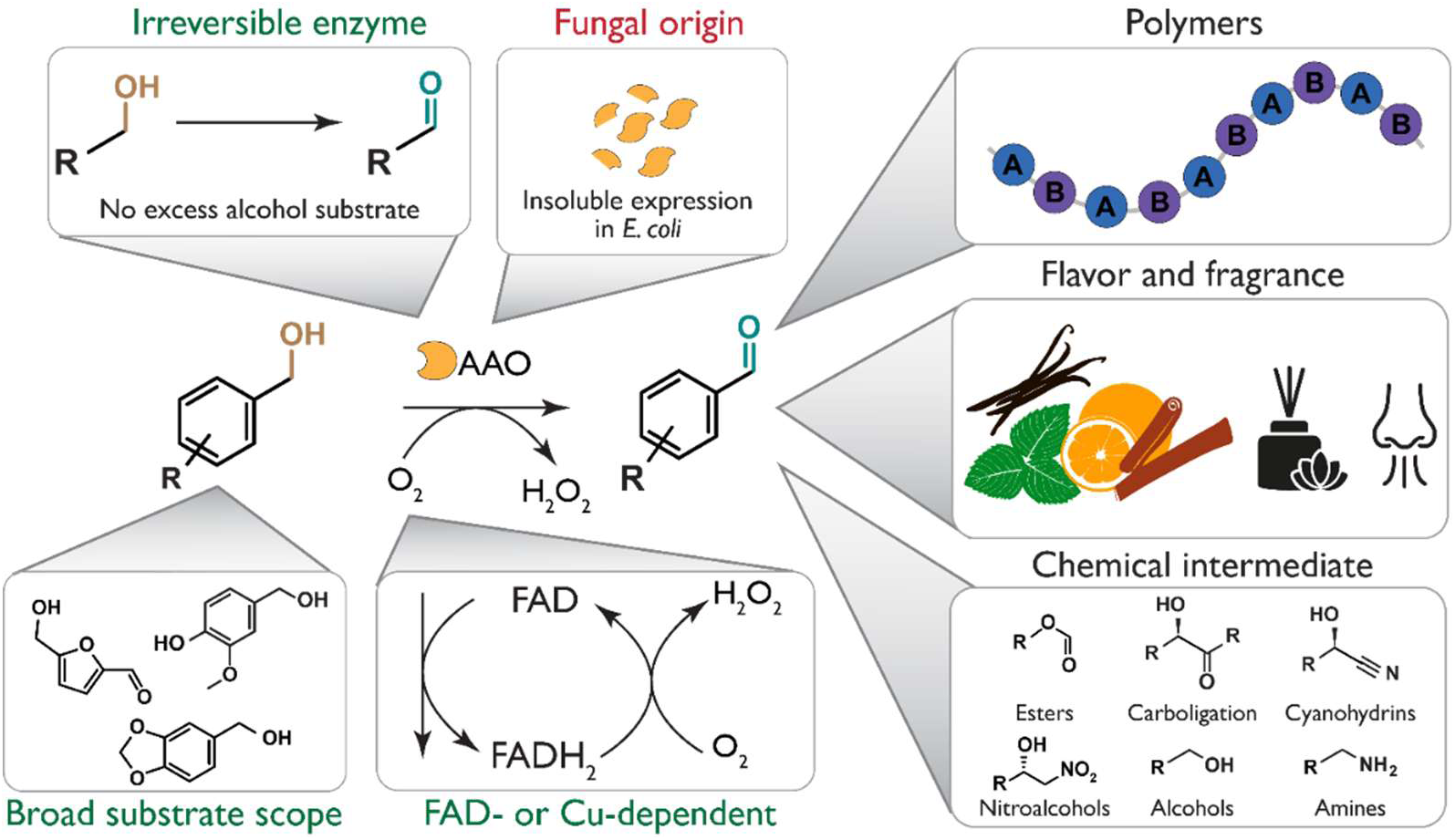
Opportunities and barriers for air-driven aldehyde synthesis in bacteria. AAOs use enzyme-bound flavin and molecular oxygen to oxidize alcohols to aldehydes, avoiding direct dependence on cellular nicotinamide redox cycling and favoring aldehyde formation over the reversible chemistry of alternative enzymes such as ADHs. In engineered bacteria, this chemistry could provide access to aldehydes as products or as intermediates for polymer, flavor and fragrance, and chemical intermediate synthesis. However, bacterial implementation requires solving three incompatibilities: ADH-based oxidation is coupled to cellular redox metabolism, many characterized AAOs are fungal enzymes that express poorly in *E. coli*, and aldehyde products are rapidly modified by endogenous host enzymes.

Aldehyde instability in cellular settings presents a second barrier to using AAOs in biochemical pathways. In wild-type bacteria, native ADHs, ALDHs and aldo-keto reductases rapidly erase aldehydes by reduction or oxidation, making intracellular AAO activity difficult to detect and difficult to connect to downstream chemistry. We and others have shown that multiplex deletion of aldehyde-consuming enzymes can stabilize diverse aldehydes under growing and resting-cell conditions^8,9,26,27^. This suggested that aldehyde-retentive bacteria could do more than host AAOs: they could reveal soluble AAOs during discovery and preserve their products for subsequent C-N and C-C bond formation. Here we report a paired enzyme-discovery and host-engineering strategy for air-driven aldehyde synthesis in living bacteria. From 51 bacterial AAO candidates, we identified an expression-enriched sequence cluster with predicted active-site architectures distinct from characterized bacterial AAOs. Four soluble homologs oxidized diverse aromatic and furan-based alcohols, and the top enzyme enabled modular cascades that converted alcohol substrates to representative amines and non-standard amino acids. These findings establish bacterial AAOs as genetically tractable, air-driven aldehyde generators and show that host aldehyde stabilization can transform alcohol oxidation into a programmable entry point for whole-cell synthesis.

## Results

### Aldehyde-retaining *E. coli* preserves diverse target aldehydes

We first asked whether past and newly engineered *E. coli* strains could stabilize a broad set of aromatic and heterocyclic aldehydes relevant to AAO-mediated alcohol oxidation to allow enzyme discovery in live cells. We previously showed that the *E. coli* ROAR strain, denoted here as the ROAR.Δ12 strain, improves stability of 8 supplemented aldehydes in resting cells, a format in which aldehydes are otherwise vulnerable to both reduction and oxidation^28^. In parallel work, we showed that the *E. coli* RARE.Δ16 strain prevents reduction of terephthalaldehyde in growing cells^9^. We later introduced 10 gene deletions to generate the *E. coli* ROAR.Δ22 strain for a terephthalaldehyde-dependent cascade in resting cells^29^, but the broader aldehyde scope of this strain had not been evaluated. Thus, we performed an extensive comparison of the wild-type MG1655, RARE.Δ16, ROAR.Δ12, and ROAR.Δ22 strains (**Fig. 2A,B**).

**Figure 2.**
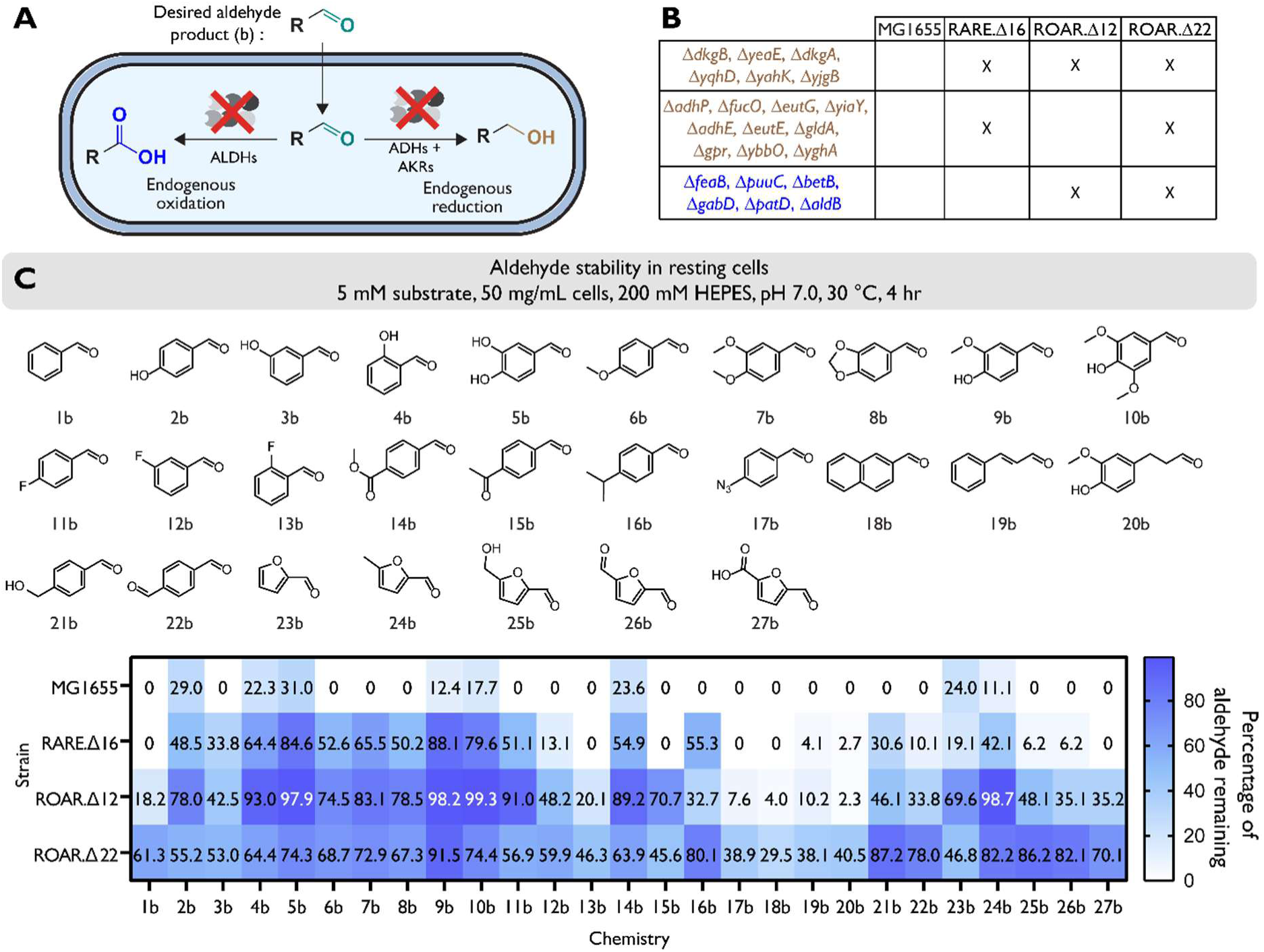
Aldehyde stability in engineered *E. coli* strains. (A) Resting cells with endogenous genes were knocked out to stabilize aldehyde products. (B) Diagram of the knockouts made in each engineered strain tested for aldehyde stability. (C) We tested 23 chemistries with variations in the position, size, and electrostatic properties of aryl ring functional groups as well as furan and bicyclic aromatic compounds. We supplemented 5 mM of each aldehyde to resting cells of MG1655, RARE.Δ16, ROAR.Δ12 and ROAR.Δ22. The percentage of aldehyde remaining was measured after 4 h using HPLC compared to a no cell control to account for mass loss due to aldehyde volatility. Samples sizes are *n* = 3 using technical replicates. Data shown are the mean conversion.

We supplied resting cells of each strain with a panel of 27 structurally diverse aldehydes and quantified aldehyde retention after 4 h. As expected, MG1655 rapidly consumed most substrates, leaving no detectable aldehyde for 19 of 27 compounds (**Fig. 2C**). Removing reductive activities alone improved but did not fully address retention, as RARE.Δ16 fully depleted only 6 of 27 aldehydes. However, only 3 of 27 compounds exhibited >70% retention, indicating significant aldehyde instability remained. In contrast, ROAR.Δ12 and ROAR.Δ22 substantially broadened aldehyde retention across the panel. For every aldehyde tested, at least 30% remained in the best-performing ROAR background, with 15 of 27 compounds showing >70% retention. ROAR.Δ22 generally provided the highest retention, particularly for aldehydes poorly stabilized by ROAR.Δ12. However, ROAR.Δ12 outperformed ROAR.Δ22 for a few substrates, indicating that additional deletions do not uniformly improve aldehyde stability and may alter compensatory oxidoreductase activity or cellular redox balance. These results demonstrate the ability of the *E. coli* ROAR hosts to preserve aldehydes that could be generated by AAOs, suggesting their suitability for cell-based AAO discovery screening.

### Profiling candidate bacterial AAOs in aldehyde-retaining cells reveals an expression hotspot

To discover AAOs compatible with live-cell aldehyde synthesis, we searched broadly across AAO-like Glucose-Methanol-Choline (GMC) oxidoreductase sequence space (**Fig. 3A,B**). We generated a protein sequence similarity network (SSN) with over 4,000 sequences using BLASTP searches seeded with characterized fungal AAOs from *Komagataella phaffii*^30,31,3^ and *Aspergillus terreus*^32–34^ (AtAAO), together with bacterial GMC oxidoreductases from *Rhodococcus sp*. strain RHA1 (RjAAO) and *Pseudarthrobacter chlorophenolicus* (PclAAO)^35^. We included fungal sequences to maximize biological diversity, but experimental screening focused on bacterial candidates to maximize the likelihood of soluble expression in *E. coli*. This broader search also avoided imposing conservation of two active-site histidines used to constrain a prior bacterial AAO discovery effort^24^. The SSN contained distinct fungal and bacterial regions, including several bacterial clusters without characterized AAOs. We selected 29 bacterial candidates from 10 clusters for first-round evaluation and included the fungal AtAAO as a control, with pairwise sequence identities ranging from 12 to 88%. Western blot analysis of soluble lysate fractions obtained after expression of His-tagged proteins showed detectable soluble expression for 16 candidates (**Fig. 3C**). Expression was not evenly distributed across the SSN: clusters containing a multispecies *Burkholderia* GMC oxidoreductase (BuAAO), a metagenome-derived oxidoreductase from a marine Gammaproteobacteria bacterium (GbAAO), or the previously characterized PclAAO were enriched for soluble homologs.

**Figure 3.**
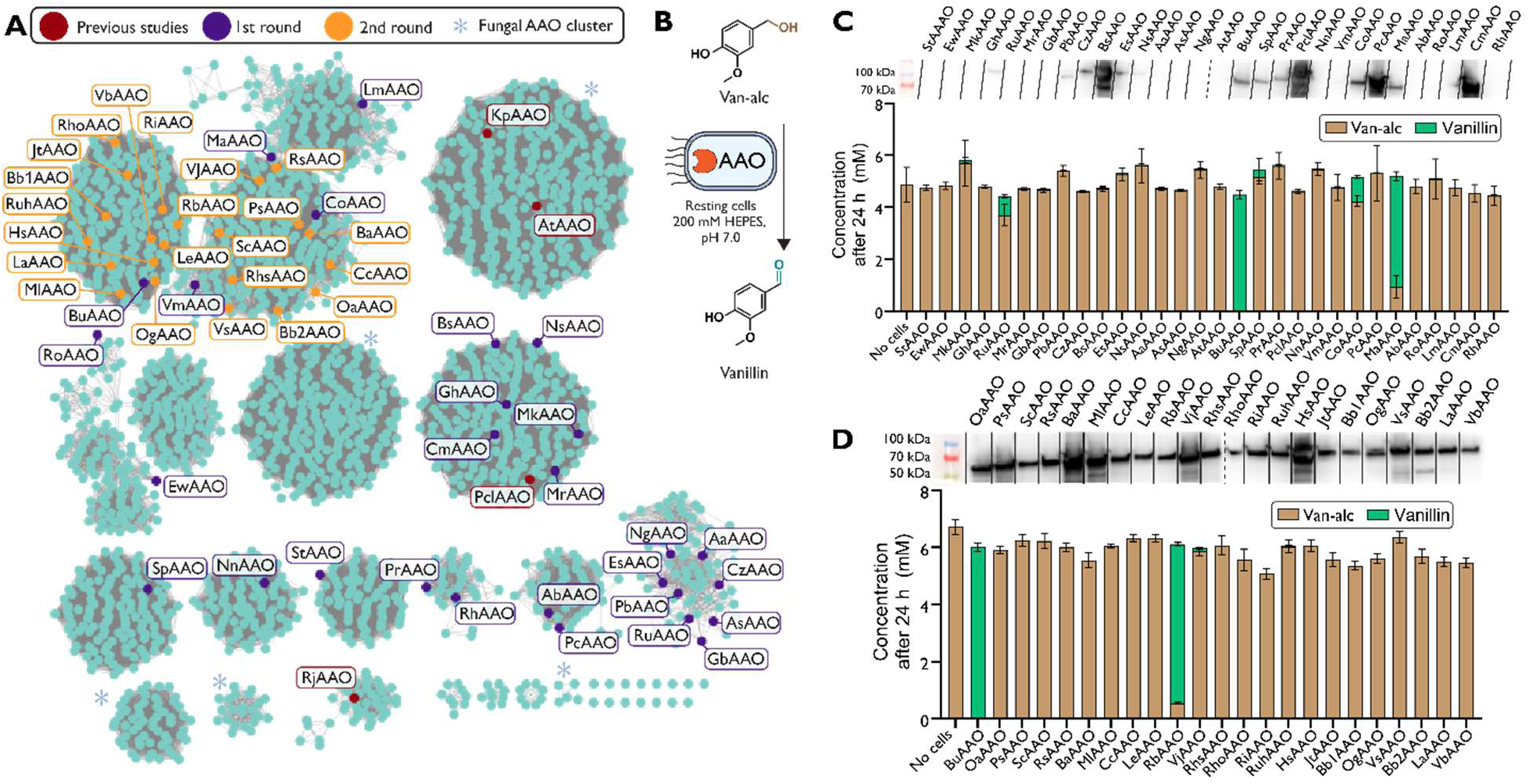
Bioprospecting, expression and whole cell activity of putative aryl-alcohol oxidases. (A) A Protein Sequence Similarity Network (SSN) containing 2,154 sequences related to KpAAO, AtAAO, RjAAO and PclAAO with selected putative AAOs highlighted in purple (1^st^ round), orange (2^nd^ round) and the characterized enzymes in the literature highlighted in red. Edges are drawn between nodes with a minimum alignment score of 100. (B) Resting whole cell activity assay of all AAOs via supplementation of 5 mM of vanillyl alcohol to resting cells expressing each AAO in ROAR.Δ12. Solubility was tracked via western blot and activity via endpoint conversion after 24 h for both the (C) first and (D) second round of selected AAOs with the top performing AAO (BuAAO) included in the second round for comparison. Samples sizes are *n* = 3 using technical replicates. Data shown are mean ± s.d.

We next asked whether soluble expression translated into alcohol oxidation activity. We chose vanillyl alcohol as our model substrate because its aldehyde product, vanillin, is well retained in ROAR.Δ12 and is an industrially relevant flavor and fragrance compound. We prepared ROAR.Δ12 resting cells expressing each candidate AAO at 50 mg wet cell weight/mL in 200 mM HEPES at pH 7.0 and supplied 5 mM vanillyl alcohol. After 24 h, most candidates showed little to no detectable conversion by HPLC (**Fig. 3C**). However, three homologs generated vanillin: BuAAO, a compost metagenome oxidoreductase (CoAAO), and a GMC oxidoreductase from an unclassified Massilia (MaAAO). MaAAO converted 82% of the applied substrate, whereas BuAAO gave complete conversion under the conditions tested. Notably, all three active homologs came from the same SSN cluster.

We therefore expanded sampling within this cluster to include 22 additional candidate sequences, resulting in a total of 51 bacterial AAO-like candidates. All second-round candidates exhibited high levels of soluble expression, confirming that this region of the SSN represents an expression-enriched bacterial AAO hotspot (**Fig. 3D**). Once again, high expression alone did not guarantee activity on vanillyl alcohol, as only one of the second-round homologs converted vanillyl alcohol detectably. This was the GMC oxidoreductase from Rhizobiaceae bacterium (RbAAO), which reached 91% conversion under conditions identical to the prior experiment. BuAAO remained the highest-performing enzyme in the expanded screen. Together, these results show that aldehyde-retentive cells can support HPLC-based discovery of intracellular AAO activity and identify an expression-enriched bacterial sequence cluster containing rare and highly active AAOs.

### Sequence and structure analyses show distinct features of the expression-enriched AAO cluster

We briefly investigated whether the expression-enriched cluster contained features associated with soluble expression or activity on vanillyl alcohol. Two protein solubility predictors, NetSolP^36^ and SoluProt^37^, generated output that did not correlate well with soluble expression scores obtained from Western blot image analysis (**Supplementary Table 1**), suggesting that generic solubility models were not sufficient for this enzyme family. We therefore aligned the tested sequences by using MAFFT^38^, calculated pairwise identities, and performed hierarchical clustering (**Supplementary Figure 1**). Partitioning the alignment into seven clusters identified one cluster in which 26 of 30 sequences exhibited high soluble expression (≥ 75% of the highest intensity band) and a mean expression level of 80.3% for the full cluster. We compared the 26 highly expressed members of this cluster with all other tested sequences to identify alignment positions associated with expression. After gap filtering and Jensen-Shannon divergence^39^ analysis, 16 positions most strongly differentiated the groups (**Supplementary Table 2**). The high-expression group showed strong consensus at these positions, whereas the four low-expression sequences that still fell within the same cluster deviated from the consensus at seven or more of these positions. Most of these expression-associated positions involved changes in amino acid property between groups and mapped to the surface of an AlphaFold^40^-predicted structure of VjAAO, one of the most highly expressed homologs (**Fig. 4A,B**).

**Figure 4.**
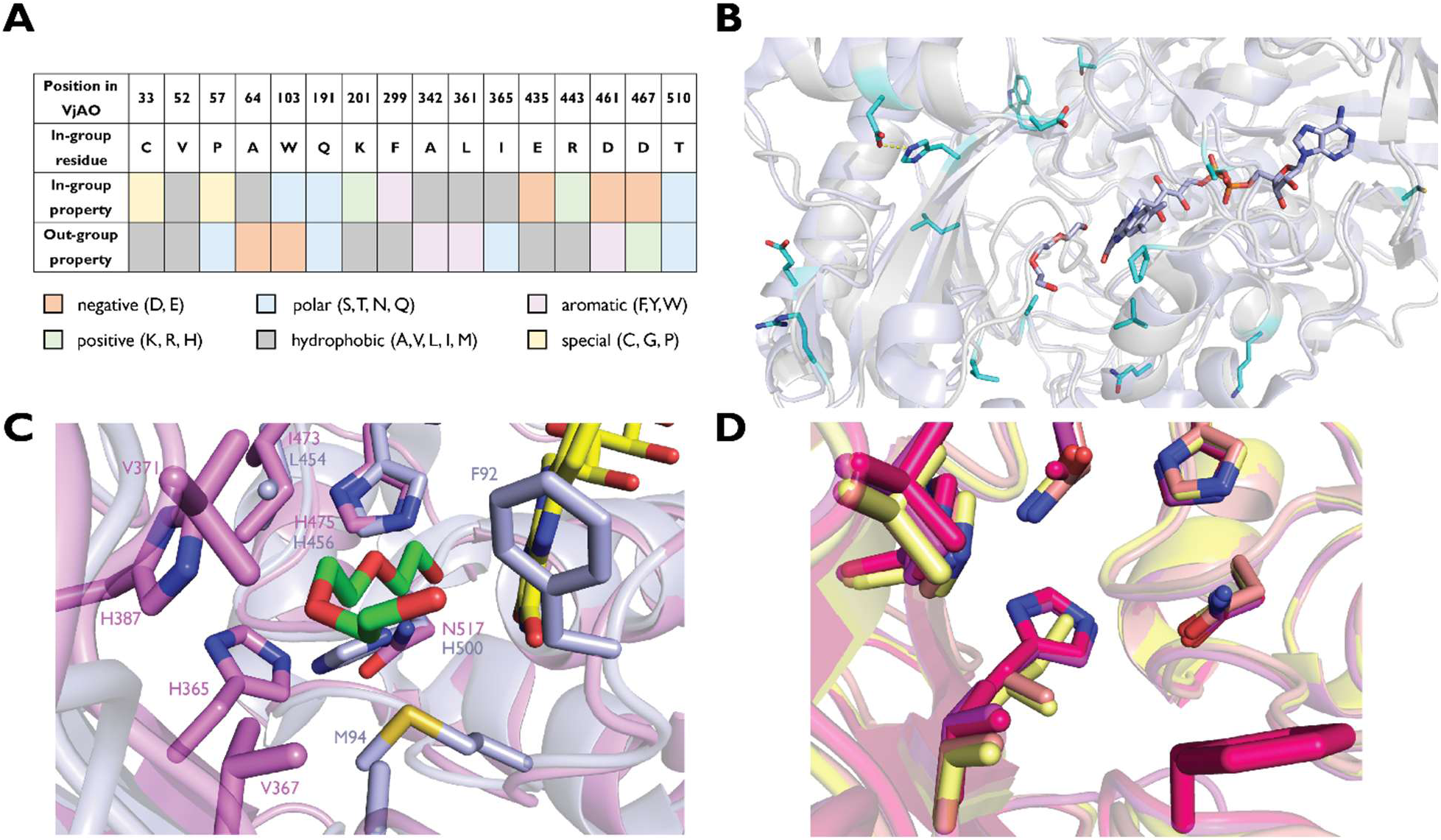
Comparative analysis of AAOs reveals features associated with expression and activity. (A) The 16 most divergent positions in terms of dominant residue identity between the in-cluster and high-expression group (in-group) and all others (out-of-group), as ranked by Jensen-Shannon divergence followed by manual curation. Listed are the position numbers with respect to a cluster representative that exhibited 100% relative soluble expression (VjAAO) and the dominant in-group residue (which was not always shared by VjAAO). The bottom two rows reflect a more chemically aware analysis that examines the property of the dominant residue in each group and showcases larger changes more clearly. (B) Illustration of the predicted positions of the top 16 residues using an AlphaFold-predicted structure of VjAAO (transparent, gray) aligned to the previously obtained crystal structure of ShAAO (PDB ID 8RPG, transparent light blue). The residues of VjAAO are colored in cyan. The distance between two residues predicted to be proximal (D461 and H299) and that may form a salt bridge is shown by a dashed line (2.9 Å). The bound alcohol substrate and FAD from 8RPG are shown for reference. (C) Active site comparison of BuAAO (magenta) and ShAAO (light blue), with residues within 4 Å of the alcohol substrate shown as sticks and labeled. (D) Active site comparison among the 4 most active bacterial AAOs discovered in this study. Colors: BuAAO, magenta; RbAAO, hot pink; MaAAO, salmon; CoAAO, yellow.

We also compared AlphaFold-predicted structures of the four active homologs with the two bacterial AAOs whose structures were recently reported. We focused on ShAAO (PDB ID 8RPG) given its distinct substrate scope compared to fungal AAOs. In contrast to prior sequences that contain His at two conserved positions in the active site (H456 and H500, 8RPG numbering), our AAOs contain only one of these His, with the other (H500) replaced by Asn, suggesting that our hits occupy a distinct region of bacterial AAO-like sequence space. Structural alignment of BuAAO to ShAAO reveals a large predicted shifted in a loop region near the substrate binding pocket (**Supplementary Figure 2A,B**) and a more open FAD-proximal region (**Fig. 4C**). Comparison of the four most active homologs further highlighted two positions near the predicted pocket, corresponding to BuAAO H365 and I473, that differed between the more and less active enzymes (**Fig. 4D**). Together, these computational analyses support the view that the expression-enriched cluster is distinct from characterized bacterial AAOs, with some identified features that may guide future engineering.

### Novel AAO homologs efficiently oxidize diverse aromatic and heterocyclic alcohols in cells

Given that many aromatic and heterocyclic aldehydes were retained by ROAR.Δ22, we proceeded to examine whether the four active homologs could generate this product space directly from alcohol substrates in resting cells (**Fig. 5A**). We selected 27 alcohols whose products span flavor and fragrance compounds^41^, pharmaceutical precursors^42–44^, common synthetic intermediates^45,46^, and monomers^47–53^. At 5 mM substrate loading, the four homologs gave generally high conversion after 4 h at 30 °C. At least one homolog gave complete or near-complete conversion for 19 of 27 substrates, and 25 substrates reached over 70% conversion (**Fig. 5B**). Although the four enzymes showed overlapping substrate profiles, BuAAO was the most broadly active homolog under these conditions, giving the highest conversion for most substrates in the panel. The results showed that the active bacterial homologs are not limited to vanillyl alcohol and can support air-driven formation of diverse aldehydes in aldehyde-retaining cells.

**Figure 5.**
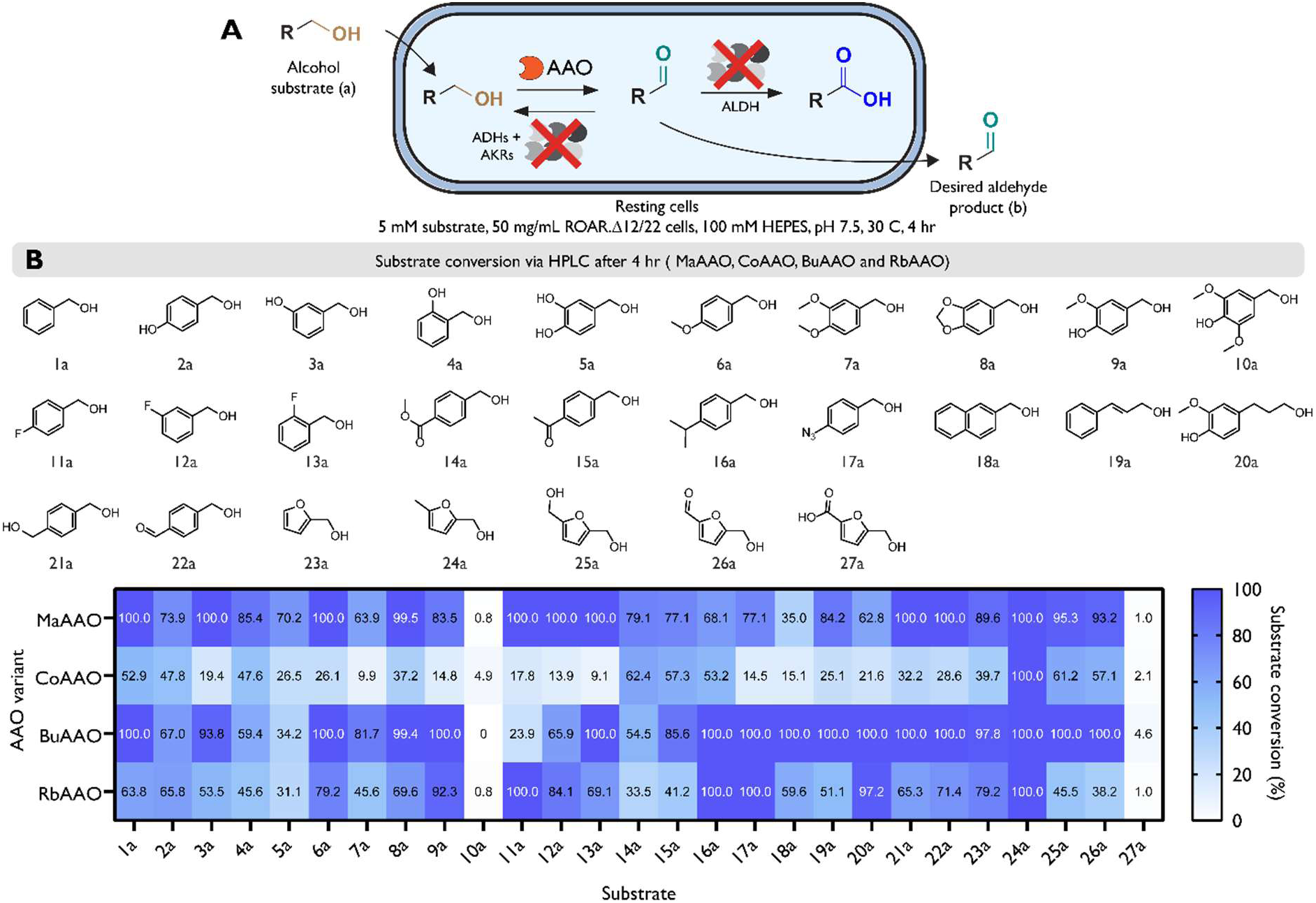
The top four newly discovered bacterial AAOs exhibit high activity on a broad range of benzyl and furan-based alcohols. (A) Resting whole cell assay of 4 active AAOs (MaAAO, CoAAO, BuAAO and RbAAO) via supplementation of 5 mM of alcohol substrates to resting cells expressing each AAO in the best performing strain (ROAR.Δ12 or ROAR.Δ22). (B) We tested 27 chemistries with variations in the position, size, and electrostatic properties of aryl ring functional groups as well as furan and biphenyl compounds. The conversion was measured after 4 h using HPLC. Samples sizes are *n* = 3 using technical replicates. Data shown are the mean conversion.

Because BuAAO gave the highest overall performance in the endpoint substrate screen, we next followed whole-cell reaction kinetics with vanillyl alcohol and piperonyl alcohol (**Fig. 6A**). At 5 mM substrate loading, both reactions proceeded rapidly in resting ROAR.Δ12 cells expressing BuAAO. Vanillyl alcohol reached 78% conversion after 15 min and complete conversion after 45 min (**Fig. 6B**). Piperonyl alcohol reached 92% conversion after 15 min and full conversion after 30 min (**Fig. 6C**). We then investigated conversion at increased substrate loadings. At 10, 20, and 40 mM substrate, BuAAO expressing cells produced vanillin at 92%, 80% and 74% conversion after 4 h, respectively, and 98%, 95% and 90% conversion after 20 h (**Fig. 6D,E**). These reactions were performed in aqueous whole cell format without an organic overlay, without exogenous nicotinamide cofactors, and without additional carbon sources for cofactor regeneration. Thus, pairing BuAAO with the ROAR strains enables high-loading aldehyde formation under simple aerobic resting cell conditions.

**Figure 6.**
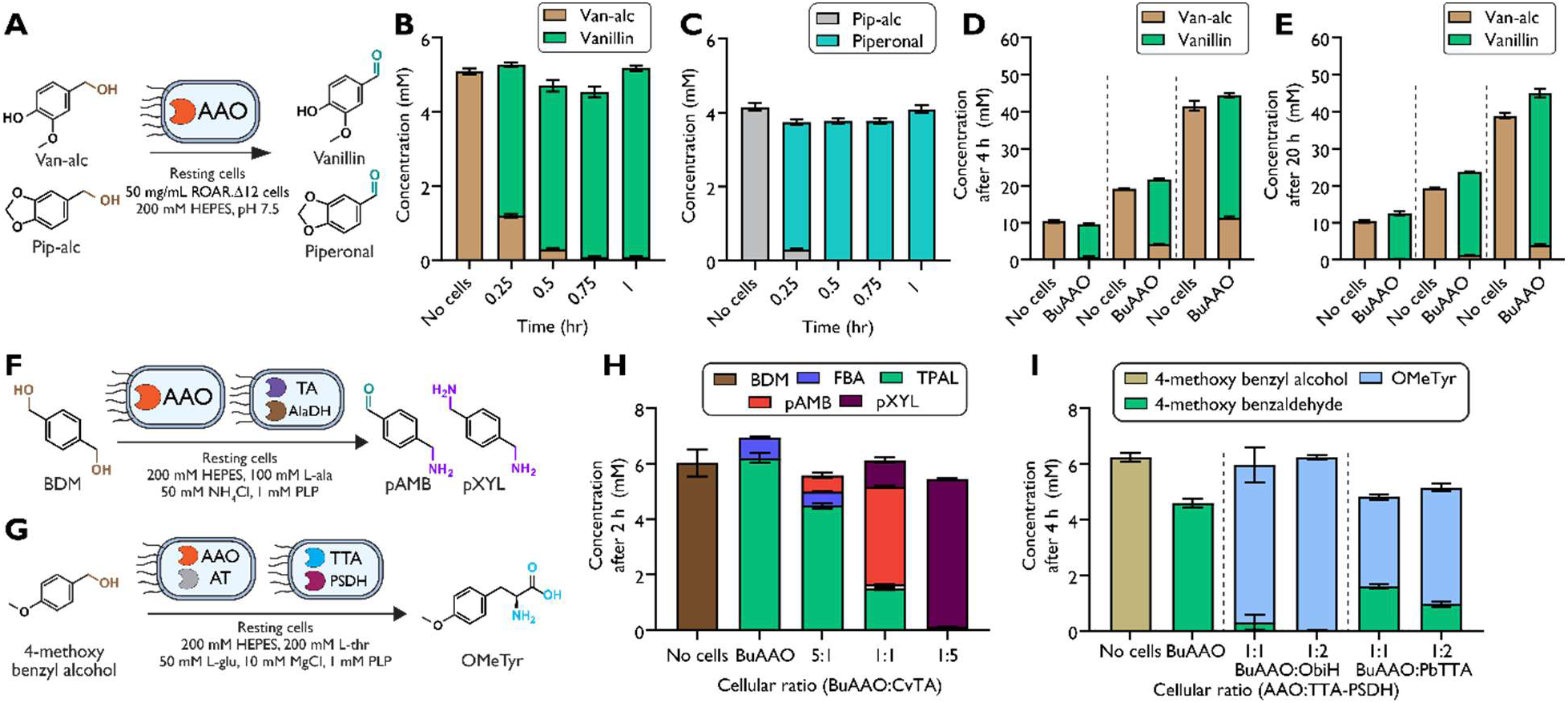
Resting cells expressing BuAAO exhibit fast reaction kinetics at higher substrate loadings and enable modular cascades for C-N and C-C bond formation. (A) Resting whole cell assay of BuAAOs in ROAR.Δ12 via supplementation of 5 mM of vanillyl alcohol or 5 mM of piperonyl alcohol to resting cells. Conversion was measured over 1 h with timepoints taken every 0.25 h after supplementation with (B) vanillyl alcohol or (C) piperonyl alcohol. Resting cell assays using BuAAO with increased substrate loading (10 mM, 20 mM and 40 mM) were performed with samples taken at (D) 4 h and (E) 20 h. (F) Reaction diagram and conditions for diamine production. An ω-TA from *Chromobacterium violaceum* (CvTA) was used for amination and an alanine dehydrogenase from *Bacillus subtilis* (AlaDH) was used for cofactor recycling (G) Reaction diagram and conditions for nsAA production. A SUMO tagged PSDH gene from *Ralstonia pickettii* (PSDH) as well as a SUMO tagged TTA from *Pseudomonas fluorescens* (ObiH) or Parachlamydiales bacterium (PbTTA) were used for production of nsAAs. (H) The resting cell reaction was performed by supplementing 6 mM of 1,4 benzenedimethanol (BDM) to a 200 mM HEPES, 100 mM L-alanine, 50 mM NH_4_Cl, and 1 mM PLP buffer containing 100 mg wet cell weight/ml of ROAR.Δ22 cells expressing either BuAAO or CvTA-AlaDH at different cellular ratios (5:1, 1:1 or 1:5) (I) The resting cell reaction was performed by supplementing 6 mM of 4-methoxy benzoic acid to a 200 mM HEPES, 200 mM L-threonine, 50 mM L-glutamate, 10 mM MgCl and 1 mM PLP buffer containing 100 mg wet cell weight/ml of ROAR.Δ22 cells expressing either BuAAO, ObiH-PSDH or PbTTA-PSDH at different cellular ratios (1:1 or 1:2). Samples sizes are *n* = 3 using technical replicates. Data shown are mean ± s.d.

### Air-driven alcohol oxidation feeds C-N and C-C bond forming pathways in live cells

Finally, to illustrate the breadth of products that can be produced from AAO coupled reactions, we examined the biosynthesis of a diamine polymer target *para*-xylylenediamine (pXYL) as well as the non-standard amino acid O-methyl-L-tyrosine (OMeTyr) from their associated alcohols (1,4-benzenedimethanol and 4-methoxybenzyl alcohol respectively) (**Fig. 6F, G**). The synthesis of pXYL features a model C-N bond forming reaction catalyzed by an ω-transaminase using a biosynthesized aldehyde, whereas the synthesis of OMeTyr features a model C-C bond forming reaction catalyzed by an L-threonine transaldolase (L-TTA) using a biosynthesized aldehyde, thereby highlighting the potential value and versatility of integrating AAOs into biochemical pathways in live cells that stabilize aldehydes.

For pXYL production, we tested an ω-transaminase from *Chromobacterium violaceum* (cvTA) which has been previously demonstrated to act upon an array of aromatic aldehydes including the aldehyde intermediate, terephthalaldehyde^54^. We chose L-alanine as our amine donor and co-expressed an alanine dehydrogenase (AlaDH) from *Bacillus subtilis* for cofactor recycling as it is capable of recycling the pyruvate product back to L-alanine. We prepared resting whole cells of BuAAO and CvTA in separate ROAR.Δ22 cells. After we harvested cells, we resuspended these cells in a buffer consisting of 200 mM HEPES, 100 mM L-alanine, 50 mM NH_4_Cl and 1 mM PLP at pH 7.5. We ran the reactions at a total cell loading of 100 mg/mL wet cell weight at BuAAO:CvTA cell ratios of 5:1, 1:1 and 1:5. We began the reactions by supplementing 5 mM of 1,4-benzenedimethanol substrate. After 2 h, at high BuAAO loadings (5:1) we did not observe detectable pXYL production (**Fig. 6H**). However, as we increased the CvTA cell ratio, we obtained greater pXYL formation. At a 1:5 BuAAO to CvTA cell ratio, we saw complete conversion towards our desired pXYL product. Despite endogenous aldehyde oxidation to 4-formylbenzoic acid (FBA) being observed in our AAO-only and 5:1 ratio reactions, we observed no oxidation products when CvTA loading was high. These findings demonstrate that, in our engineered strains, downstream enzymatic pathways can efficiently convert aldehydes into alternative products, even in the presence of residual endogenous oxidation.

For OMeTyr production, we coupled our AAO reactions to our recently designed nsAA semi-synthesis pathway consisting of an L-TTA, a phenylserine dehydratase (PSDH), and endogenous aminotransferase^55,56^. We used an expression plasmid where the PSDH gene from *Ralstonia pickettii* and either the L-TTA from *Pseudomonas fluorescens* (ObiH) or *Parachlamydiales bacterium* (PbTTA)^57^ were tagged using a Small Ubiquitin Modifier (SUMO) tag. We prepared resting whole cells of BuAAO and L-TTA-PSDH in separate ROAR.Δ12 cells. We resuspended these cells in a buffer consisting of 200 mM HEPES, 200 mM L-threonine, 50 mM L-glutamate, 10 mM MgCl_2_ and 1 mM PLP at pH 7.5. We then ran the reactions at a total cell loading of 100 mg/mL wet cell weight at BuAAO:L-TTA-PSDH cell ratios of 1:1 and 1:2. We began the reactions by supplementing 5 mM of 4-methoxybenzyl alcohol substrate. After 4 h, we observed that our reactions were L-TTA-PSDH limited as we observed unreacted 4-methoxybenzaldehyde in both ObiH and PbTTA reactions (**Fig. 6I**). However, in the case where the relative amount of L-TTA-PSDH cells were twice that of the BuAAO cells, we observed complete conversion of the alcohol to our non-standard amino acid (nsAA) OMeTyr product. Given the promiscuity of BuAAO observed in this work, along with our previous findings of high polyspecificity among the L-TTA, PSDH, and endogenous aminotransferases, this cascade has potential to produce other L-phenylalanine derivatives from their corresponding aromatic alcohols.

## Discussion

Our work systematically analyzed the recently discovered class of bacterial AAOs using strains of *E. coli* that were engineered for aldehyde stabilization. Aldehyde stabilizing strains such as ROAR.Δ12 and ROAR.Δ22 represent an enabling technology for the identification, characterization, and application of AAOs in simple cell-based formats. The ROAR.Δ12 and ROAR.Δ22 strains can stabilize a wide range of aromatic and furan aldehydes, with 15 out of the 23 tested compounds having over 70% of the aldehyde remaining after 4 h. Equipped with these strains, we performed genome mining to ultimately screen 51 candidate bacterial AAOs for their soluble expression level and their activity on vanillyl alcohol. We were excited to discover a previously unknown cluster of bacterial AAOs whose mean expression level was >80% relative to the highest expressing sequence. In this cluster were 4 highly active bacterial AAOs that were capable of converting a broad range of aromatic and furan substrates. We achieved over 70% conversion on 25 of our tested 27 chemistries with one or more of our tested AAOs in a resting whole cell reaction. We also showed that these AAO whole cell reactions are rapid, easily scalable, and can be coupled to downstream enzymes for the production of a diverse set of alternative products. Notably, we demonstrated that the level of aldehyde stability currently achieved allows us to efficiently couple aldehyde generation with downstream enzymatic pathways with no observed shunt oxidative products.

Given our encouraging findings with respect to bacterial AAO expression and activity, we conducted extensive analyses using alignments of primary sequence and predicted structures. By comparing the dominant residue identity for the in-cluster and high-expression sequence group (26 members, 92.3% mean expression level) against the others (26 members, 22.9%) at every MSA alignment position, sorting by Jensen-Shannon divergence, and then curating by eye, we found 16 position-identity pairs that are associated with high expression. Additionally, we found that our active AAOs have distinct binding pocket access, geometry, and active site residues compared to the few known bacterial AAOs. Finally, we identified subtle but potentially important active site differences among our 4 most active AAOs. Collectively, these observations substantially increase our understanding of bacterial AAOs and should have value for future efforts related to additional genome mining or AAO engineering through targeted mutagenesis or generative design.

The use of AAOs in modular cascade or pathway development is particularly attractive given the reactivity and toxicity of aldehydes, the ubiquity of aromatic and heterocyclic alcohols, and the relative orthogonality of the reaction catalyzed due to the lack of depletion of a stoichiometric, nicotinamide-based redox co-factor. Here, we demonstrated a proof-of-concept of this vision by coupling the AAO with downstream enzymes in our aldehyde stabilizing bacteria to biosynthesize higher-value products created via C-N and C-C bond formation. TAs have been extensively used for the green production of highly enantiomerically enriched amines. This AAO and TA coupled reaction could be utilized for the production of diamine polymer building blocks such as pXYL^58,59^, capsaicin analogues from lignin-derived compounds^60–62,11^, pharmaceuticals^63^, and others^64^. As a large section of the tested chemistries can be derived from lignin feedstocks, AAO-mediated oxidation may create alternative routes for lignin valorization towards flavor and fragrance compounds or other aldehyde derived products^65^. Additionally, we have recently shown that the production of nsAAs through a biocatalytic cascade consisting of a TTA, PSDH and endogenous TA works on a wide range of chemistries, and many of the associated alcohols are substrates that one or more of our discovered AAOs act upon^55^. Thus, this work provides proof-of-concept that L-phenylalanine derivatives can be prepared from aromatic alcohol substrates. Overall, by making both the oxidase and the aldehyde intermediate compatible with the cellular environment, our approach advances air-driven oxidation of diverse alcohols as a programmable route to aldehyde-derived chemistry in living bacteria.

## Acknowledgements

The authors would like to thank the members of the University of Delaware Center for Plastics Innovation for guidance and support on this project.

## Author Contributions

Conceptualization: RMD, AMK

Data curation: RMD, JB, VS, SA, NP, AMK

Methodology: RMD, JB, VS, SA, NP

Investigation: RMD, JB, VS, SA, NP, AMK

Visualization: RMD, AMK

Funding acquisition: AMK

Project administration: AMK

Supervision: AMK

Writing – original draft: RMD, AMK

Writing – review & editing: RMD, AMK, JB, VS, SA, NP

## AI Statement

AlphaFold was used for protein structure prediction central to structural analyses. Most comparative analyses performed in this study were supported by code that was developed with the assistance of ChatGPT, manually inspected, and then adjusted when necessary.

## Methods

### Strains and plasmids

*Escherichia coli* strains and plasmids used are listed in Supplementary Table 3. Molecular cloning and vector propagation were performed in DH5α (NEB). Polymerase chain reaction (PCR) based DNA replication was performed using KOD XTREME Hot Start Polymerase (MilliporeSigma) for plasmid backbones. Cloning was performed using Gibson Assembly. Oligos for PCR amplification and translational knockouts are shown in Supplementary Table 4. Oligos were purchased from Integrated DNA Technologies (IDT). The DNA sequence and translated sequence of proteins overexpressed in this paper are found in Supplementary Table 5.

### Materials and chemicals

The following compounds were purchased from MilliporeSigma: sodium borate decahydrate, sodium phosphate dibasic anhydrous, chloramphenicol, kanamycin sulfate, dimethyl sulfoxide (DMSO), boric acid, L-alanine, and HEPES. D-glucose and m-toluic acid. The following compounds were purchased from Alfa Aesar: agarose and ethanol were purchased. The following compounds were purchased from Fisher Scientific: isopropyl ß-D-1-thiogalactopyranoside (IPTG), acetonitrile, sodium chloride, trifluoroacetic acid, LB Broth powder (Lennox), and LB Agar powder (Lennox). A MOPS EZ rich defined medium kit was purchased from Teknova. Taq DNA ligase was purchased from GoldBio. Anhydrotetracycline (aTc) was purchased from Cayman Chemical. Phusion DNA polymerase and T5 exonuclease were purchased from New England BioLabs (NEB). Sybr Safe DNA gel stain was purchased from Invitrogen. The following compounds were purchased from TCI America: Pyridoxal 5’-phosphate (PLP), *ortho*-phthalaldehyde and 3-mercaptopropionic acid. The following were purchased from TCI America: D-glucose, 4-acetylbenzoic acid, 4-azidobenzoic acid, 2-naphthoic acid, terephthalaldehyde, biphenyl-4-carboxaldehyde, 1-naphthaldehyde. The following were purchased from Thermo Scientific Chemicals: 2-fluorobenzoic acid, 3-fluorobenzoic acid, 4-fluorobenzoic acid, 1-naphthoic acid, 4-benzoylbenzoic acid, 3-fluorobenzaldehyde, and 4-fluorobenzaldehyde. The following were purchased from Sigma-Aldrich: benzaldehyde, terephthalic acid, 4-methoxybenzoic acid, biphenyl-4-carboxylic acid, 4-anisaldehyde, and 2-naphthaldehyde. Agarose, ethanol, 2-fluorobenzaldehyde, L-glutamic acid monopotassium salt monohydrate, and L-threonine were purchased from Alfa Aesar. Taq DNA ligase was purchased from GoldBio. 4-cyanobenzoic acid, and 4-azidobenzaldehyde were purchased from ChemCruz.

### Culture conditions and resting cell preparation

Cultures were grown in LB-Lennox medium (LB: 10 g/L bacto tryptone, 5 g/L sodium chloride, 5 g/L yeast extract) or MOPS EZ rich defined media (Teknova M2105) with 2% glucose (MOPS media).

To prepare cells for resting cell assays, confluent overnight cultures of *E. coli* strains were used to inoculate 50 mL cultures in LB media with appropriate antibiotics (AAO: 50 µg/mL kanamycin; TA: 34 µg/mL chloramphenicol; TTA-PSDH: 50 µg/mL carbenicillin) in 250 mL baffled shake flasks. The cultures were grown at 37 °C until mid-exponential phase (OD_600_ = 0.5–0.8) and then induced (AAO: 0.1 µg/mL aTc; TA and TTA-PSDH: 1 mM IPTG). After induction, the cultures were moved to 18 °C for 18 h. Cells were then pelleted and used or frozen at −80 °C. Cells used in this study were stored at -80 for less than 24 h. Cells were washed twice with 200 mM HEPES, pH 7.0 buffer. After washing, the mass of cell pellets was then measured and resuspended in the desired reaction buffer.

### Curation of Sequence Similarity Network

To generate the sequence similarity network, we performed NCBI BLAST search to obtain the 1000 most closely related sequences to each of these previously characterized AAOs (fungal AAOs: *Komagataella phaffii* and *Aspergillus terreus*; and bacterial AAOs: *Rhodococcus* sp. strain RHA1 and *Pseudarthrobacter chlorophenolicus*) as measured by BLASTP. After deleting duplicate sequences, we then submitted to the Enzyme Function Initiative-Enzyme Similarity Tool^66,67^ (EFI-EST) to generate a sequence similarity network (SSN). To finalize the SSN, sequences exhibiting greater than 95% similarity were grouped into single nodes, resulting in 2270 unique nodes. The SSN was visualized and labeled in Cytoscape^68^ using the yFiles Organic Layout.

### Cloning and expression of pathway enzymes

Molecular cloning and vector propagation were performed in *E. coli* DH5α (NEB). The 52 AAOs were cloned by Golden Gate assembly in collaboration with the Joint Genome Institute. AAOs were cloned into a pZE vector harboring a C-terminal His6-tag for purification, kanamycin resistance gene for antibiotic selection, and ColE1 ori. For the transaminase (TA), the TA gene from *Chromobacterium violaceum* (CvTA) on a pACYC vector harboring an N-terminal His6-tag for purification, p15a ori, and chloramphenicol acetyltransferase gene for antibiotic selection from Gopal, et. al was used^54^. For the L-threonine transaldolase (TTA) and phenylalanine dehydratase (PSDH), the SUMO tagged PSDH gene from *Ralstonia pickettii* (RpPSDH) was expressed with either the SUMO tagged TTA genes from *Pseudomonas fluorescens* (ObiH) or from *Parachlamydiales bacterium* (PbTTA). The PSDH and TTA genes on a pColA vector with IPTG-inducible expression of both *Rp*PSDH with a hexahistidine-SUMO tag at the N-terminus and an TTA with a hexahistidine-SUMO tag at the N-terminus (either ObiH or *Pb*TTA), colA origin of replication and carbenicillin resistance from Anderson et al was used^56^. All primers were ordered from IDT. Plasmids were verified by Sanger sequencing.

### Aldehyde Stability Assays

For resting cell stability testing, cells of MG1655, RARE.Δ16^9^, ROAR.Δ12^28^ and ROAR.Δ22^27^ were prepared as described previously. The mass of cell pellets was then measured, and the pellets were resuspended in 200 mM HEPES, 7.0 and at a wet cell weight of 50 mg/mL. The resuspended resting cells were then aliquoted into 96-deep-well plates and supplemented with 5 mM aldehydes of interest (prepared in 100 mM stocks in DMSO) at a reaction volume of 400 µL. Resting cells were then incubated at 30 °C with shaking at 1000 RPM and an orbital radius of 3 mm. Samples were taken by pipetting 100 µL from the cultures, centrifuging in a round bottom plate, and collecting the extracellular broth. Compounds were quantified over a 24 h period using HPLC with samples collected at 4 h and 24 h.

### AAO resting cell assays

To assay AAOs in resting cells, AAO expression plasmids were transformed into ROAR.Δ12 and ROAR.Δ22. Resting cells were prepared as previously described. Overexpressed cells were then resuspended in buffer with 200 mM HEPES at pH 7.0. The resuspended plate and cells were then aliquoted into 96-deep-well plates at a wet cell weight of 50 mg/mL and supplemented with 5 mM substrate of interest (prepared in 100 mM stocks in DMSO) at a reaction volume of 400 µL. Resting cells were then incubated at 30 °C with shaking at 1000 RPM and an orbital radius of 3 mm. Samples were taken by pipetting 100 µL from the cultures, centrifuging in a round bottom plate, and collecting the extracellular broth. Compounds were quantified over a 24 h period using HPLC with samples collected at 4 h and 24 h. For whole cell reaction kinetics compounds were monitored over an hour with timepoints taken every 15 minutes. For scaleup experiments, vanillyl alcohol stocks were prepared at 800 mM to minimize DMSO percentage. All experiments were run with 5% DMSO and timepoints were taken at 4 and 20 h.

For AAO and CvTA coupled reactions, two separate cells were utilized to easily modulate enzymatic ratios. To prepare CvTA cells, pACYC-CvTA-AlaDH was transformed into ROAR.Δ12 and ROAR.Δ22. Resting cells were prepared as previously described. Cells were resuspended in buffers with 200 mM HEPES,100 mM L-alanine, 50 mM NH4Cl and 1 mM PLP at a pH of 7.5. Alcohol substrates were supplemented at 5 mM to a total cell concentration of 100 mg/mL wet cell weight of AAO and CvTA cells at ratios of 5:1, 1:1 and 1:5 (AAO cells: TA cells). Samples were taken by pipetting 100 µL from the cultures, centrifuging in a round bottom plate, and collecting the extracellular broth. Compounds were quantified over a 24 h period using HPLC with samples collected at 4 h and 24 h.

For AAO and TTA coupled reactions, two separate cells were utilized to easily modulate enzymatic ratios. To prepare TTA cells, pCola-TTA-PSDH plasmids were transformed into ROAR.Δ12 and ROAR.Δ22. Resting cells were prepared as previously described. Cells were resuspended in buffer with 200 mM HEPES, 10 mM MgCl2, 200 mM L-threonine, 50 mM L-glutamate and 1 mM PLP at a pH of 7.5. Substrates were supplemented at 5 mM to a total cell concentration of 100 mg/mL wet cell weight of AAO and TTA cells at ratios of 1:1 and 1:2 (AAO cells: TTA cells). Samples were taken by pipetting 100 µL from the cultures, centrifuging in a round bottom plate, and collecting the extracellular broth. Compounds were quantified over a 24 h period using HPLC with samples collected at 4 h and 24 h.

### Growing cell assays

For metabolically active AAO assays, pZE plasmids expressing each AAO variant were transformed into RARE.Δ16 cultures. Strains were then inoculated from a frozen stock and grown to confluence overnight in 5 mL of LB media with 50 µg/mL kanamycin. Confluent overnight cultures were then used to inoculate experimental cultures in a 96-deep-well plate initial starting OD_600_ of 0.05 in 300 µL in MOPS media with 50 µg/mL kanamycin at pH 7.5. At mid-exponential phase, we induced each culture then supplied 5 mM alcohol substrate (prepared in 100 mM stocks in 100% DMSO). Cultures were then incubated at 30 °C with shaking at 1000 RPM and an orbital radius of 3 mm. Samples were taken by pipetting 100 µL from the cultures, centrifuging in a different 96-deep-well plate and collecting the extracellular broth. Compounds were quantified over a 24 h period using HPLC with samples collected at 4 h and 24 h.

