## Supplementary Information for "Air-driven aldehyde synthesis in engineered bacteria via gene deletion and aryl-alcohol oxidase profiling"

### Supplementary Tables

**Supplementary Table 1**. Performance of two online predictors of protein solubility for our candidate AAOs

| sequence_id | activity on vanillyl alcohol | measured soluble expression (based on image analysis) | NetSol predicted solubility | SoluProt predicted solubility |
|---|---|---|---|---|
| OaAAO | 0.000 | 95 | 0.585 | 0.3431 |
| PsAAO | 0.000 | 95 | 0.595 | 0.2425 |
| ScAAO | 0.000 | 90 | 0.710 | 0.4501 |
| RsAAO | 0.000 | 95 | 0.557 | 0.3476 |
| BaAAO | 0.000 | 100 | 0.589 | 0.4106 |
| MlAAO | 0.000 | 100 | 0.585 | 0.2421 |
| CcAAO | 0.000 | 95 | 0.654 | 0.5233 |
| LeAAO | 0.000 | 95 | 0.547 | 0.258 |
| RbAAO | 5.565 | 95 | 0.572 | 0.5186 |
| VjAAO | 0.104 | 100 | 0.555 | 0.2475 |
| RhsAAO | 0.000 | 95 | 0.457 | 0.3986 |
| RhobAAO | 0.000 | 75 | 0.574 | 0.3101 |
| RiAAO | 0.000 | 95 | 0.560 | 0.3162 |
| RuhAAO | 0.030 | 95 | 0.558 | 0.3417 |
| HsAAO | 0.000 | 100 | 0.520 | 0.286 |
| JtAAO | 0.000 | 95 | 0.591 | 0.3586 |
| Bb1AAO | 0.000 | 95 | 0.491 | 0.4339 |
| OgAAO | 0.000 | 95 | 0.541 | 0.1817 |
| VsAAO | 0.000 | 95 | 0.513 | 0.5233 |
| Bb2AAO | 0.000 | 95 | 0.589 | 0.2078 |
| LaAAO | 0.000 | 95 | 0.503 | 0.4486 |
| VbAAO | 0.000 | 95 | 0.493 | 0.2901 |
| NgAAO | 0.060 | 0 | 0.543 | 0.648 |
| AsAAO | 0.040 | 0 | 0.522 | 0.3949 |
| AaAAO | 0.032 | 0 | 0.537 | 0.6058 |
| NsAAO | 0.030 | 5 | 0.599 | 0.7517 |
| EsAAO | 0.048 | 10 | 0.524 | 0.5491 |
| BsAAO | 0.053 | 95 | 0.614 | 0.7808 |
| CzAAO | 0.024 | 70 | 0.569 | 0.5334 |
| PbAAO | 0.017 | 20 | 0.604 | 0.6111 |
| GbAAO | 0.039 | 0 | 0.552 | 0.4881 |
| MrAAO | 0.024 | 0 | 0.627 | 0.6606 |
| RuAAO | 0.728 | 0 | 0.530 | 0.6076 |
| GhAAO | 0.016 | 10 | 0.550 | 0.7231 |
| MkAAO | 0.115 | 0 | 0.623 | 0.7137 |
| EwAAO | 0.015 | 0 | 0.552 | 0.321 |
| StAAO | 0.018 | 0 | 0.603 | 0.2016 |
| RhAAO | 0.020 | 0 | 0.638 | 0.3321 |

| | | | | |
|---|---|---|---|---|
| CmAAO | 0.007 | 100 | 0.588 | 0.6487 |
| LmAAO | 0.012 | 10 | 0.547 | 0.5235 |
| RoAAO | 0.021 | 0 | 0.568 | 0.3206 |
| AbAAO | 0.015 | 0 | 0.606 | 0.3215 |
| MaAAO | 4.263 | 80 | 0.564 | 0.3172 |
| PcAAO | 0.009 | 100 | 0.619 | 0.2885 |
| CoAAO | 0.944 | 80 | 0.550 | 0.3127 |
| VmAAO | 0.012 | 0 | 0.540 | 0.4954 |
| NnAAO | 0.019 | 0 | 0.590 | 0.3236 |
| PclAAO | 0.043 | 100 | 0.561 | 0.6467 |
| PrAAO | 0.037 | 75 | 0.550 | 0.3426 |
| SpAAO | 0.279 | 75 | 0.485 | 0.2704 |
| BuAAO | 6.008 | 80 | 0.505 | 0.2585 |
| AtAAO | 0.000 | 0 | 0.464 | 0.521 |
| **$R^2$** | | | **-0.034** | **-0.358** |

**Supplementary Table 2.** Complete alignment results for the top-16 positions that exhibited greatest Jensen-Shannon divergence between "in-cluster, high expression" and "other" groups.

| Position in Alignment | 61 | 90 | 95 | 105 | 144 | 250 | 265 | 399 | 451 | 505 | 509 | 600 | 608 | 630 | 636 | 726 | |
|---|---|---|---|---|---|---|---|---|---|---|---|---|---|---|---|---|---|
| Position in VjAAO | 33 | 52 | 57 | 64 | 103 | 191 | 201 | 299 | 342 | 361 | 365 | 435 | 443 | 461 | 467 | 510 | |
| **consensus** | **C** | **V** | **P** | **A** | **W** | **Q** | **K** | **F** | **A** | **L** | **I** | **E** | **R** | **D** | **D** | **T** | expression |
| OaAAO | C | V | P | A | A | H | K | I | A | L | R | N | R | D | D | T | 95 |
| PsAAO | C | V | P | A | L | Q | K | Y | A | L | E | E | G | D | D | T | 95 |
| CoAAO | C | M | P | A | H | Q | K | Y | A | L | P | E | R | D | D | T | 80 |
| RsAAO | C | V | P | G | W | H | K | Y | A | L | L | E | R | D | D | T | 95 |
| BaAAO | C | Y | P | A | W | Q | N | Y | G | L | I | E | R | D | D | T | 100 |
| RhsAAO | C | A | P | A | W | Q | K | Y | T | L | I | E | R | D | D | T | 95 |
| MaAAO | C | V | P | A | W | H | K | H | A | L | L | E | R | D | D | T | 80 |
| Bb2AAO | C | V | P | A | H | Q | K | Y | A | L | V | E | R | D | D | T | 95 |
| VjAAO | C | V | P | A | W | Q | K | H | A | L | V | E | R | D | D | T | 100 |
| ScAAO | C | T | P | A | W | Q | K | Y | A | L | I | H | R | E | D | T | 90 |
| CcAAO | C | L | P | A | W | I | K | Y | A | L | P | E | R | D | D | K | 95 |
| VsAAO | C | V | P | A | Y | Q | K | L | A | F | V | E | R | D | D | T | 95 |
| VmAAO | C | C | P | G | Y | Q | K | L | A | F | V | Q | R | D | D | T | 0 |
| MlAAO | C | V | P | A | S | Q | A | F | A | L | I | E | R | H | D | T | 100 |
| VbAAO | C | T | P | A | D | Q | A | F | A | L | I | R | R | H | D | T | 95 |
| LeAAO | C | T | P | A | A | Q | A | F | A | L | I | M | R | H | D | T | 95 |
| RiAAO | C | V | P | A | Q | Q | A | F | A | L | I | E | R | H | D | T | 95 |
| RuhAAO | C | A | P | A | Q | Q | S | F | A | L | I | E | R | H | D | K | 95 |
| RbAAO | C | V | P | A | T | Q | A | F | A | L | I | M | R | H | D | T | 95 |
| RhobAAO | C | V | P | A | Q | Q | A | F | A | T | I | E | R | V | D | L | 75 |
| JtAAO | C | V | P | A | Q | Q | A | F | A | T | I | T | I | H | D | T | 95 |
| Bb1AAO | C | V | P | A | S | Q | L | F | A | L | I | E | K | H | D | T | 95 |
| OgAAO | C | V | P | A | S | Q | A | F | A | L | I | Q | C | H | D | T | 95 |
| HsAAO | C | V | P | A | Q | Q | A | F | A | L | I | M | C | M | D | T | 100 |
| LaAAO | C | M | P | A | Q | Q | A | F | A | L | I | M | R | K | D | T | 95 |
| BuAAO | C | T | P | A | Q | Q | A | F | A | L | I | L | R | L | D | T | 80 |
| RhAAO | A | A | P | A | S | Q | R | F | A | L | V | Q | V | V | D | T | 0 |
| PrAAO | C | T | P | A | W | Q | R | F | A | L | V | Q | I | L | D | T | 75 |
| LmAAO | C | F | S | G | L | Q | R | T | S | L | T | A | R | F | D | Q | 10 |
| StAAO | C | F | P | Q | F | Q | R | Y | A | F | P | A | A | L | D | R | 0 |
| NnAAO | L | V | K | G | T | I | R | V | L | L | I | A | R | W | E | L | 0 |
| SpAAO | V | Y | K | L | S | I | V | V | A | F | P | A | I | Y | T | R | 75 |
| EwAAO | T | F | K | W | E | T | D | C | C | Y | A | A | V | F | Q | F | 0 |
| RoAAO | L | Y | G | G | A | N | N | V | V | F | A | A | L | F | R | K | 0 |
| AbAAO | L | F | S | G | E | T | V | L | V | F | P | A | V | F | R | R | 0 |
| PcAAO | L | F | A | G | G | T | V | L | V | F | P | A | I | F | R | R | 100 |
| NsAAO | A | R | L | D | E | R | V | A | W | M | S | A | A | Y | N | E | 5 |
| MrAAO | A | R | L | D | E | R | V | G | W | F | S | A | A | Y | N | E | 0 |
| MkAAO | A | R | L | D | E | R | V | G | W | M | S | S | A | Y | N | E | 0 |

| | | | | | | | | | | | | | | | | | |
|---|---|---|---|---|---|---|---|---|---|---|---|---|---|---|---|---|---|
| PclAAO | A | R | L | D | E | R | V | G | W | M | S | A | T | Y | N | E | 100 |
| CmAAO | A | R | L | D | E | R | V | G | W | F | S | A | A | Y | N | E | 100 |
| BsAAO | G | R | L | D | E | R | V | G | W | M | S | S | A | Y | N | E | 95 |
| GhAAO | A | R | L | D | E | S | V | G | W | M | S | K | A | Y | N | E | 10 |
| AtAAO | G | F | T | L | Q | V | T | S | S | Y | Q | S | V | A | M | G | 0 |
| NgAAO | P | A | N | - | R | R | - | A | W | Y | K | R | D | W | H | Q | 0 |
| AaAAO | P | A | N | - | R | R | - | A | W | Y | K | R | D | W | H | Q | 0 |
| EsAAO | P | A | G | - | R | R | - | I | W | Y | K | R | D | W | H | Q | 10 |
| CzAAO | K | A | N | - | K | N | - | L | W | Y | K | R | D | W | H | Q | 70 |
| PbAAO | K | A | T | - | K | N | - | L | W | Y | K | R | D | W | H | K | 20 |
| AsAAO | A | A | S | - | K | R | - | A | W | Y | K | - | E | W | H | Q | 0 |
| RuAAO | E | A | N | - | K | R | - | V | W | Y | K | R | Q | W | H | R | 0 |
| GbAAO | Q | A | D | - | R | E | - | V | W | Y | S | R | D | W | H | M | 0 |

**Supplementary Table 3.** Strains and plasmids used in this study.

| Name | Relevant genotype | Source |
|---|---|---|
| | ***E. coli* strains** | |
| DH5α | F– Φ80*lac*ZΔM15 Δ(*lac*ZYA-argF) U169 *rec*A1 *end*A1 *hsd*R17 (rK–, mK+) *pho*A *sup*E44 λ– *thi*-1 *gyr*A96 *rel*A1 | NEB |
| MG1655 | F- λ- *ilv*G- *rfb-50 rph-1* | ATCC 700926 |
| MG1655 (DE3) | F- λ- *ilv*G- *rfb-50 rph-1* (λ DE3)<br>λ DE3 = λ sBamHIo ΔEcoRI-B int::(*lac*I::PlacUV5::T7 gene1) i21 Δ*nin*5 | Previous study[1] |
| RARE.Δ6 | MG1655(DE3) Δ*dkgB* Δ*yeaE* Δ*(yqhC-dkgA)* Δ*yahK* Δ*yjgB* | Previous study [1] |
| RARE.Δ16 | RARE.Δ6 Δ*adhP,* Δ*fucO,* Δ*eutG,* Δ*yiaY,* Δ*adhE,* Δ*eutE,* Δ*gldA,* Δ*gpr,* Δ*ybbO,* Δ*yghA* | Previous study[2] |
| ROAR.Δ12 | RARE.Δ6 Δ*yiaY,* Δ*gpr,* Δ*ybbO,* Δ*yghA* | Previous study[3] |
| ROAR.Δ22 | RARE.Δ16, ROAR.Δ12 | Previous study |
| RMD001-RMD052 | ROAR.Δ12 harboring pZE-AO (each AO variant tested) | This study |
| RMD053 | ROAR.Δ22 harboring pZE-BuAAO | This study |
| RMD054 | ROAR.Δ22 harboring pZE-MaAAO | This study |
| RMD055 | ROAR.Δ22 harboring pZE-CoAAO | This study |
| RMD056 | ROAR.Δ22 harboring pZE-RbAAO | This study |
| RMD057 | ROAR.Δ22 harboring pACYC-CvTA-AlaDH | This study |
| RMD058 | ROAR.Δ22 harboring pColA-PSDH-ObiH | This study |
| RMD059 | ROAR.Δ22 harboring pColA-PSDH-PbTTA | This study |
| | **Plasmids** | |
| pZE-AAO (each variant) | ColE1 ori, $Kan^R$, TetR, Tet promoter with a codon optimized AAO cloned with help from the Joint Genome Institute | This study |
| pACYC-CvTA-AlaDH (Addgene ID: 206430) | P15a ori, $Cm^R$, LacI, Lac promoter with the codon-optimized gene for ω-transaminase from *Chromobacterium violaceum* and a separate Lac promoter with the codon-optimized gene for alanine dehydrogenase from *Bacillus subtilis* | Previous study |
| pColA-PSDH-ObiH | ColA ori, $carb^R$, LacI, Lac promoter with the codon-optimized gene for phenylalanine dehydratase from Ralstonia pickettii and a separate Lac promoter with the codon-optimized gene for L-threonine transaldolase from Pseudomonas fluorescens | Previous study |
| pColA-PSDH-PbTTA | ColA ori, $carb^R$, LacI, Lac promoter with the codon-optimized gene for phenylalanine dehydratase from Ralstonia pickettii and a separate Lac promoter with the codon-optimized gene for L-threonine transaldolase from Parachlamydiales bacterium | Previous study |

**Supplementary Table 4.** Oligonucleotides used in this study

| Oligo Name | Sequence (5’ to 3’) |
| --- | --- |
| pZE-sequencing | taaaaataggcgtatcacgagg |

**Supplementary Table 5.** Sequences of proteins expressed in this paper.

| Plasmid name (NCBI number) | DNA CDS | Protein Sequence |
|---|---|---|
| pZE-NgAAO (WP_103096198.1) | ATGCCCCAGGGAGTTGATGCGGCCACCTTTACAAAAGCGCTGGATGAACTGGCCGTGATTGTTGGGAAAGAATGGGTTTTCGTTGATGAACTGCCGCTATCCGCCTATCGCGACGCCTATAGCCCGCTGGCGGATGGCGAAATGTTACCAAGCGCCGCTGTTGCACCAGCAAACATGGAACAGATTCAGCAGGCATTAAAAGTCTTTAACGCATATAAACTGCCCATTTGGACTTTTGGGAACGGGCGGAACTTTGCCTACGGCGGACCAGCTCCGAGACAATCCGGCTATGTTATGTTTGACCTGAAAAGGATGAATAGGATTCTCGAAGTCAACGAGAAATATGGGTATGCATTAGTGGAGCCCGGAGTCTCGTACTTCCAACTTCATCGTCATCTTCGAAAGATTGGCTCCAAACTTTGGGTTGATCCTGCCGCGCCTGGGTGGGGAGGAGTGATGGGTAACGCACTAGAACATGGTGCCGGCTACACACCGTATGGCGATCATTTCGTGATGCAATGCGGTATGGAGGTTGTATTAGCCGATGGAAAAGTGGTCCGGACGGGCCAGGGAGCTATCGAGGGCTCGCACCATTGGCAATCAACCAAGCACGCTGCCGGTCCGCATTTCGACGGAATGTTCACTCAGTCGAATTTTGGCATAGTGACCAAAATGGGGATATGGCTCATGCCAGAGCCACCAGGGTACAAACCGTTCATGATAACCTATGAACGAGAGGAAGACCTGGCGGCGATTTTTGATGCTGTTTTGCCGCTGAAAATAAATCAAGTCATTCCCAATGCGGCTGTTGCAGTTGATCTCCTGTGGGAAGTTAGTGCCAAAACCACACGCCGACATTATTTTGATGGCAAAGGTCCCATACCTGATAGTATTCGGAAAAAAATAGCTAGCGACCACGGTCTGGGCATGTGGAATTTCTATGCAGCATTATATGGCCCGCCGCCTATTATTGAAAACAACTGGAAACTCGTAGAGGAAGCGATGATGTCTATACCCGGTGCCAAACTCCATCTCGATAGGGAAAACGATCCTGCATGGGATTACCGGGTGCAACTAATGCGTGGCGAACCGAATATGACGGAATTTAGTATCATGAACTGGATAGGAGGTGGCGGGCATATCAATTTTTCCCCGATCTCAGCCCCCGATGGTAAAGAGGCCCTGAGTCAGTATAATCTGATTAAGCAGCGGTGTCACGATTTCGGTTTTGATTACATTGGGGAATTTTTGGTTGGTTGGAGAGATATGCATCATATTCTGATGATAATGTATGATCGCGCGGACGATGGTATGCGTAAGTCAGCATATGATTTGTTCGGCAAACTTGTGGATGAAGCAGCGGGGGCAGGGTTTGGAGAATACCGTACTCACCTGGCTTTTATGGATCAGATCGCCAAGACATATAAGCATAATGACGGAGCACTTTGGGATTTGCACCATCGTCTTAAAGATGTTCTGGATCCGAACGGTATACTGTCGCCTGGCAAGCAGGGTATATGGCCCCAGGCGATGCGCAATCAGGCA | MPQGVDAATFTKALDELAVIVGKEWVFVDELPLSAYRDAYSPLADGEMLPSAAVAPANMEQIQQALKVFNAYKLPIWTFGNGRNFAYGGPAPRQSGYVMFDLKRMNRILEVNEKYGYALVEPGVSYFQLHRHLRKIGSKLWVDPAAPGWGGVMGNALEHGAGYTPYGDHFVMQCGMEVVLADGKVVRTGQGAIEGSHHWQSTKHAAGPHFDGMFTQSNFGIVTKMGIWLMPEPPGYKPFMITYEREEDLAAIFDAVLPLKINQVIPNAAVAVDLLWEVSAKTTRRHYFDGKGPIPDSIRKKIASDHGLGMWNFYAALYGPPPIIENNWKLVEEAMMSIPGAKLHLDRENDPAWDYRVQLMRGEPNMTEFSIMNWIGGGGHINFSPISAPDGKEALSQYNLIKQRCHDFGFDYIGEFLVGWRDMHHILMIMYDRADDGMRKSAYDLFGKLVDEAAGAGFGEYRTHLAFMDQIAKTYKHNDGALWDLHHRLKDVLDPNGILSPGKQGIWPQAMRNQA |
| pZE-AsAAO (WP_126536808.1) | ATGCCCGCGACCCTCCCGCCGGGCTTATCCGCAGCAGCATTCAGTCAAGCCCTTGCGGGGTTTCGCCAGGCCGTTGGCAAGACATATGTTTATGCAGACGAGACTGCTCTTTCTTCGTACTTGGATCCTTATAGCACTACGGAAGATGCCGCTCATACACCGGCGGCAGCAGTCGCTCCACACTCAGTTGAGGAGATCCAAGCAGTATTAAAGGTAGCACGAGAATACGGTATTCCGTTGTGGCCCGTTTCCACTGGTAAAAATTACGCTTATGGTGGTCCAGCACCGCGAAAGTCGGGGTATGTGGTACTGGATTTAGCGCGTATGAATCGTATTATTGAGGTTAACGCGCGCGACGGATATGCTGTCGTGGAACCTGGTGTGAGCTATTTTGATCTTTATCGATATCTGAAAGAGCACGATATTCCTCTGTGGATCGACTGTGCAGCACCGGGTTGGGGATCTGTGTTGGGGAACCTATTAGATCACGGTGCAGGCTATACCCCGTATGGGGAGCACTTAACTATGCAGTGTGGCATGCAGGTTGTACTCGCCGACGGCACAGTTGTTGATACGGCAACGGGAGCGTTGCCGGGCGCACGCGCTTCACAACTTTACAAATGGGGTGCCGGGCCGTGGATCGACGGACTCTTCACGCAGTCTGGTCTAGGAATTGTAACACGCTTAGGCGTTTGGCTCATGCCGGAGCCGCCTGGTTATCGGCCATTTATGGTGACGTTTCCAGATGAAGATGCGCTGCATGATCTGACAGAAGCGATCCGCCCGTTGAAATTAAACATGGTGATCCCCAACGGGGCTACATCTGTAGAGTTACTGTGGGAAGCAGCCACACGGGTAACTAAAGCGCAGTATTATGGAGGTAAAGGGCCCTTACCCCCGTCCGTTCGACGTAAGCTAATGGCTGATCTGGATATTGGTGCTTGGAACTTTTATGCAGCTCTTTATGGTCCTCCTCCGATGATTGAAAGATCCTGGAGTGTCGTACGGGACGCACTGGGCAAAGTTAAAGGAGCCAGATTCTACACCGACCGCCCGGGTGATACTGCATGGGAATACAGGGCCAAGTTAATGCGCGGCATTCCGAACATGACAGAATACTCCCTGATGAATTGGATTGGATCAGGCGCGCATATTGATTTTTCACCCATGTCTGCTCCAACTGGCGACGATGCTGTTCGCCTGTACCATCTTATTAAAGATAAGGTGGAAGCTCGAGGGTTTGATTATATTGGCGAGTTTCTGATTGGGTGGCGTGACCAACACCATATTTTCATGCTGATATTTGATCGTACAGATCCACAAGAGCGCGAACGAGCCCATCAGGTATTTGGGGAGATCGTAGCAGATGCAGCAGCCCAAGGCTTTGGGGAATATCGGACACACCTGGATTTTATGGATCAGATTGCCGCAACATACGACTGGAATGACCATGCTCTGCGCCGACTCAACGAGAGGGTTCGCGACGCATTAGATCCCGGCGGCATTATGGCACCGGGTAAGATGGGCGTGTGGCCCCGCCGTATGCGCGCT | MPATLPPGLSAAAFSQALAGFRQAVGKTYVYADETALSSYLDPYSTTEDAAHTPAAAVAPHSVEEIQAVLKVAREYGIPLWPVSTGKNYAYGGPAPRKSGYVVLDLARMNRIIEVNARDGYAVVEPGVSYFDLYRYLKEHDIPLWIDCAAPGWGSVLGNLLDHGAGYTPYGEHLTMQCGMQVVLADGTVVDTATGALPGARASQLYKWGAGPWIDGLFTQSGLGIVTRLGVWLMPEPPGYRPFMVTFPDEDALHDLTEAIRPLKLNMVIPNGATSVELLWEAATRVTKAQYYGGKGPLPPSVRRKLMADLDIGAWNFYAALYGPPPMIERSWSVVRDALGKVKGARFYTDRPGDTAWEYRAKLMRGIPNMTEYSLMNWIGSGAHIDFSPMSAPTGDDAVRLYHLIKDKVEARGFDYIGEFLIGWRDQHHIFMLIFDRTDPQERERAHQVFGEIVADAAAQGFGEYRTHLDFMDQIAATYDWNDHALRRLNERVRDALDPGGIMAPGKMGVWPRRMRA |
| pZE-AaAAO (WP_048886471.1) | ATGAGCAATCGGTTACCTCAAGGCATCGATGCGCAGACGTTCCAAGCAGCTTTGGAAGAACTGGGTGCTATAGTCGGACGAGACTGGGTGTTTGTCGATGAACTTCCGCTCAGCACTTATAGAGATGCTTATTCTCCCCTCGCGGATGGTGAACT | MSNRLPQGIDAQTFQAALEELGAIVGRDWVFVDELPLSTYRDAYSPLADGELLPS |

| | | |
|---|---|---|
| | GCTACCGTCTGCGGCGATCGCGCCTGCCAATGTAGATGAGATCCAGCGCGCC<br>CTGCGGGTTTTAACTCATACAAACTTCCTATTTGGACATTTGGTAATGGCCG<br>TAACTTTGCGTATGGTGGTCCTGCACCACGGCAGAGTGGATACGTTATGTTC<br>GACTTGAAACGGATGAATAGGATCCTGGAAGTTAACGAAAAATATGGATAT<br>GCCTTGGTAGAGCCGGGTGTAACCTATTATCAGCTCGATAGATATTTACGGC<br>AGACAGGTTCAAAGCTCTGGATTGATCCAGCAGCTCCAGGCTGGGGTGGTG<br>TCATGGGGAACGCCCTCGAACATGGTGCAGGATATACCCCGTACGGTGATC<br>ATTTCGTGATGCAATGTGGCATGGAAGTTGTGCTCGCAGACGGTGAAATAGT<br>TCGGACCGGTCAAGGTGCTATAGAGGGCTCACAGCACTGGCAGGCAACTAA<br>ACACGCGGCAGGCCCTCATTATGACGGCATGTTTACGCAAAGCAACTTTGGC<br>GTAGTTACTAAAATGGGTATGTGGCTGATGCCAGAACCACCTGGTTATAAAC<br>CTTTTATGATAACCTACCAGAGAGAAGAAGATCTCGCTGCCATCTTTGACGC<br>GGTACGGCCACTGAAAGTCAATCAGGTAATACCGAATGCGGCAGTCGCAGT<br>AGATCTTTTATGGGAAGTCAGTGCAAAGACCACGCGTCGGCATTACTTTGAT<br>GGCAAGGGGCCGTTGCCTGACAGCATCAGAGCGAAAATTGCCGCAGATCAC<br>GATCTCGGAGCATGGAATTTTTATGCAGCGATGTACGGTCCACCGCCAATCA<br>TTGAAAACAATTTTAAGCTAGTGCAGGATGCGCTCTTGTCGATTCCGGGTGC<br>AAAGTTGTATCTGGATCGACAAAATGACCCGGCATGGGATTATAGAGTTAG<br>ATTGATGCGCGGAGAACCAAATATGACGGAGTTCAGCATTATGAACTGGAT<br>TGGTGGGGGCGGCCATATTAATTTTAGTCCGATATCTGCTCCAGATGGCAAC<br>GAGGCATTGCTGCAGTACAATATGATTAAAGAGCGTTGCCACGACTTTGGTT<br>TCGATTACATTGGTGAGTTTTTGGTGGGATGGCGCGATATGCATCACATCCT<br>CATGATCATGTATGATCGTGCCGATGAAAAAATGCGTAAAGATGCGTACGA<br>CCTGTTTGGGCGTCTGGTCGACGAGGCCGCTGATGCGGGATTTGGAGAATAT<br>CGGACACACCTCGCTTTCATGGACCAGATTGCAGGCACTTATAAATACAATG<br>ATGGTGCCCTCTGGGACTTACATCATCGTCTAAAGGACGTCCTGGATCCGAA<br>TGGGATTCTCTCACCTGGCAAACAGGGCATCTGGCCGCAAAGTATGCGCAGT | AAIAPANVDEIQRALRVF<br>NSYKLPIWTFGNGRNFAY<br>GGPAPRQSGYVMFDLKR<br>MNRILEVNEKYGYALVEP<br>GVTYYQLDRYLRQTGSKL<br>WIDPAAPGWGGVMGNAL<br>EHGAGYTPYGDHFVMQC<br>GMEVVLADGEIVRTGQG<br>AIEGSQHWQATKHAAGP<br>HYDGMFTQSNFGVVTKM<br>GMWLMPEPPGYKPFMITY<br>QREEDLAAIFDAVRPLKV<br>NQVIPNAAVAVDLLWEVS<br>AKTTRRHYFDGKGPLPDS<br>IRAKIAADHDLGAWNFYA<br>AMYGPPPIIENNFKLVQD<br>ALLSIPGAKLYLDRQNDP<br>AWDYRVRLMRGEPNMTE<br>FSIMNWIGGGGHINFSPIS<br>APDGNEALLQYNMIKERC<br>HDFGFDYIGEFLVGWRD<br>MHHILMIMYDRADEKMR<br>KDAYDLFGRLVDEAADA<br>GFGEYRTHLAFMDQIAGT<br>YKYNDGALWDLHHRLKD<br>VLDPNGILSPGKQGIWPQS<br>MRS |
| pZE-NsAAO<br>(PRZ12759.1) | ATGGAAAACACGACATTCGATTATGTCGTTATCGGAGGCGGCTCTGGTGGTG<br>CAGCAGTAGCAGCAAGACTAAGCGAAGATCCTAACGTGACGGTAGCACTGT<br>TAGAAGCAGGTCCTACAGACGCGGATAAACCCGAAATTCTGCAGCTCAACC<br>GGTGGATGGAACTGCTGGAAAGCGGTTTCGACTGGGATTATCCCATAGAGG<br>AACAGGAAAATGGTAACTCCTTTATGCGCCACGCTCGAGCTAAAGTCCTAG<br>GCGGCTGCTCTTCTCACAATTCATGTATTGCTTTTTGGGCTCTGAGAGAAGA<br>CATGGATGAATGGGAAGCCAAAGGTGCCACGGGATGGAACGCAGAATCTAC<br>GTATCCTTTGTTTCAACGTTTAGAAACGAATGATGCTCCTGGCGACCATCAT<br>GGTCGGAGCGGTCCCGTTAACTTGATGACGATACCGCCCAAGGATCCGTGC<br>GGGGTGGCCCTTCTTGATGCCTGCGAAGAGCAAGGGATTCCGAGAGTTGCA<br>TTTAACACTGGAGAGACGGTGCTGAAAGGAGCCAACTTCTTTCAGGTGAATC<br>GTAAAGAAGACGGTACACGTTCGAGTTCGTCGGTATCCTATCTGCATCCAAT<br>TCTGCAGCGGCCCAATTTGTCAATACTCACTGATACATGGGTACGGGAAATT<br>ATCTTCGAAGGTGACCGTGCTGTAGGCGTAGATGCGGTTAATAATCCATTCG<br>GACGTGCACATAGGATAAGTGCCACCAAAGAAATTGTGTTAAGTGCGGGTG<br>CGATCGATTCCCCAAAACTACTGATGCTAAGCGGTATTGGTCCAGCACAACA<br>TTTAGCGGAATTCGGTATCGAAGCACGGGTGGATTCCCCGGGCGTGGGTGA<br>ACATCTGCAAGACCATCCAGAAGCTGTAATCGCATGGGAGGCAAAGCAACC<br>CATGGTGAAAGACTCCTATCAGTGGTGGGAGATCGGTATATTTTCGGCGACC<br>CGTGAGGGCTTGGATCGCCCGGATCTGATGATGCATTATGGGTCTGTCCCGT<br>TTGACATGCACACACTCAGGCAAGGTTATCCGACCGCTGATAATACTTTCTG<br>TCTCACCCCCAATGTAACCCACGCTAAAAGCCGCGGCACGGTTAGGCTGCGC<br>TCAAAAGATTATCGTGACAAACCGAAGGTTGACCCGAGGTACTTTACTGATC<br>CAGAGGGTCATGATATGCGGGTAGCAGTTGAAGGCATTAAACTTGCCAGAA<br>AAATCGCAGCACAGCCGGCCATGGCCGACTGGGCTGGTCAGGAGCTGTTTC<br>CCGGTGCGGATACCCAGACTGATGAGGAGCTCGCAGATTACGTAAGCCGAA<br>CTCATAATACTGTCTATCATCCAGCGGGCTCTGTTCGGATGGGCGCTGATGA<br>TGACGCCATGAGCCCAGTGGATAGCCAGTTAAGAGTAAAGGGCGTGCATGG<br>CCTGAGAGTAGCGGATGCAAGTGTGATGCCGGAGCTGACAACGGTTAATCC<br>TAATATTACCACAATGATGATTGGTGAACGCTGTGCCGATCTCATTAAAGCC<br>GCGAACGCA | MENTTFDYVVIGGGSGGA<br>AVAARLSEDPNVTVALLE<br>AGPTDADKPEILQLNRWM<br>ELLESGFDWDYPIEEQEN<br>GNSFMRHARAKVLGGCS<br>SHNSCIAFWALREDMDE<br>WEAKGATGWNAESTYPL<br>FQRLETNDAPGDHHGRSG<br>PVNLMTIPPKDPCGVALL<br>DACEEQGIPRVAFNTGET<br>VLKGANFFQVNRKEDGT<br>RSSSSVSYLHPILQRPNLSI<br>LTDTWVREIIFEGDRAVG<br>VDAVNNPFGRAHRISATK<br>EIVLSAGAIDSPKLLMLSG<br>IGPAQHLAEFGIEARVDSP<br>GVGEHLQDHPEAVIAWE<br>AKQPMVKDSYQWWEIGI<br>FSATREGLDRPDLMMHY<br>GSVPFDMHTLRQGYPTAD<br>NTFCLTPNVTHAKSRGTV<br>RLRSKDYRDKPKVDPRYF<br>TDPEGHDMRVAVEGIKLA<br>RKIAAQPAMADWAGQEL<br>FPGADTQTDEELADYVSR<br>THNTVYHPAGSVRMGAD<br>DDAMSPVDSQLRVKGVH<br>GLRVADASVMPELTTVNP<br>NITTMMIGERCADLIKAA<br>NA |
| pZE-EsAAO<br>(WP_07267379<br>4.1) | ATGAGTTCAGTGGTTCCGCAAGGAGTGGCTGCAGCGGACTTTTCTAACGCCA<br>TTAAAGAATTTGCGGGCGTGGTGGGCAAGGACCGTGTTTATGTAGATGAGC<br>AGCCCCTTTCATCATATAGGGACGCTTACTCACCCCTGGCAGACGGTGAATT<br>CATTCCGAGCGCAGCCGTTGCACCCGATGGCGTCGAGCAAATTCAGAAAAT<br>TCTGAAGATAGCGAACGCCTACAAGATTCCTCTGTGGACGATTGGAACAGG<br>ACGAAATTTCGCTTACGGTGGTCCTGCACCACGTAAATCTGGTTATGTAATG<br>CTAGATCTGCGTCGTATGAACCGGATAATTGAGGTTAATGAAAAATATGGTT<br>ACGCCTTAGTTGAACCGGGAGTTTCCTATATGCAGCTCTATCGGCACCTGCG<br>TGCTATTGGTTCAAAGTTATGGATTGACCCCGCCGCCCCCGCATGGGGGGGA<br>GTGATGGGTAATGCGCTGGAACATGGTGCAGGGTATACGCCGTATGGTGAC<br>CATTTTATAATGCAGTGCGGTATGGAAGTGGTGCTGGCGGACGGTGAAGTG<br>GTACGGACTGGCCAGGGCGCCCTGGCAGGTTCCAAACACTGGCAGGTGACC | MSSVVPQGVAAADFSNAI<br>KEFAGVVGKDRVYVDEQ<br>PLSSYRDAYSPLADGEFIP<br>SAAVAPDGVEQIQKILKIA<br>NAYKIPLWTIGTGRNFAY<br>GGPAPRKSGYVMLDLRR<br>MNRIIEVNEKYGYALVEP<br>GVSYMQLYRHLRAIGSKL<br>WIDPAAPAWGGVMGNAL<br>EHGAGYTPYGDHFIMQCG<br>MEVVLADGEVVRTGQGA<br>LAGSKHWQVTKHAAGPQ |

| | | |
|---|---|---|
| | AAACATGCTGCAGGTCCCCAGTTTGATGGTATGTTCACGCAGTCAAACTTCG<br>GAGTGGTTACCAAAATGGGCATTTGGCTTATGCCTGAACCGCCCGGGTATAA<br>GCCTTTTATGATAACATACGAGCGTGAAGAGGATCTGGAAGCTATTTTTGAA<br>ATAACCCGTCCGCTGAAGGTGAATCAAATAATCCCTAATGCTGCAGTGGCCG<br>TCGATCTGCTCTGGGAAGCATCAGCTAAAGTTACCAGACGCCATTACTTTGA<br>TGGCAAAGGTCCTCTTCCGCAAAGCGCCCGAACGCGGATTGCTAGTGATTTA<br>AAGTTAGGTATGTGGAATTATTATGGAGCGCTTTATGGCCCGCCGCCCATAA<br>TCGAAAACAACTGGAAAATAGTTGAAGAAGCGCTTATGAGCATACCTGGTG<br>CAAAACTGTACTTGGATAGAGAGAATGATCCTGCGTGGGATTATCGTGTTCG<br>CTTGATGAGAGGTGAGCCGAATATGACCGAATTTTCTATAATGAACTGGATA<br>GGCGGGGGGGGCACGTGAATTTTAGTCCGATTTCAGCCCCTAGCGGTGCA<br>GAGGCATTAGCACAGTACAACATGATTAAACGTCGCTGCCATGAATACGGT<br>TTCGACTATATTGGTGAGTTTCTGGTGGGATGGCGGGATATGCATCATATCT<br>TAATGATTATGTATGACCGTGCAGATGAGGCAATGCGTAACCGTGCGTATGA<br>TCTATTTGGGCTGTTAGTCGATGAAGCGGCAGAAGTGGGCTTTGGGGAGTAT<br>CGTACCCATCTCGCGTTTATGGACCAAATCGCTAAAACCTATAAACATAATG<br>ATGGTGCACTTTGGGATCTGCACCACAGGATTAAAGACGTGATGGATCCAA<br>ACGGTATACTGTCCCCGGGGAAGCAAGGTATTTGGCCTAAGGCCATGCGCG<br>AGACAAGC | FDGMFTQSNFGVVTKMGI<br>WLMPEPPGYKPFMITYER<br>EEDLEAIFEITRPLKVNQII<br>PNAAVAVDLLWEASAKV<br>TRRHYFDGKGPLPQSART<br>RIASDLKLGMWNYYGAL<br>YGPPPIIENNWKIVEEALM<br>SIPGAKLYLDRENDPAWD<br>YRVRLMRGEPNMTEFSIM<br>NWIGGGGHVNFSPISAPSG<br>AEALAQYNMIKRRCHEY<br>GFDYIGEFLVGWRDMHHI<br>LMIMYDRADEAMRNRAY<br>DLFGLLVDEAAEVGFGEY<br>RTHLAFMDQIAKTYKHN<br>DGALWDLHHRIKDVMDP<br>NGILSPGKQGIWPKAMRE<br>TS |
| pZE-BsAAO (WP_09201288 8.1) | ATGAAAACTACGCACAGCTTTGACTATGTCGTAGTGGGTGGGGGTTCAGCA<br>GGTGCCGCAGTCGCGGCACGCCTGAGCGAGGATCCAGAAGTCACGGTGGGC<br>CTGCTGGAAGCCGGTCCTACTGACACAGACAAAGATGTAGTTCTGCAACTCA<br>ATCGCTGGATGGAACTGTTGGAGTCTGGGTACGACTGGGACTACCCGATTGA<br>AGAACAAGAAAACGGGAACAGCCATATGCGCCATGCGCGCGCGAAAGTTCT<br>CGGAGGATGCAGCTCCCACAACAGTTGCATTGCCTTTTGGGCTCCTCGTGAG<br>GACCTGGATGAGTGGGAAAACAAACATGGTGCTGCAGGATGGGGCTCAAAA<br>GATACTTTCGGCCTGTATAAGAAATTGGAAAATAACGAATTACCAGGTGAC<br>CACCATGGGCATGATGGTCCGGTCAGACTGATGAACGTACCTCCTGTTGACC<br>CATGCGGCATAGCCCTGCTCGATGCGTGCGAGCAGGCGGGCATTCCTCGTGT<br>CCAGTTTAATGAGGGGCAGACGGTTATAAATGGGGCGAATTTTTTCCAAGTT<br>AACCGAAAAGATGACGGTACGCGCAGTAGTTCATCTGTAAGTTACCTGCAC<br>CCCGTGTTAGATCGCGAAAATTTAACTATCCTGACCGACACACAGGCCAAA<br>GAACTGGAATTCGATGAAAATGATAATTGTACAGCAGTATTGGTGGTCAAC<br>AACGCATTTGGGAAGACAAACCGTATAGAGGCTAAGAAAGAAGTAATTGTT<br>TCCTCAGGCGCAATAAATACTCCTCAGTTGCTCATGCTGAGCGGTATCGGTC<br>CAAAGGATCATTTAGCAGAGGTTGGCATTGAATCCCGTGTGGATAGCCCAG<br>GCGTCGGAGAACATTTAGGTGATCATCCGGAAGGGGTTATCTCCTGGGAAG<br>CGAAGAAACCTATGGTTGAGGACTCCACGCAGTGGTGGGAAATCGGGATTT<br>TTTCTCCAACAGAAGACGGTTTGGATAGACCCGATCTCATGATGCATTACGG<br>GTCGGTCCCGTTCGACATGCACACGCTGAGACAGGGCTATAAAACGAGTGA<br>GAATTCTTTCTGTTTGACACCCAACGTCACTCACGCAAAAAGTCGTGGCACC<br>GTTCGCCTTCGTTCGAAAGATTTTCGCGATAAACCGAAAGTAGATCCTCGCT<br>ATTTCACCGATCCGGAAGGCCATGACATGCGTGTAATGGTGGCTGGGATCA<br>AAAAGGCACGCGAAATCGTTAGCCAGGACGCCATGGCTGAATGGGCAGGCG<br>AAGAATTATTTCCGGGTAAAGAAGTTCAGACTGATGAACAGATTGCAGATT<br>ATATTACTCGAACCCATAATACGGTCTATCATCCTGTAGGTACTGTTCGTAT<br>GGGTGCGCACAATGATGAAATGAGCCCGTTAGATCCGCAATTGCGTGTGAA<br>AGGAGTCAATCGGTTACGCGTAGTTGATGCGTCTGCGATGCCGGAGATAAC<br>GACGGTTAACCCGAATATCACGGTAATGATGATGGGCGAAAAATGCGCTGA<br>AATGATCAAAAACGGTCAA | MKTTHSFDYVVVGGGSA<br>GAAVAARLSEDPEVTVGL<br>LEAGPTDTDKDVVLQLNR<br>WMELLESGYDWDYPIEE<br>QENGNSHMRHARAKVLG<br>GCSSHNSCIAFWAPREDL<br>DEWENKHGAAGWGSKD<br>TFGLYKKLENNELPGDHH<br>GHDGPVRLMNVPPVDPC<br>GIALLDACEQAGIPRVQFN<br>EGQTVINGANFFQVNRKD<br>DGTRSSSSVSYLHPVLDRE<br>NLTILTDTQAKELEFDEND<br>NCTAVLVVNNAFGKTNRI<br>EAKKEVIVSSGAINTPQLL<br>MLSGIGPKDHLAEVGIESR<br>VDSPGVGEHLGDHPEGVI<br>SWEAKKPMVEDSTQWWE<br>IGIFSPTEDGLDRPDLMMH<br>YGSVPFDMHTLRQGYKTS<br>ENSFCLTPNVTHAKSRGT<br>VRLRSKDFRDKPKVDPRY<br>FTDPEGHDMRVMVAGIK<br>KAREIVSQDAMAEWAGE<br>ELFPGKEVQTDEQIADYIT<br>RTHNTVYHPVGTVRMGA<br>HNDEMSPLDPQLRVKGV<br>NRLRVVDASAMPEITTVN<br>PNITVMMMGEKCAEMIK<br>NGQ |
| pZE-CzAAO (WP_02093298 3.1) | ATGACCAAGCCATTACCGCAAGGCATTACATTAGGACAGCTGAATGCTGCC<br>ATAGCCGAAATTGAAAAAGTAGTTGGGAAAGAATATGTATACTTAGATGAT<br>ATCAAGGAATTGCGATCGTATAGAGATCCGTATAACACCACAAATGACGCG<br>GATTTCGCACCGTCAGCCGCCGTTGCCCCACACAATGTAGAACAGATCCAGA<br>AAATCCTTGCCATCGTTAATAATTACAAAATTCCTGTTTGGACAATCAGTAC<br>AGGTAAAAACTTCGCTTACGGTGGTCCAGCTCCACGAAAACCGGGATATAT<br>CGTCCTAGACCTGAAATTAATGAATCGTATCCTGGAGGTAAACGAAAAACA<br>TGGTTATGCTATCGTCGAACCAGGGGTGAGTTATTACGATCTGTATAATTAC<br>CTGCAGGAAAGAGGTTCTAAATTATGGATCGACTGCGCAGCACCAGCGTGG<br>GGGGGGATAGTAGGTAATACGAGTGAACATGGCGCTGGTTACACACCGTAC<br>GGGGATCACCTGCTCATGCAGTGCGGCATGCAGGTCGTGCTGGCCGATGGA<br>ACCGTGGTGGAGACAGGCATGGATACGACCGCAAACTCGAAAACGGGTGGA<br>CTGTATAAATATGGTCAGGGGGCAGCAATTGATGGCTTATTTACTCAGTCCA<br>ACTTCGGGGTTATCACACGAATGGGAATCTGGCTTATGCCCGAGCCTCCGGG<br>CTATACTCCCTTTATGATTACTCTGGAGAACGAACATGATCTTGAAACCGTT<br>ACGGATCTGAGTCTTCCACTGAGAGTGAATCAAGTTATCCATAATGCTCCGA<br>TGACGGTAGATCTGCTGTGGGAAGCGGCCATGTCAGTCAGCCGAAGTCAGT<br>ACTATACGGGCAAGGGGCCAATGCCAGATTCTCACCGGAAGAAGATGGCTA<br>AAGATCTGGATCTGGGGATTTGGAATTACTACGGGGCTGTTTACGGCCCGGG<br>CCCGATGATGAAAAATAACATGGATGTGATCCGCGATACATTTGGAAGCAT | MTKPLPQGITLGQLNAAI<br>AEIEKVVGKEYVYLDDIK<br>ELRSYRDPYNTTNDADFA<br>PSAAVAPHNVEQIQKILAI<br>VNNYKIPVWTISTGKNFA<br>YGGPAPRKPGYIVLDLKL<br>MNRILEVNEKHGYAIVEP<br>GVSYYDLYNYLQERGSKL<br>WIDCAAPAWGGIVGNTSE<br>HGAGYTPYGDHLLMQCG<br>MQVVLADGTVVETGMDT<br>TANSKTGGLYKYGQGAAI<br>DGLFTQSNFGVITRMGIW<br>LMPEPPGYTPFMITLENEH<br>DLETVTDLSLPLRVNQVIH<br>NAPMTVDLLWEAAMSVS<br>RSQYYTGKGPMPDSHRK<br>KMAKDLDLGIWNYYGAV<br>YGPGPMMKNNMDVIRDT<br>FGSIPGAKFHYDDTRKGD |

| | | |
|---|---|---|
| | TCCTGGTGCGAAATTTCATTATGATGATACACGCAAAGGCGATCCCGGTTGG<br>GACTACCGGGTCAAACTGATGAAAGGCGTTCCCAACATGACCGAATTTAGC<br>CTGCTCAACTGGGTGGGCGGGGGCGGGCACATTAATTTTTCCCCGATCTCAG<br>CAGCGGATGGGAAGTCCAGCATGAACCAATACAAGATGATTGAAAAAAGA<br>GCGCATGAGTACGGTTTCGACTATATAGGGGAATTCCTGATCGGATGGCGTG<br>ATCAACACCATATCTTCATGTTGATGTTTAACCGCATGTCACCAGAAGAGAA<br>GAAACGTGCACATGACCTGTTTGCTATCCTTGTGGACGAAGCCGCGGCGTTA<br>GGTTACGGCGAATACAGAACCCATTTATCATTCATGGACCAGATTGCTGGCA<br>CATTTTCATGGAACGATAATGCTTTGTGGGACACTCACCATGCGCTTAAGGA<br>TGAGTTAGACCCTAATGGGATACTGTCCCCGGGCAAACAAGGAATTTGGCCT<br>AAGAATCTGCGCGGTGAAAGT | PGWDYRVKLMKGVPNM<br>TEFSLLNWVGGGGHINFS<br>PISAADGKSSMNQYKMIE<br>KRAHEYGFDYIGEFLIGW<br>RDQHHIFMLMFNRMSPEE<br>KKRAHDLFAILVDEAAAL<br>GYGEYRTHLSFMDQIAGT<br>FSWNDNALWDTHHALKD<br>ELDPNGILSPGKQGIWPKN<br>LRGES |
| pZE-PbAAO<br>(PCI69553.1) | ATGAATAAGCCTTTGCCACAAGGAATTTCACTCGAGCAATTTAACGGAGCA<br>ATCGCTGATATTATTAAAATTATCGGCGATGAATTCGTCTTTACTGATGATAT<br>AAAGGAACTGCGTTCTTACCGCGACCCTTATAATACAACTCGGGATGAAGAT<br>TTTGCTCCTTCGGCTGCCGTCGCTCCCAAAACGTTGGAGCATATCCAAAAAA<br>TTCTGGCCGTGGTCAATCAGTATAAGATTCCTGTTTGGACGATCAGTACCGG<br>TAAAAATTTTGCGTACGGTGGTGCTGCCCCGCGCAAACCGGGATATATCGTC<br>TTAGACCTGAAGCGTATGAATCGTATTATAGAAGTTAATGAAAAACATGGTT<br>ATGCTATCGTTGAGCCAGGGGTCAGCTACTATGACCTTTATAACTATATTCA<br>AGAAAAGGGATACAAGCTGTGGATTGATTGCGCGGCTCCTGCGTGGGGAGG<br>CATTGTAGGTAACACATCCGAGCATGGTGCAGGATACACACCATACGGTGA<br>CCACCTGCTTATGCAATGTGGGATGCAGGTTGTGCTGGCCGATGGTACAGTG<br>GTTGAAACAGGTATGGCTGGATCAGATAATTCAAAAACTGGTTCACTGTACA<br>AATATGGCCAAGGCCCGTGGGTCGATGGGCTGTTTACACAATCAAATTTTGG<br>TGTCATTACTCGTATGGGCATTTGGTTAATGCCAGAACCTCCGGGGTACACA<br>CCCTTTATGGTGACCCTGCCGCATGAACAAGATCTGGAATTAATCACAGACT<br>TGGCGCAGCCGCTTAAATTGAATATGGTGATTCCGAATGCGGCAACAACTGT<br>TGATCTGCTGTGGGAAGCCGCGATGTCACATACAAAGGCGCAGTATTACAC<br>AGGTAAGGGTCCCATGCCGGATTCCGCACGGAAAAAACTCGGTGAGGATCT<br>CGGACTCGGCGTGTGGAATTTCTATGCTGCTGTATACGGCCCACCGCCAATC<br>ATGAAAAATAACATGGACTTAATCAGAGATACGTTCAGCAAAATTCCTGGT<br>GCAGAATTTCATTATGACGACACTAGAAAGGGCTCTCCAGGATGGGACTAC<br>CGCGTTGAGCTCATGCGAGGCAAACCAAACATGACAGAATTCTCGCTGCTC<br>AATTGGGTTGGTGGCGGTGGTCATATCAATTTTAGCCCTATTAGCGCGGTGA<br>ATGGTAAAGACGCCTATCGACAGTTTGATATGATTCGCAACCGAGCGCATG<br>AATTTGACTTCGACTACATTGGGGAATTTCTAGTCGGCTGGAGGGACCAGCA<br>TCATATTTTTATGCTGATGTTTGACAGAATGAATGTGGAAGAGAAAAAACGT<br>GCACATGAACTGTTTGATGTTCTGATCGATGATGCCGCGGATCATGGATATG<br>GTGAATATCGTACCCATCTCAGCTTTATGGATAAAATAGCCAAAAGTTATGG<br>TTGGAATGATAATGCGCTGCTGAAGGCGCAGCAGAAAATCAAAGATGAATT<br>AGATCCGAATGGGATACTTTCTCCGGGCAAGATGGGCATTTGGCCGAAGCG<br>GATGCGCGATGATGCG | MNKPLPQGISLEQFNGAIA<br>DIIKIIGDEFVFTDDIKELR<br>SYRDPYNTTRDEDFAPSA<br>AVAPKTLEHIQKILAVVN<br>QYKIPVWTISTGKNFAYG<br>GAAPRKPGYIVLDLKRMN<br>RIIEVNEKHGYAIVEPGVS<br>YYDLYNYIQEKGYKLWID<br>CAAPAWGGIVGNTSEHG<br>AGYTPYGDHLLMQCGMQ<br>VVLADGTVVETGMAGSD<br>NSKTGSLYKYGQGPWVD<br>GLFTQSNFGVITRMGIWL<br>MPEPPGYTPFMVTLPHEQ<br>DLELITDLAQPLKLNMVIP<br>NAATTVDLLWEAAMSHT<br>KAQYYTGKGPMPDSARK<br>KLGEDLGLGVWNFYAAV<br>YGPPPIMKNNMDLIRDTFS<br>KIPGAEFHYDDTRKGSPG<br>WDYRVELMRGKPNMTEF<br>SLLNWVGGGGHINFSPISA<br>VNGKDAYRQFDMIRNRA<br>HEFDFDYIGEFLVGWRDQ<br>HHIFMLMFDRMNVEEKK<br>RAHELFDVLIDDAADHGY<br>GEYRTHLSFMDKIAKSYG<br>WNDNALLKAQQKIKDEL<br>DPNGILSPGKMGIWPKRM<br>RDDA |
| pZE-GbAAO<br>(HCU90459.1) | ATGGCAGTAAATCGCCTGCCAGAAGGTGTTTCCGCGGAAGCCTTTGCTGCTG<br>CCTGTGATGAGATGAGGGGTGCTGTTGGTGATAAATGGCTTATTCGCGACAA<br>CCTGCAGCATCTGAACTCATATCGAGATAGCTATACACCGGGTGACATTGAC<br>TCTAACGCGCCAAGCGCAGCAGTTGCCCCAAAAGACGTGGCTGAAATCCAA<br>AAGGTACTGGCAGTGGCCCGGAAGTATCGTACCCCCTTATGGACCGTGTCAA<br>CCGGGCGGAATTACGGATACGGTGGCGCGGCTCCGCGCAAAGCAGGCTGCA<br>TCGTTCTGGATCTAAAACGTATGAATCGGATCCTGGAAGTGAATGAGAAAC<br>ATGCATATGCCGTAGTAGAACCCGGAGTTACCTATCATGAATTGTACGAGTA<br>CTTACAAAGACGTGATATAAAACTTTGGGCCGACCCAGCGGCCCCTGCGTG<br>GGGCGGAGTAGTGGGCAATACGGCAGATCGGGGTGTGGGCTATACACCGTA<br>TGGTGAACACTTTACAGTGCAGTGCGGCATGCAAGTCGTACTGGCCGATGGT<br>AGCGTAGTCGAGACTGGCATGGGTGCCACAGCCAATGCCAGCACTCGCAAT<br>CTCTATAAATATGGTCATGGTGCATGGGTGGATGGGATTTTTACCCAGTCAA<br>ACTTCGGTGTAATCACACAGATGGGTGTATGGCTCTTGCCTGAGCCGCCTGG<br>CTACAGACCGTATATGGTTACCTTTCCCAAAGAAGATGATATCCATCAGATC<br>ACTGAAGTAGTTCGCCCACTGAAAATAAGCATGCTGATCCCCAATGCGGCG<br>GTAACAGTCGGACTGATATGGGAAGCAGCTGCGAAAGTATCAAAATCCCAG<br>TATTATGCGGGGTCTGGTGTAATGCCCGAATCGGCCCGCCGTAAGATGGCAA<br>CTGACTTAGATATAGGGAACTGGAATTTTTATGGAGCATTATACGGTCCGGA<br>ACCTATAATGGATAACAATTGGGCTATTATTGAGGAATCATTTGCCCAAGTT<br>CCAGGTGCCAAGTTTTACTTTGCAGAAGACCGTAAAGGTGATGCCGCCTTTG<br>ATTACCGGGCCAAACTGATGCGCGGAGTGCCGAACATGACAGAATTTGGCT<br>TACTGAACTGGCTGGGGCCAGGGAGCCATATTGATTTTAGCCCGATGAGTCC<br>CGTCACAGGTGATGGAGCTCTGCAGCAGTTCCATTTTATGCGGGATACGTGC<br>CATGAATATGGTGTAGATTATATTGGGGAATTTCTGATTGGTTGGCGGGATA<br>TGCATCATGTACTGATGCTGGTATTTAAACGCGATGATGAAGCCCACAAAAG<br>AACCATTAAAGAGTTATTTAAGGTTTTGGTGGGTGAAGCGACATCCAAAGG<br>GTATGGTGAATACCGTACCCACTTAGACTATATGGACATGATTTCTGGTACG | MAVNRLPEGVSAEAFAA<br>ACDEMRGAVGDKWLIRD<br>NLQHLNSYRDSYTPGDID<br>SNAPSAAVAPKDVAEIQK<br>VLAVARKYRTPLWTVST<br>GRNYGYGGAAPRKAGCI<br>VLDLKRMNRILEVNEKHA<br>YAVVEPGVTYHELYEYLQ<br>RRDIKLWADPAAPAWGG<br>VVGNTADRGVGYTPYGE<br>HFTVQCGMQVVLADGSV<br>VETGMGATANASTRNLY<br>KYGHGAWVDGIFTQSNF<br>GVITQMGVWLLPEPPGYR<br>PYMVTFPKEDDIHQITEVV<br>RPLKISMLIPNAAVTVGLI<br>WEAAAKVSKSQYYAGSG<br>VMPESARRKMATDLDIGN<br>WNFYGALYGPEPIMDNN<br>WAIIEESFAQVPGAKFYFA<br>EDRKGDAAFDYRAKLMR<br>GVPNMTEFGLLNWLGPGS<br>HIDFSPMSPVTGDGALQQ<br>FHFMRDTCHEYGVDYIGE<br>FLIGWRDMHHVLMLVFK<br>RDDEAHKRTIKELFKVLV<br>GEATSKGYGEYRTHLDY<br>MDMISGTYNWNNSALAR |

| | | |
|---|---|---|
| | TACAATTGGAATAACAGTGCCCTAGCTCGCGTTAATCAAAGCCTAAAAGAT<br>GTGCTGGATCCCAACGGCATTCTGTCTCCGGGTAAGAGTGGTATTTGGCCGC<br>AAGATGACCGTAAAGGT | VNQSLKDVLDPNGILSPG<br>KSGIWPQDDRKG |
| pZE-MrAAO (WP_087135908.1) | ATGTCAGAGAAAAATGTCCAGGAATACGATTATGTAGTGGTGGGAGCTGGC<br>TCTGCTGGCGCGGCTGTGGCTGCTCGGCTGTCTGAGGATCCGGAAGTCACCG<br>TTGCCCTTATAGAGGCAGGACCTGACGACGTAGGCGACGATGCGATTCTGA<br>AACTGGATCGTTGGATGGATCTTTTAGAAAGCGGGTACGATTGGGACTATCC<br>AATAGAGCCGCAGGCGCATGGTAACTCCTTTATGCGTCATGCCCGCGCTAAG<br>GTTATGGGTGGCTGCAGCAGTCACAATAGTTGCATTGCGTTCTGGGCACCGA<br>GAGAAGACATCGACGAATGGGAACAGCGTTTCGGCGCCACTGGCTGGAACA<br>GTGATATGGCATACCGACTCTATAAGAAGCTTGAAACAAATGAGGATGCGG<br>GTCCTGACGCACCACATCACGGCGATTCTGGCCCGGTGCACCTTATGAACGT<br>GCCCGCGAAAGACCCCTGTGGGGTGGCACTCCTTGAAGCCTGTGAACAGGC<br>GGGCATCCCTCGCGTTCATTTTAATACCGGTGAGACAGTTGTAAACGGCGCA<br>TCCTTTTTTCAGATTAACCGTCAGGCCGACGGTACGCGTGCATCTTCGTCAGT<br>GTCATATATCCATCCCATTAGACAGCGTCCTAACCTGCATGTGGTGACCGGA<br>CAGCAGGTAACTCGTATCCTTTTTGATGATGACCGCAGAGCGACCGGAGTGG<br>CTACCGCAGGTAGTTCCTTCGGGGCTGGGGGACGCATTGATGCGCGCAGAG<br>AGGTAATTCTGTCTGCCGGTGCCATCGGTTCACCTAAGCTCCTAATGTTAAG<br>CGGCATTGGCCCTGCAGAACATCTTGCGGAGGTTGGCGTAGACGTGCTTGTG<br>GATTCACCGGGCGTCGGCGCCAACCTCCAGGACCATCCGGAGGGTGTTATCC<br>AATGGGAAGCGAAAAAACCAATGGTGCAGACATCTACTCAGTGGTGGGAGG<br>CCGGGATCTTTACTACAACGGAGGAAGGCTTAGACCGGCCGGATTTAATGTT<br>TCATTACGGGTCAGTACCCTTTGATATGCATACAATTCGACAGGGCTATCCG<br>ACGAGTGATAATGCTTTTTGTTTGACTCCTAATGTTACCCATGCGCGTTCTAG<br>AGGCACCGTACGTTTACGCTCTCGTGATTGGAGAGATAAACCGAGAGTCGA<br>TCCCAACTATTTCACCGATCCGCATGACATGAGGGTGATGATCTTCGGAATA<br>AGAAAAGCAAGGGAAATAGTCGCAGAGCCAGCCATGGCCGAATGGGCAGG<br>TGAGGAACTGTTTCCGGGCGCAGAAGTACAAACAGACGAACAGATTGCAGA<br>TTACATTACTCGAACCCATAATACTGTGTATCATCCAGCAGGAACCGTCAGA<br>ATGGGCGCGGTAGACGACGAAGGAAGTCCCCTGGATCCGGAACTGCGCGTT<br>AAAGGCGTGAGTGGCTTGAGAGTGGCTGACGCAAGTGTGATGCCAGAGCTT<br>ACAACAGTCAATCCAAATATCACAACTATGATGATTGGGGAACGTTGCGCT<br>GAGTTAGTGTCTGGAAGG | MSEKNVQEYDYVVVGAG<br>SAGAAVAARLSEDPEVTV<br>ALIEAGPDDVGDDAILKL<br>DRWMDLLESGYDWDYPI<br>EPQAHGNSFMRHARAKV<br>MGGCSSHNSCIAFWAPRE<br>DIDEWEQRFGATGWNSD<br>MAYRLYKKLETNEDAGP<br>DAPHHGDSGPVHLMNVP<br>AKDPCGVALLEACEQAGI<br>PRVHFNTGETVVNGASFF<br>QINRQADGTRASSSVSYIH<br>PIRQRPNLHVVTGQQVTRI<br>LFDDDRRATGVATAGSSF<br>GAGGRIDARREVILSAGAI<br>GSPKLLMLSGIGPAEHLAE<br>VGVDVLVDSPGVGANLQ<br>DHPEGVIQWEAKKPMVQ<br>TSTQWWEAGIFTTTEEGL<br>DRPDLMFHYGSVPFDMH<br>TIRQGYPTSDNAFCLTPNV<br>THARSRGTVRLRSRDWR<br>DKPRVDPNYFTDPHDMR<br>VMIFGIRKAREIVAEPAM<br>AEWAGEELFPGAEVQTDE<br>QIADYITRTHNTVYHPAG<br>TVRMGAVDDEGSPLDPEL<br>RVKGVSGLRVADASVMP<br>ELTTVNPNITTMMIGERC<br>AELVSGR |
| pZE-RuAAO (TAJ20364.1) | ATGGAGTCCGTATTACCACCAGGTATCAAACGCGAACAGTTTGCGGCGCTAC<br>TGGCCGATCTGAAGAAAGTGGTTGGCCGAGAGTTCGTATTTACAGATACTCA<br>ATGGGAGCTTCCTGCGTATAACGATGCATATCTTACAACCCCAGCAGAACTT<br>CACCAGCCGTCGGCCGCGGTGGCGCCAGCGAATGTTGAGGAGTTACAGAGA<br>GTACTGGAAGTTGCTCGTCATTACAAGGCACCTTTGTGGACTATTAGCACCG<br>GTAAAAATTTTGCCTATGGAGGTCCTGCGCCGCGCAAGGCCGGCTACATCGT<br>TTTGGACCTGAAACGGATGAATCGCATTTTAGAGGTGAATGAAAAACATGG<br>TTATGCTGTTGTCGAACCTGGTGTAAGCTATATGGATTTATACCGGCACCTC<br>CAGAAAATAGGATCTAAACTATGGGTTGACTGTGCAGCACCGGGATGGGGT<br>GGTGTCCTGCCTAATATGACGGAACATGGAGTAGGTTATACGCCTTATGGTG<br>ATCATGTCACGATGCAGTGTGGGATGGAAGTTATGTTGGCAGATGGGACCTT<br>GGTGCAGACGGGAATGGGAGCATTGCCCGGCTCTCAGACGAATCATTTATA<br>CAAATACGGTCTTGGCCCCAGTGTCGATGGCCTTTTTACCCAGAGCAATTTC<br>GGTGTTGTGACAAAGGTCGGCATGTGGTTAATGCCTGAACCGCCCGGGTACC<br>GCCCCTATATGATCACTTTTGAGAAAGAATCTGATCTGCACGCTATTACCGA<br>GGTTTTACGTCCACTGAAGGTCAATATGTTAATTCCGGCCGTTGCTATGACT<br>GTCGAGATGCTGTGGGAGGCAGCAGTCCAGATCACGCGCCGCGATTACTAT<br>ACCGGGAAAGGCCCGATTCCAGACAGTGTACGAAAGAAAATGAGAAGTGAT<br>CTGAAAATTGGCGCATGGAATTTTTATGGTGCGCTGTACGGTCCGCCCCCTA<br>TGATGGATAATACATGGGAAGTGATACATGACGCATTTGCCTCGATCCCGGG<br>AGCCAAATTTTACTTTGATGAAGATCGTTCCAAAGATGATGTTGCCTGGCAG<br>TACCGGAAAAAATTAGGTGCGGGCATCCCCAATATGACAGAATACAATGTT<br>ATGAACTGGATCCCGAATGGTGCACATATAGACTTCTCACCAATCAGTCCTG<br>TGACCGGGGCGGATGCGACGAAACAATACGAATTAATTCGTGATCGTTGCA<br>ACAAAGCCGGGTTTGACTATTGCGGAGAGTTTGCAGTTGGATGGCGTGACAT<br>GCATCACATCTTCTGCCTTACGTTCGATCGGAATGATGCCAAGCAGAAAGCA<br>AGGGCGAACAAACTGTTTGGAGATCTCGTCGATGCGGCCGCTGCCGCCGGC<br>TATGGAGAGTACCGAACCCATGTTGACTTTATGGATCGTATCGCTGGTACCT<br>ATAATTGGAACGATAGTGCCCTGCTACGGATGCATGAAAAGGTCAAAGATG<br>TAGTCGATCCGTCCGGCATATTATCACCTGGTAAAATGGGGATATGGCCGAA<br>AAGTATGAGGAAAGGTAAAGCA | MESVLPPGIKREQFAALL<br>ADLKKVVGREFVFTDTQ<br>WELPAYNDAYLTTPAELH<br>QPSAAVAPANVEELQRVL<br>EVARHYKAPLWTISTGKN<br>FAYGGPAPRKAGYIVLDL<br>KRMNRILEVNEKHGYAV<br>VEPGVSYMDLYRHLQKIG<br>SKLWVDCAAPGWGGVLP<br>NMTEHGVGYTPYGDHVT<br>MQCGMEVMLADGTLVQT<br>GMGALPGSQTNHLYKYG<br>LGPSVDGLFTQSNFGVVT<br>KVGMWLMPEPPGYRPYM<br>ITFEKESDLHAITEVLRPL<br>KVNMLIPAVAMTVEMLW<br>EAAVQITRRDYYTGKGPIP<br>DSVRKKMRSDLKIGAWN<br>FYGALYGPPPMMDNTWE<br>VIHDAFASIPGAKFYFDED<br>RSKDDVAWQYRKKLGAG<br>IPNMTEYNVMNWIPNGA<br>HIDFSPISPVTGADATKQY<br>ELIRDRCNKAGFDYCGEF<br>AVGWRDMHHIFCLTFDR<br>NDAKQKARANKLFGDLV<br>DAAAAAGYGEYRTHVDF<br>MDRIAGTYNWNDSALLR<br>MHEKVKDVVDPSGILSPG<br>KMGIWPKSMRKGKA |
| pZE-GhAAO (WP_066166971.1) | ATGTCCACCGTCGTCGATGGCGAAGTCCGCGAGTTCGATTATGTTGTTGTGG<br>GAGGAGGCTCGGGAGGCTGCGCTGCGGCGGCACGCTTGAGTGAAGACCCAA<br>GCGTGTCAGTGGCCCTGATCGAAGCAGGGCCTGATGATCGTGGAATTCCAG<br>AGGTACTTCAGCTGAATAGATGGATGGAGCTACTAGAGAGCGGGTATGATT<br>GGGATTACCCGATCGAGCAGCAAACGCATGGCAATAGCTTTATGCGTCATG | MSTVVDGEVREFDYVVV<br>GGGSGGCAAAARLSEDPS<br>VSVALIEAGPDDRGIPEVL<br>QLNRWMELLESGYDWDY<br>PIEQQTHGNSFMRHARAK |

| | | |
|---|---|---|
| | CGCGTGCAAAAGTTCTGGGTGGTTGCTCGAGCCACAATAGCTGCATCGCATT<br>TTGGCCTCCTGCAGAAGATATGAACTCATGGGAAAGCACCTACGGAGCTAC<br>GGGGTGGAACGCTGATGCGCTGTGGCCAATTCTACAGCGACTGGAAACGAA<br>TGAGGACGCAGGCCCAGATGCCCCGCATCACGGTGACTCCGGTCCTGTCAAT<br>TTAATGAACGTTCCCCCCACTGATCCGTGCGGTGTTGCCTTACTTCAAGCCG<br>CGAGCGAGTTTGGCATTCCGACTGTGACATTTAATACAGGAACCACAGTCCG<br>TTCGGGGGCTAATTTCTTTCAAATTAACAGTAGACGCGATGGGACTCGGGCA<br>TCGTCCTCTGTTAGCTATATCCATCCTATTGCCGATCGCGAAAACTTCACCCT<br>GCTGACTGGCCTTAGAGCAAAGAAACTCCTCCTGGACGGGCTGCGCTGTGTG<br>GGTGTTGAAGTTGTCGATAACCAGTTTGGAAAAACACAGGTGGTAAGGGCG<br>GGTCGTGAACTAGTCCTTGCCGCAGGCGCGATAGATACGCCGAAATTATTGA<br>TGTTATCAGGAATAGGTCCGGCTGAGCATCTAGCAGGCCGCGGTGTTGAAAT<br>CGTGCTGGATAGCCCTGGAGTTGGTAGTAATTTACAGGATCACCCTGAAGGA<br>GTTATTAGTTGGGAAGCTAAACAACCGATGGTAACGGATAGCACCCAGTGG<br>TGGGAGATAGGGATATTCGATACTGTCGATTCTGGTTTAGATAGACCGGACT<br>TAATGATGCACTACGGCTCTGTGCCTTTTGACATGCATACAGTCCGCCGTGG<br>CTATCCTACCGCGGAGAATGTGTTCTGCCTGACTCCGAATGTTACCCATGCG<br>CGTAGCAGGGGTACCGTTCGTCTCGCGTCTCGTGATTTCCGCGACAAGCCTC<br>TTGTAGATCCAAAATATTTTACTGACCCGGAGGGCTATGATATGAGAATTAT<br>GACGGAAGGTATCCGGCGCGCTCGCGAAATAGTGAAACAACCCGCCATGGC<br>CGACTGGGCCGGTCGCGAACTGTTTCCCGGTCCAGATACCGTAGAAGATAG<br>AGATTTAGAAAAGTACATAATAAGAACTCATAACACAGTGTATCATCCAGTT<br>GGCACTGCTGCTATGGGCTCCGCAGACGATGCTCCGGTGGATGCGGAACTG<br>AGAGTGAAGGGTATTGATGGGCTCCGGGTTGCGGATGCCAGCGTTTTCCCAG<br>AACATATTAGTGTTAATCCAAACATTACCGTCATGATGATCGGTGAAAAATG<br>CGCGGATTTAGTTAAGGCGGCACGT | VLGGCSSHNSCIAFWPPA<br>EDMNSWESTYGATGWNA<br>DALWPILQRLETNEDAGP<br>DAPHHGDSGPVNLMNVP<br>PTDPCGVALLQAASEFGIP<br>TVTFNTGTTVRSGANFFQI<br>NSRRDGTRASSSVSYIHPI<br>ADRENFTLLTGLRAKKLL<br>LDGLRCVGVEVVDNQFG<br>KTQVVRAGRELVLAAGAI<br>DTPKLLMLSGIGPAEHLA<br>GRGVEIVLDSPGVGSNLQ<br>DHPEGVISWEAKQPMVTD<br>STQWWEIGIFDTVDSGLD<br>RPDLMMHYGSVPFDMHT<br>VRRGYPTAENVFCLTPNV<br>THARSRGTVRLASRDFRD<br>KPLVDPKYFTDPEGYDMR<br>IMTEGIRRAREIVKQPAM<br>ADWAGRELFPGPDTVEDR<br>DLEKYIIRTHNTVYHPVGT<br>AAMGSADDAPVDAELRV<br>KGIDGLRVADASVFPEHIS<br>VNPNITVMMIGEKCADLV<br>KAAR |
| pZE-MkAAO<br>(WP_15676174<br>7.1) | ATGAGTTCTGAACATACATTGACTCAGACCGATTTTGATTATATAGTTATTG<br>GGGGGGGAAGTGCAGGCGCGGCGGTAGCCGCTCGTCTGTCTGAAGATCCCG<br>CGGTCGAGGTGGCTCTCGTCGAAGCTGGGCCAGATGATGCTGATCTGCCCGA<br>GATCCTGCAACTGGATCGCTGGATGGAGCTGTTGGAATCGGGTTACGACTGG<br>GATTATCCGATTGAACCGCAGGAAAACGGAAATAGTTTTATGCGGCATGCTC<br>GTGCCCGAGTGATGGGTGGGTGCTCATCTCATAATTCTTGTATTGCTTTTGG<br>GCACCTAGAGAAGATTTAGATGAGTGGGAACAAAAATTCGGTGCGACCGGT<br>TGGAACGCGGAAAACCTTTATCCCTTATTTCAGCGCTTAGAAACCAACGAAG<br>ATGCTGGTCCGGACGCGCCACATCACGGTGACTCTGGCCCTGTTCATTTAAT<br>GAATGTCCCACCGAACGATCCTTCAGGCGTTGCGCTGTTAGACGCAGCGGA<br>GCAAGCTGGCATTCCGCGCGCCACTTTCAACTCAGGTGAGACGGTGGTCAAC<br>GGGGCAAACTTTTTCCAGATTAACCGTAGAGCGGATGGAACACGGTCTTCCA<br>GCTCGGTCAGCTATATTCATCCTATACGTGATCGTAAAAACTTTCATCTCCTT<br>ACCGGCCTGCGGGCTAAAGAGCTTCTCTTTGATGGGGACGCTTGTGCAGGTG<br>TTGCGGTTGTTGATAATGCATTCGCGCGGACTCACGCCCTGAGCGCACGCCG<br>CGAGGTGGTCTTATCTGCAGGCGCTATAGACAGCCCGAAACTTCTGATGTTA<br>AGTGGTATCGGCCCTGCAGCCCACCTGGCGGAGAATGGTATTGAAGTTCGC<br>GTTGACTCCCCAGGTGTGGGGGAAAACCTTCAGGATCATCCAGAAGGCGTA<br>ATTCAGTGGGATGCGAAGAAACCCATGGTGACCGAGTCCACACAGTGGTGG<br>GAAATTGGCGTGTTTACTCCGACTGAAGAGGGGTTAGACCGTCCTGACTTAA<br>TGATGCATTATGGCAGTGTTCCGTTTGATATGCATACTCTGCGACAGGGTTA<br>TCCGACTACCGAAAATGGCTTTTGTTTAACGCCTAATGTGACCCATGCTCGT<br>TCTAGAGGTACGGTTCGGTTACGTTCGCGCGATTTTCGTGATAAGCCTAAAG<br>TAGATCCCAGATATTTTACAGATCCCGAGGGCCACGATATGAGAGTCATGGT<br>CGCAGGTATTCGCAAAGCCCGCGAAATCGTCAGCCAGCCGGCCATGGCTGA<br>ATGGGCCGGAGAAGAATTGTATCCAGGTGCCGATGCACAGACAGATGAGCA<br>GCTGCAAGACTATATTCGCAAAACACATAACACTGTGTACCATCCAGCGGG<br>GACCGTTCGTATGGGTGCCGCGGATGATGAGATGGCGCCCCTCGACCCTGA<br>ATTACGCGTGAAAGGGGTAACCGGTTTACGCGTGGCTGATGCCAGCGTTATG<br>CCGGAACTTGTTACTGTTAACCCGAATATAACAACGATGATGATCGGCGAAC<br>GTTGCGCGGATTTAATCAGAGGTGCC | MSSEHTLTQTDFDYIVIGG<br>GSAGAAVAARLSEDPAVE<br>VALVEAGPDDADLPEILQ<br>LDRWMELLESGYDWDYP<br>IEPQENGNSFMRHARARV<br>MGGCSSHNSCIAFWAPRE<br>DLDEWEQKFGATGWNAE<br>NLYPLFQRLETNEDAGPD<br>APHHGDSGPVHLMNVPP<br>NDPSGVALLDAAEQAGIP<br>RATFNSGETVVNGANFFQ<br>INRRADGTRSSSSVSYIHPI<br>RDRKNFHLLTGLRAKELL<br>FDGDACAGVAVVDNAFA<br>RTHALSARREVVLSAGAI<br>DSPKLLMLSGIGPAAHLA<br>ENGIEVRVDSPGVGENLQ<br>DHPEGVIQWDAKKPMVT<br>ESTQWWEIGVFTPTEEGL<br>DRPDLMMHYGSVPFDMH<br>TLRQGYPTTENGFCLTPN<br>VTHARSRGTVRLRSRDFR<br>DKPKVDPRYFTDPEGHD<br>MRVMVAGIRKAREIVSQP<br>AMAEWAGEELYPGADAQ<br>TDEQLQDYIRKTHNTVYH<br>PAGTVRMGAADDEMAPL<br>DPELRVKGVTGLRVADAS<br>VMPELVTVNPNITTMMIG<br>ERCADLIRGA |
| pZE-EwAAO<br>(WP_02662248<br>9.1) | ATGCCGGATTTTATCGTGGTTGGAGGTGGATCCGCGGGTTGTGCAATTGCCG<br>GTCGCCTTTCTGAGGACCCGGATGTGTCGGTCACCCTGTTTGAGGCTGGCCC<br>CCGTGATAGCAGCATCTGGATTAGATTTCCCGTAACGTTTTATAAGTCTTTTA<br>AATCAAGCCTGCTCCATTGGTACAAAATCGAAAAGCTGAAACATCAAAATG<br>ATCTGGAAACCCAAGTCGGACAGGCGAGAGTGCTGGGGGGCGGTAGCAGTT<br>TGAACGCGATGATTTATATTCGCGGAGCACCGGAAGACTACGATCGATGGG<br>CAGCGCACGGAGCTGAGGGCTGGGGTTACAAAGATGTTCTGCCATATTTTAG<br>AAAAGCGGAGAACAATGAAGTCTATTCTAATGATGCCCACGGTCAGGAAGG<br>CCCTCTGAGTGTTTCAAATCAGCAGCACACGTTGCCGCTCACAAAGGCCTGG<br>GTGAAAGCGTGTCAGGAGGCGGGTATGCCATATAATCCGGATTTTAATTCAG<br>GTCAACTTCAGGGTGCTGGGCTGTATCAGCTGACCACGAAGAACGGACGGC<br>GCTGCTCTTCCGCAGACGCTTATCTTCATACAGCACGAAAACGGCGTAATCT<br>GAATATTGTCACTAATAAACAAGTCACCAAAATAATTGTTGAAGGGGGTAG | MPDFIVVGGGSAGCAIAG<br>RLSEDPDVSVTLFEAGPR<br>DSSIWIRFPVTFYKSFKSSL<br>LHWYKIEKLKHQNDLETQ<br>VGQARVLGGGSSLNAMIY<br>IRGAPEDYDRWAAHGAE<br>GWGYKDVLPYFRKAENN<br>EVYSNDAHGQEGPLSVSN<br>QQHTLPLTKAWVKACQE<br>AGMPYNPDFNSGQLQGA<br>GLYQLTTKNGRRCSSADA<br>YLHTARKRRNLNIVTNKQ<br>VTKIIVEGGRAVGVQYVE |

| | | |
|---|---|---|
| | AGCGGTGGGGGTGCAGTATGTTGAAAACGGTCGTTTGATGACAATGCGGGC<br>GGAACGAGAGGTGGTTGTATCCTCTGGTGCTATTGGTTCTCCACGGCTTCTG<br>CTGTTGTCGGGGATCGGCCCCGCATCTGATCTACAACGGGTAGGGGTTGATG<br>TAGTGCATGATCTCCCGGGTGTCGGTCAGAATCTGCAGGATCATACCGATTG<br>CTTCCTGATTTATAATCTGAAATCGAACACTTCCTATGATAAGTATAAAAAG<br>CTGCGATGGCAACTGGCTGCAGCGGCGCAGTATGCGATGTTTGGTAGTGGCC<br>CCATTACGAGTAATATTTGCGAGGGTGGCGCCTTTTGGTGGGGGGATCGTAC<br>CGATCCTATCCCGGACCTACAATATCATTTTCTTGCGGGGGCGGGTATAGAA<br>GAAGGTGTTGAGACTACAGCATCTGGTTCCGGTTGCACGTTAAATGTGTATG<br>CGTGTCGGCCTAAATCTCGCGGCAGAGTCGCACTGCGTAGCTCTGATCCATC<br>TGTCCCGCCTCTGGTTGACCCGAACTATCTTAGTCACCCACACGATGTAGAT<br>CGCCTGGTTGATGGCATTCGTTTTGGTCAGGAGATAATGGCCCAGCCATCTA<br>TGCGGAAATTTGTAAGCGAGGCGCACTTGCCGGAAAAACCTCTGAAAACGA<br>GGGCGGAATTCGAAGCATTTGTACGCAAATATACGCAGGGCGCCTACCACC<br>TCTCAGGTGCTTGTAAAATTGGAACAGACAAGATGGCTGTTGTAGACCCGCA<br>GCTGAGAGTCCATGGCATAGACGGTCTTCGTATCGCTGACACTAGCGTAATG<br>CCTTTCGTTACCAGTGGTAATTTAAATGCCCCGGCAATTATGATTGGTGAAC<br>GGGCCGCGGATTTCCTGAAGGGCAACCGCATT | NGRLMTMRAEREVVVSS<br>GAIGSPRLLLLSGIGPASD<br>LQRVGVDVVHDLPGVGQ<br>NLQDHTDCFLIYNLKSNT<br>SYDKYKKLRWQLAAAAQ<br>YAMFGSGPITSNICEGGAF<br>WWGDRTDPIPDLQYHFLA<br>GAGIEEGVETTASGSGCTL<br>NVYACRPKSRGRVALRSS<br>DPSVPPLVDPNYLSHPHD<br>VDRLVDGIRFGQEIMAQP<br>SMRKFVSEAHLPEKPLKT<br>RAEFEAFVRKYTQGAYHL<br>SGACKIGTDKMAVVDPQL<br>RVHGIDGLRIADTSVMPF<br>VTSGNLNAPAIMIGERAA<br>DFLKGNRI |
| pZE-StAAO<br>(AMU94088.1) | ATGGATCAGTTCGACATTATTGTGATAGGCGGTGGTAGCGCGGGAAGCGCT<br>GCGGCTGGAAGACTGGCCGAGGATGGAACGAGACGGATTTGCCTCATCGAA<br>GCGGGTGGTAGCAACGATAACATGTGGGTGAAAACCCCCGGTTTCATGCCTT<br>TCATCCCCAAAAAAAGTAATTATCAATATGACACATTGCCGCAAAAAGGCC<br>TGAACGGACGCACTGGTTACCAACCTCGGGGTAGAGGACTGGGTGGCTCTT<br>CTGCAATTAACGCTATGATTTATATTCGCGGCCATGCGTTTGATTACGATCA<br>GTGGGCGGCACTCGGCGCATCGGGATGGAGCTATGCTGATGTCTTACCGTAT<br>TTTAAGAGAAGCGAGGGGAATGAACGGGGGGCCGACCAGTGGCATGGTGG<br>GGATGGCCCGTTAAATGTCATGGATCAACGCTGGCCGAATGTCACAAGTAG<br>ACGTTTCGTGGAATCAGCTGCCGCACTCCAATTACCTCGCACTCCGGATTTC<br>AACGGCGCGCAGCAAGAAGGTTTCGGTCTGTATCAAGTTACGCAAAAAGGT<br>GGCGAACGGTGGTCTGCCGCCCGTGCGTATGTGGAGCCATTACGTGGGCGT<br>GCAAACTTTGCCATTCGGACAGGGGCCCTGGTTGAGAAAATCATTGTAGAA<br>AACGGTCGTGCTACCGGGGTGGCAATCCGTCGTGGCAAAGCAAGAGAAACT<br>CTTAGGGCAGCGGGTGGAGTTATTTTGTCGGCCGGCGCGTTCGGCTCGCCCC<br>AGTTGCTGATGTTGAGTGGTATTGGCCCTGCAGCGCATTTAAAAGCAATGGG<br>TATTGACCCTGTGGCTGATCGCGCGGCAGTGGGCGCAGATTTGCAGGACCAT<br>ATTGATTACGTGTCTAGCTGGGAGACGCGAAGCGGCGATCCATTCGGTGATA<br>GTCTTGCAGGGTCCTGGCGTATGCTGAAAGCCATATTCGAACATCGCACCGG<br>CCGCACCGGTATTATGACAACACCATTTGCCGAGGCAGGGGGTTTCTGGAA<br>ATCTCGTCCAGATGTAGCGGCCCCAGACATCCAGTTTCATTTTGTGCCAGCG<br>ATGTTAGAAGATCACGGACGTACTAAAGTTAAAGGCCACGGCTTTTCATGCC<br>ACGCCTGTGTGCTGCGGCCGGAATCCCGGGGTTCAGTTACTCTAGCATCCCC<br>AGATGCTGCAGCGGCGCCTTTGATAGATCCGGGATTTTTAACTGATGATCGC<br>GATATGGCCACACTGCGCGCCGGCGTACGAATGATGCACCGTATTGTTGCAG<br>CCCCACCGTTATCGGACTATGCAGGAGTGGACCGCCATCCGGTTAATATTGA<br>TGATGATGCAGCCCTGGATGCACTGATCCGCTCCCGCGCAGATACGGTTTAC<br>CATCCGGTGGGTACGTGTCGTATGGGGAGTGATGCCGAAGCAGTTGTAGAT<br>CCTACGTTGAAACTGAACGGAGTGGACGGTTTATGGGTGGCGGATGCCAGT<br>ATTATGCCGCGTCTGGTTAGCGGCAACACAAACGCTCCCTCTATCATGATTG<br>GAGAACGCGCGGCAGACTTTGTAAAATCCGCTTTACAG | MDQFDIIVIGGGSAGSAA<br>AGRLAEDGTRRICLIEAGG<br>SNDNMWVKTPGFMPFIPK<br>KSNYQYDTLPQKGLNGRT<br>GYQPRGRGLGGSSAINAM<br>IYIRGHAFDYDQWAALGA<br>SGWSYADVLPYFKRSEGN<br>ERGADQWHGGDGPLNV<br>MDQRWPNVTSRRFVESA<br>AALQLPRTPDFNGAQQEG<br>FGLYQVTQKGGERWSAA<br>RAYVEPLRGRANFAIRTG<br>ALVEKIIVENGRATGVAIR<br>RGKARETLRAAGGVILSA<br>GAFGSPQLLMLSGIGPAA<br>HLKAMGIDPVADRAAVG<br>ADLQDHIDYVSSWETRSG<br>DPFGDSLAGSWRMLKAIF<br>EHRTGRTGIMTTPFAEAG<br>GFWKSRPDVAAPDIQFHF<br>VPAMLEDHGRTKVKGHG<br>FSCHACVLRPESRGSVTL<br>ASPDAAAAPLIDPGFLTDD<br>RDMATLRAGVRMMHRIV<br>AAPPLSDYAGVDRHPVNI<br>DDDAALDALIRSRADTVY<br>HPVGTCRMGSDAEAVVD<br>PTLKLNGVDGLWVADASI<br>MPRLVSGNTNAPSIMIGER<br>AADFVKSALQ |
| pZE-RhAAO<br>(WP_05680765<br>4.1) | ATGACAGAAACTAATGAAGTTGATTTTCTTATTGTTGGAGGTGGATCTGCAG<br>GTTGCGTTTTAGCAAACCGCTTATCCGCCGATCCTGCAAATACAGTTGCACT<br>TATTGAAGCCGGCGGGGACGGGCGTGGTGCCTTAATTGATATGCCAGTCGG<br>CGCCGTTACCATGCTGCCAACGCGCCTGAACAACTGGGCGTTTGAAACTGTT<br>CCTCAGACTGGATTAAATGGCCGACGTGGCTATCAGCCGCGCGGTAAAGCC<br>TTGGGAGGGTCAAGTGCAATCAATGCTATGATTTACATCCGCGGCCATCGTT<br>CTGATTATGATCATTGGGCGGCCCTTGGTAACCCTGGCTGGAGTTACGATGA<br>GGTTCTTCCGTATTTTAAGAAAGCTGAAAACAATGAACGACTGCATGATGAA<br>TTTCATGGTTCAAGCGGCCCGCTTCATGTGGGCGAGTTACGCTCTGCAAACC<br>CGTGGCAGCAGGTGTGGCTGCAAGCGGCGCGGGAAGCCGGTTTTGCACTGA<br>ATGATGACTTCAACGGCGCGAGCCAGGATGGTATCGGACTGTATCAGGTGA<br>CCCAGCACCGTGGGGAAAGATGGTCAGCAGCTCGCGCTTATCTGCATCCTGT<br>AATGGATCGGCGTGATAACTTGCACGTTTTAACAAAGGCCCAAACATTGCGC<br>GTTCTCTTTGACGGAAAACGTGCTGTGGGCATAGAAATGGCTCGTGGTGGGC<br>ACGTAGAGCGCCTGATGGCAAGGAGAGAGGTCATTTTATGTGCCGGAGCAC<br>TGCAAACACCACAACTCCTCCAGGTGTCCGGGGTGGGTGACGGTGCCTTCCT<br>TCAGCGCACCGGTATCCATCTCAAACATCATCTGCCTGGGGTAGGTCGTAAC<br>CTACAAGATCATATAGATTTCACGTTTGGCTATCGTGTGCCAGATACGGCGC<br>TTTTCGGCTTTTCACCGGCGGGTTCGCTGTCAGCCTTACGCGCCTTTGGACGC<br>TACCGCCGGGAGAGGTCGGGTCCACTGACCAGCAACTTTGCCGAAGGAGGT<br>GGTTTCTTACGTGTAAGTCCGGAATCGACTGCCCCGGACGTGCAGTTGCACT | MTETNEVDFLIVGGGSAG<br>CVLANRLSADPANTVALI<br>EAGGDGRGALIDMPVGA<br>VTMLPTRLNNWAFETVPQ<br>TGLNGRRGYQPRGKALG<br>GSSAINAMIYIRGHRSDYD<br>HWAALGNPGWSYDEVLP<br>YFKKAENNERLHDEFHGS<br>SGPLHVGELRSANPWQQV<br>WLQAAREAGFALNDDFN<br>GASQDGIGLYQVTQHRGE<br>RWSAARAYLHPVMDRRD<br>NLHVLTKAQTLRVLFDGK<br>RAVGIEMARGGHVERLM<br>ARREVILCAGALQTPQLL<br>QVSGVGDGAFLQRTGIHL<br>KHHLPGVGRNLQDHIDFT<br>FGYRVPDTALFGFSPAGSL<br>SALRAFGRYRRERSGPLTS<br>NFAEGGGFLRVSPESTAP<br>DVQLHFVVALVDDHARK |

| | | |
|---|---|---|
| | TCGTTGTTGCCCTTGTTGATGACCATGCGAGGAAACTGCACCTGCGCCATGG<br>TCTGTCCTGTCACGTATGTCTGCTCCGGCCGCGCTCTAGAGGGACAGTGCTG<br>GTCCAGTCTGGCGATATTCGTGATGTGCCACTGATTGACCCTAAATATTTTG<br>ATGATCCCGATGACCTGGAAGTTTTGGTTGAAGGGTGGAAATTGACGCAAC<br>GGCTGCTCCAAGCCCCTGCTATTGCTAGTCGTGTTAGACGTGATCTGTTCACT<br>GCTGGCGTGTCTAGCGACCATGAAATCCGGGAGGTGATTAGAAACAGAGCC<br>GACACTATTTATCACCCGAGCGGTACATGTCGCATGGGTGCCGACGCTATGG<br>CGGTTGTTGACGCGACCTTGCGCGTCCATGGTCTGCAGGGGTTACGGGTAGT<br>AGATGCTTCCGTAATGCCTACCCTAATAGGAGGAAATACAAATGCCCCTACA<br>ATCATGATTGCGGAAAAAGCTGCCGACCTTATCTTAGCGGCG | LHLRHGLSCHVCLLRPRS<br>RGTVLVQSGDIRDVPLIDP<br>KYFDDPDDLEVLVEGWK<br>LTQRLLQAPAIASRVRRD<br>LFTAGVSSDHEIREVIRNR<br>ADTIYHPSGTCRMGADA<br>MAVVDATLRVHGLQGLR<br>VVDASVMPTLIGGNTNAP<br>TIMIAEKAADLILAA |
| pZE-CmAAO (WP_086507454.1) | ATGAGCGACAGTCGCGGGGATGATATGGCAGGAACGGAATACGATTACGTA<br>GTGCTTGGTGGTGGTTCAGCAGGGGCCGCCGTTGCCGCGAGACTTTCGGAAG<br>ATCCGGATATGCAAGTTGCACTTATAGAAGCCGGTCCACATGATAGAGATCT<br>AGATGTAGTTCTGCGTCTGGATCGTTGGATGGAACTTCTAGAAAGTGGTTAT<br>GATTGGGATTACCCTATTGAGCCTCAAGAGAACGGAAATAGCTCTATGCGG<br>CACGCACGGGCAAAAGTTTTAGGTGGATGTAGCAGCCATAATTCATGTATTG<br>CGTTTTGGGCCCCAAGGGAGGATCTCGACGGTTGGGCGCACGAACATGGCG<br>CCACAGGCTGGGATGCAGATTCCACTTATCCCCTGTATCAACGGCTGGAAAC<br>CAATGAGGACCCGGAACCGCACCATGGTCATGACGGTCCGGTACATCTTAT<br>GAACGTTCCTCCTCGAGATCCTTCTGGGGTTGCACTGTTAGATGCCTGTGAA<br>CAAGCGGGCATTCCACGTACGCGTTTCAACACCGGTGAAACTGTCCGGAAC<br>GGAGCCGGCTTCTTTCAGATCAATCGCCAGGCTGATGGTACTCGCGCATCTA<br>GTTCAGTTAGCTACTTACATCCCTTAGCTGATCGTGCCAATTTAACGGTTCTG<br>ACAGATCTGAAAGCACGTGCATTAGAATTGGATGATGATGACCGTTGTACTG<br>GAGTGCAAGTCGTGGATAATGCCTTTGCCCGGACTAGGACCATCAGGGCAC<br>GTCGCGAAGTGATTTTGTCGGCAGGCGCAATTGATAGTCCTAAATTGCTGAT<br>GTTAAGTGGTATTGGCCCAAAAGCCCATCTTGAAGATTTGGGTATTGCTGTT<br>AGAGTCGACTCTCCGGGAGTTGGAGAGCACCTGCAGGATCACCCTGAAGGG<br>GTTATTCAATGGGAAGCAAAACAGCCTATGCCAACTGAAAGCACACAGTGG<br>TGGGAGATAGGAATTTTTGCTACCACGGAAGAAGGCCTTGACAGACCGGAC<br>CTGATGTTTCATTATGGCTCGGTCCCCTTTGACATGCATACAGTTCGCCAGG<br>GCTACCCGACCACCGAAAACGGGTTTTGCCTGACACCGAATGTAACCCACG<br>CCCGTAGTAGAGGTACCGTTCGTCTACGTTCTCGTGATCACCGCGATAAACC<br>GTTAGTGGATCCGAGATACCTGACGGATCCACACGATATGCGGGTGCTGAT<br>AGCTGGAATTAGATTAGCAAGAGAAATTGTAGCCCAGCCTGCGATGGCCGA<br>ATGGGCCGGCCGAGAACTTTATCCCGGTGTTGAAGCACAGACAGATGAAGA<br>ACTTGCTGACTATATTTCTCGTACCCATAACACGGTGTATCACCCGGCTGGG<br>ACCGTAAGAATGGGCGCCGTGGATGACGCCATGTCGCCATTAGATCCGGAA<br>CTGCGGGTCAAAGGAGTCACAGGGTTACGTGTAGCTGATGCTTCCGTCATGC<br>CAGAGCTGACGACAGTGAATCCAAATATCACCACGATGATGATCGGAGAGC<br>GATGCGCGGATATGGTCAAAAGGGCCCGGGCAGAGGAACTGGTGGCCCGT | MSDSRGDDMAGTEYDYV<br>VLGGGSAGAAVAARLSE<br>DPDMQVALIEAGPHDRDL<br>DVVLRLDRWMELLESGY<br>DWDYPIEPQENGNSSMRH<br>ARAKVLGGCSSHNSCIAF<br>WAPREDLDGWAHEHGAT<br>GWDADSTYPLYQRLETNE<br>DPEPHHGHDGPVHLMNV<br>PPRDPSGVALLDACEQAG<br>IPRTRFNTGETVRNGAGFF<br>QINRQADGTRASSSVSYL<br>HPLADRANLTVLTDLKAR<br>ALELDDDDRCTGVQVVD<br>NAFARTRTIRARREVILSA<br>GAIDSPKLLMLSGIGPKAH<br>LEDLGIAVRVDSPGVGEH<br>LQDHPEGVIQWEAKQPMP<br>TESTQWWEIGIFATTEEGL<br>DRPDLMFHYGSVPFDMH<br>TVRQGYPTTENGFCLTPN<br>VTHARSRGTVRLRSRDHR<br>DKPLVDPRYLTDPHDMR<br>VLIAGIRLAREIVAQPAMA<br>EWAGRELYPGVEAQTDE<br>ELADYISRTHNTVYHPAG<br>TVRMGAVDDAMSPLDPE<br>LRVKGVTGLRVADASVM<br>PELTTVNPNITTMMIGERC<br>ADMVKRARAEELVAR |
| pZE-LmAAO (WP_108421602.1) | ATGAACCAGCAGGAATTCGACTATGTGATTGTTGGAGCCGGTAGCGCTGGG<br>TGCTGCGTAGCTGGACGCCTTTCCGAGGATCTGAACACTAGTGTGTGTTTAC<br>TGGAGGCAGGTGGTCCCGATGATAGCGTATTCGTTAAGGCGCCTCTGGGCTT<br>TGCTGCAACAAGCAGCCTGAAAATAAACTCTTGGGGTTACGAAACCGTACC<br>CCAAACGGGCTTTAACGGTCGGAAAGGGTTTCAGCCGCGTGGTAAAGTGAT<br>GGGAGGGTCCAGCTCCGTTAATGCCATGGTGTACACCCGCGGTAACCCTCTT<br>GACTATGACAATTGGTCCGCACTTGGCAATCAAGGTTGGTCTTATCAGGAGG<br>TTCTGCCTTATTTTAAAAAAAGTGAACACAACGAATGTTTCGGAGAAAACGC<br>ATACCGGGGTGTTAACGGACCCGTGAATGTGTGTTATCTGAGAAGCCCGAGT<br>CCACTCAATCAGGCCTTCATTCAGGCATGCAATGAGCAAGGGATCCCCTTTA<br>ATCCGGATTACAACGGTGCGCAACAGTTTGGAGTTAGCCCAGCACAGGTAA<br>CACAGAAGAATGGTGAGCGGTGGAATGCCGCAAGGGCATACCTGGACCCGC<br>ACAGGCATCAGGCAAACCTTCATGTCATAAGCCAGGCACATGCTTCACAAA<br>TTATGTTTGAAGGCAAACGCGCTATTGGCGTCAAATACTTCGTCAATGGGGT<br>TACGCACGAGGTACGTGCGCGTAAGGAAGTTGTCGTTAGCGGAGGTGCTTTC<br>GGATCGCCGCAACTTCTTATGCTGAGTGGCGTGGGTCCTGCGGATCACCTCA<br>AAGATTTAGGCATCCCGTTAGTACACGAACTGCCGGGCGTGGGTCAGAACTT<br>ACAGGACCATATTACCACGGTCCTGATTTACAAAACATCTAAAATCAATGAA<br>GCGATGGGTTTTTCATTAAAGGGTGCGGTTAACTTGGTGAAATCTATATTAG<br>AGTGGCGCTCAAAACGAACTGGATGGATTACGAGTAATGTGTCAGAAACTC<br>AGGGGTTTGTGTCTACTGAAGGAAATACCGCCTATCCGAACATCCAGCTGGC<br>TCTGTGTACGGGTATTGTCGATGATCATGCAAGGAAAATGCATCTGGGACAT<br>GGCTACACACTCCATGTTACGCTGATGCGCCCGAAAAGCCGGGGTACCGTTA<br>CTCTTGCATCTCGTAATCCAATGGATAAGCCTCTCATTGACCCGGCATTCTTC<br>AAACATCCGGATGACATAGAAACACTGGTTACGGCAACCCAGCTGGGACTG<br>CGGGTGATGGCCTCGCCGGGCTTAAATGCATATCGTGGAGAGATGCTCTACG<br>AAGTTGATCACAACCAGCCTGGGCAGATCAGAGATTTTCTTAAAGATCATTC<br>TGATACCGAATACCACCCAGTCGGCACTTGTAAGATGGGGCCCGTACAGGA<br>CCCACTGGCGGTGGTGGATGCAGAACTGCGCGTTCATGGTTTGAGTAGTCTC | MNQQEFDYVIVGAGSAG<br>CCVAGRLSEDLNTSVCLL<br>EAGGPDDSVFVKAPLGFA<br>ATSSLKINSWGYETVPQT<br>GFNGRKGFQPRGKVMGG<br>SSSVNAMVYTRGNPLDY<br>DNWSALGNQGWSYQEVL<br>PYFKKSEHNECFGENAYR<br>GVNGPVNVCYLRSPSPLN<br>QAFIQACNEQGIPFNPDYN<br>GAQQFGVSPAQVTQKNG<br>ERWNAARAYLDPHRHQA<br>NLHVISQAHASQIMFEGK<br>RAIGVKYFVNGVTHEVRA<br>RKEVVVSGGAFGSPQLLM<br>LSGVGPADHLKDLGIPLV<br>HELPGVGQNLQDHITTVLI<br>YKTSKINEAMGFSLKGAV<br>NLVKSILEWRSKRTGWIT<br>SNVSETQGFVSTEGNTAY<br>PNIQLALCTGIVDDHARK<br>MHLGHGYTLHVTLMRPK<br>SRGTVTLASRNPMDKPLI<br>DPAFFKHPDDIETLVTATQ<br>LGLRVMASPGLNAYRGE<br>MLYEVDHNQPGQIRDFLK<br>DHSDTEYHPVGTCKMGP<br>VQDPLAVVDAELRVHGLS |

| | | |
|---|---|---|
| | CGCGTTATTGATGCATCCGTTATGCCTCAGCTTGTAACGGGCAATACAAATG<br>CTCCGACTTTTATGATCGCAGAAAAGGCAGTGGATCTGATGCGTGGT | SLRVIDASVMPQLVTGNT<br>NAPTFMIAEKAVDLMRG |
| pZE-RoAAO<br>(WP_08542083<br>1.1) | ATGCAGGACGACTCTTTGAGCGCAGTTGATTATATTGTCATCGGCGGAGGTT<br>CTGCTGGCTGCACCCTGGCTGCTCGTCTGTCAGAAGATCCAGATGCGTCGGT<br>TTTGCTCATCGAGGCCGGTCAAAAAGACCGGAGCCCGTATATCCATCTGCCC<br>GTTACTTATTATAAGACAACAGGTCCCCAGTTTACCTGGGGTTTCGAGACGA<br>CCGCACAGGAACATCAAGGGGGCATTTCAACCCCTTTCACACAGGCACGGG<br>TTCTGGGTGGGGGTAGCTCCATTAATGCACAGGTTTATATCCGAGGAACCGC<br>TGCAGATTATGATGCCTGGAAATCGGACTACGGTTGCGCTGGCTGGGGATAC<br>GCAGATGTCCTGCCGTATTTTAAACGTGCTGAAGACAATCAGCGCCTGTCAG<br>GCGATCTCCATGGTAACGAAGGTCCGCTGAAAGTATCCGATCAAGCGTATA<br>CCCATCCACTCACCTACGCTTGGCTCAAGGCATGCCAACAAGTTGGGATGCC<br>ATATATCGACGACTTTAATCGTGAAAAACGGGGTGGTTGCGGGCTCTATCAA<br>GTGACTAATAGAGCAGGTCGGCGATCATCCACTGCAAATTGCTACCTGCGGC<br>CAGCGGCTTCGCGTCGCAACTTACTGGTACTTACGGATAGCCAAGCCATGAG<br>AATTCTCTTCGAAAATGGTCGGGCGTCAGCCGTGCAACTTCGTAGTGACACC<br>ATTACACGCACTATTGCAGCTCGGAGAGAGATCATCCTGACGTGCGGAGCG<br>ATTGGAACTCCTAACCTGTTACTGCATAGCGGCATTGGTGACGGGGCTATGT<br>TAGCTGGCAACGGCATTGATGTAGTAGCTGATTCACCGCAGGTAGGTCGTAA<br>CCTTCAGGATCATCTGGACGTATTCCTGGTGTATGAATTATTGGGCCCGTAT<br>AGCTACGATAAATATAAGAAAGCATGCTGGCAGGCTTGGGCCGGCTTGCAA<br>TTTGCACTGTTTCGCTCAGGTCCGGTGACAAGCAACGTGGTGGAGGGTGGCG<br>CATTTTGGTGGCTTGATGAAGGCGATCCGGAACCCGATGTGCAATTCCACTT<br>TTTAGCAGGAGCGGGTGTCGAAGCTGGGATCGCAGGGGTTCCCGGGGGTAA<br>TGGGTGTACTTTGAATGCGTATTTAACTAGGCCACGGAGCAGGGGTACAGTT<br>TCAATTGCCGGCAATGATCCTCTCTACCACCCGAAGATAGACCCGAACTACT<br>TGAGCGACCCTGATGACTTGGCTAAAACAGCTGATTCAGTTCGCTTGGGGCG<br>CAAGATTATGGCCGGACCAAGCTTAGCCCCTTATCTTAAAAATGAGCACTTT<br>CCGGGGGTAGATGTAGAATCCCAGCAGGAGATCGAGGATTTTGTGAGAGCA<br>CATGCGCGCACCGGATATCATCCTGCCGGCACCTGTCGGATGGGCGGCGAT<br>GCGGAATCTGTGGTGGACACCCAGCTGCGTGTGCGGGGTGTCGACGGCTTA<br>CGAATTGCTGACAATTCTGTCATGCCCAAATTAGTTAGCGGCAACACCAACG<br>CAACCGCCATCATGATTGGTGAACGTGCAGCGGACTTCATTCGAGGGAACG<br>GA | <br>MQDDSLSAVDYIVIGGGS<br>AGCTLAARLSEDPDASVL<br>LIEAGQKDRSPYIHLPVTY<br>YKTTGPQFTWGFETTAQE<br>HQGGISTPFTQARVLGGG<br>SSINAQVYIRGTAADYDA<br>WKSDYGCAGWGYADVL<br>PYFKRAEDNQRLSGDLHG<br>NEGPLKVSDQAYTHPLTY<br>AWLKACQQVGMPYIDDF<br>NREKRGGCGLYQVTNRA<br>GRRSSTANCYLRPAASRR<br>NLLVLTDSQAMRILFENG<br>RASAVQLRSDTITRTIAAR<br>REIILTCGAIGTPNLLLHSG<br>IGDGAMLAGNGIDVVADS<br>PQVGRNLQDHLDVFLVYE<br>LLGPYSYDKYKKACWQA<br>WAGLQFALFRSGPVTSNV<br>VEGGAFWWLDEGDPEPD<br>VQFHFLAGAGVEAGIAGV<br>PGGNGCTLNAYLTRPRSR<br>GTVSIAGNDPLYHPKIDPN<br>YLSDPDDLAKTADSVRLG<br>RKIMAGPSLAPYLKNEHF<br>PGVDVESQQEIEDFVRAH<br>ARTGYHPAGTCRMGGDA<br>ESVVDTQLRVRGVDGLRI<br>ADNSVMPKLVSGNTNAT<br>AIMIGERAADFIRGNG |
| pZE-AbAAO<br>(WP_02066050<br>4.1) | ATGACTCCCCCGGATGATAGCTTTGATTTCGTCATTGCTGGGGGCGGTACCG<br>CTGGTTGCGTCCTGGCTGCACGGCTCTCCGAAAATCCGGCAAATCGCGTGCT<br>CTTACTGGAAGCAGGCCGCGATGACCGTAATCCTCTGATCCATGTACCAGCA<br>GGCTTTGCAAAACTGACCAGCTCCAGCTACCAGTGGGGTTACACAACTGTAC<br>CTCAGCGTCATTGCGATAATCGAAGTGTGCAGTTTGCACAAGGCAAAGTTGT<br>GGGCGGTGGCGGTAGCATTAACGCTCAGGTGTATACCCGTGGCGCCGCAGA<br>GGATTATGATGAATGGGCGGCTGCGTACGGTTGCGAAGGATGGTCATCTAA<br>AGACGTGCAGCCGTATTTCCTGCGTTCTGAGGGTAACACGCGTCTGAGTGCT<br>CCGCACCACGGTACAGAGGGTCCCCTTGGTGTTTCTGACCCGGCCTCCAGTC<br>ATCCGTTATCGCAAGCATTCGTACGTGCAGGTCAGGAATTTGGTTTACCGTA<br>TAACGCAGACTTTAACGGCGCAGACCAATATGGGATTGGCTTTTATCAGACC<br>ACCACGAAGAATGGTAGAAGATGTAGCGCCGCGGTGGGATATCTGCGTCCG<br>GCGCGTAAACGTGCAAACCTTGTTGTAAGATCACAGGTGCTTGTTACCCGCG<br>TGCTGCTAGATGGTAGACGCGCCACAGGGATCGAATGCGTCGATGGGGGGC<br>GCTTGCGCAGATATCGTGCGGGACGCGAAGTCGTCGTTACAGCAGGTGCGTT<br>CGGTAGCCCGAAACTGCTACAATTATCAGGCATTGGAGATCCGGAAGATCT<br>GGGCCGTGCAGGTGTTGAAGTTCGGCACGCCCTGCCGGGTGTGGGGAAGAA<br>TCTCCAAGACCACTGCGACCTGGACATTATCTATCAGCTGCACAGGTATCAG<br>AGTATGGATCGCCTGCAGCGTCCAGTACCGGCAGCGGTAAGCGCCGGCATT<br>CAATATCTGGCGTTCCGTACTGGCCCGCTCGCAAGCACAGTTGTAGAAGCCG<br>GCGGATTTGGCAGATCCGATCCGGGAGAACCAACTGCCGATTTGCAATTCCA<br>CTTCTTGCCTGCCGCGGGCGTGGAAGCAGGGGTTGCGGGTGTACGTCCTGGC<br>TATGGAGCTACGCTTAATGTGGTGTTCCTCCGGCCGTACTCTAGGGGTACTG<br>TGCGAATTGCCTCCGCAGACCCGGCGCGTGCTCCCCTGATTGATCCCAATTA<br>CTTGGCTGACGAACGGGACGTAGCACGTACCGTCGATGGGGTTCGTCAGAG<br>CCGTGAAATTATGGCACAGCCGAGTATGGCTCGCGAGGTTAAATCCGAACA<br>TCTGGCCGGCCAAGCAACGCTGCGTACCACTGATGACTATCTTCGGTTTGTG<br>AGGGCGCACGGACGTACCGCCTACCACCCGGTGGGAACCTGCGCTATGGGC<br>GTTAATGACCAGGCGGTTGTAAGCCCGGAGCTTCGAGTTTACGGACTCGACG<br>GTTTACGCGTAGCGGACTCCAGCGTAATGCCACGAATCGTTTCGAGTAACAC<br>TCAGGCACCGACGGTGATGATTGCAGAGAAAGCGGCCGATTTGATTCTGGC<br>CGGT | <br>MTPPDDSFDFVIAGGGTA<br>GCVLAARLSENPANRVLL<br>LEAGRDDRNPLIHVPAGF<br>AKLTSSSYQWGYTTVPQR<br>HCDNRSVQFAQGKVVGG<br>GGSINAQVYTRGAAEDYD<br>EWAAAYGCEGWSSKDVQ<br>PYFLRSEGNTRLSAPHHG<br>TEGPLGVSDPASSHPLSQA<br>FVRAGQEFGLPYNADFNG<br>ADQYGIGFYQTTTKNGRR<br>CSAAVGYLRPARKRANL<br>VVRSQVLVTRVLLDGRRA<br>TGIECVDGGRLRRYRAGR<br>EVVVTAGAFGSPKLLQLS<br>GIGDPEDLGRAGVEVRHA<br>LPGVGKNLQDHCDLDIIY<br>QLHRYQSMDRLQRPVPA<br>AVSAGIQYLAFRTGPLAST<br>VVEAGGFGRSDPGEPTAD<br>LQFHFLPAAGVEAGVAGV<br>RPGYGATLNVVFLRPYSR<br>GTVRIASADPARAPLIDPN<br>YLADERDVARTVDGVRQ<br>SREIMAQPSMAREVKSEH<br>LAGQATLRTTDDYLRFVR<br>AHGRTAYHPVGTCAMGV<br>NDQAVVSPELRVYGLDGL<br>RVADSSVMPRIVSSNTQA<br>PTVMIAEKAADLILAG |
| pZE-MaAAO<br>(WP_10797924<br>0.1) | ATGAAACAGCAGAACGAAGTCCAGGAATTTGATTATGTTGTTGTGGGTGCTG<br>GTTCCGCGGGATGTGTGATTGCTAGCCGTCTGACGGAGGATCCGGCCGTTTC<br>CGTATGTTTGCTTGAAGCAGGTGGGCCAGATAGCAGCGTCCTGATACATGCA<br>CCAGCCGGGGTAGTTGCAATGGTCCCGCGAAAGATAAACAATTATGCCTAC | MKQQNEVQEFDYVVVGA<br>GSAGCVIASRLTEDPAVS<br>VCLLEAGGPDSSVLIHAPA<br>GVVAMVPRKINNYAYET |

| | | |
|---|---|---|
| | GAGACAGTTCCGCAACCCGGCCTCAACGGTCGCCGTGGTTACCAGCCGCGC<br>GGTAAAACACTGGGTGGAAGTAGTTCAATTAATGCGATGCTGTATGTCCGG<br>GGTTGCGCATGGGATTATGATAACTGGGCGGCACAGGGTAATCCGGGTTGG<br>AGCTATCCAGACGTACTGCCTCTGTTTAAAAGGGCCGAAAATAATGAAGATT<br>TTGGAGGCGATGATTTTCATGGCAGCGGAGGTCCTTTGAATGTGTGCTATCC<br>GCGGCATGCTTCCCCGATTAACCAAATGTTTATAGATGCTGCAGCTCAGAAT<br>GGCCTGCCGCAGGTGCAGGACTATAACGGTGCAACTCAAGAGGGCGCATTT<br>CTGTATCAGGTGACCCATAAGAATGGAGAGCGCTGCTCTGCCGCAAAGGGT<br>TATCTGACACCTCACCTCGGCCGCCCAAACCTGCATGTGGTCACCAACGCCG<br>TCTCTAGTCGTATATTGCTTGAAGAAGGCCGGGCAGCGGGTGTCGAGTATCA<br>TCATGATGGCCAGCTTAAACAAGTTCGCGCTCGTGCTGAGGTTGTCCTGAGC<br>GGAGGAGCCTTTGGTTCACCGCAGCTCCTGATGTTATCCGGAATCGGTCCGG<br>CAGCTCAGTTGCAGGCCCTTGGGATACCTGTGGTACGTGACCTTCCCGGCGT<br>GGGGCGGAATCTGCAAGACCATCTGGACCATGTCCAGGGTTGGAAGACGAG<br>ATCTGATACCGAAACCTTCGGTCTGTCACCTCGAGGAGTAGGCAAAATTGCT<br>GCAGCGATGATGGAGTGGCGATCTAAACGCACCGGAATGATCACGTCTCCC<br>TACGCGGAAGCCGGAGCCTTTCTGCGTTCCGCTCCAGATGTCCAGGTCCCCG<br>ATCTGCAATTGGTCTTTGTTTTGGCTCTGGTGGACGACCACGCGAGAAAAAT<br>GCACTTAGGGCATGGGATTTCCTGTCATGTGGATTTACTCCGGCCTCACTCT<br>AGAGGCGAAGTTACACTCCAGTCTCGCGATGCGAGGGATGCCCCGGTTATC<br>GACCCACGTTTTCTCAGCGACGAGCGTGATGTCGAAACATTGTACCGGGGCG<br>TCTGCTTGCAGCAAAAAATTATGGAATCGGCACCGCTGGATCCCGTGCGGG<br>GTAAAATGTTATACCCCGTAAGACGCGGTGATCGGGCCGCGACGATCGCCG<br>ATATCCGCAACCGCGCAGATACCCAGTACCACCCGGTTGGCACTTGCAAAAT<br>GGGCCCTCGAGACGACGCTTTAGCGGTCGTGGACGCACAGCTGCGTGTCCGT<br>GGTGTAACTGGCCTTCGCGTCGCAGACGCTAGTATCATGCCGACCATAGTCA<br>GCGGTAATACAAATGCTCCTTCAATTATGATTGGTGAAAAGTGTGCCGAACT<br>GCTTAAAGGG | VPQPGLNGRRGYQPRGKT<br>LGGSSSINAMLYVRGCAW<br>DYDNWAAQGNPGWSYP<br>DVLPLFKRAENNEDFGGD<br>DFHGSGGPLNVCYPRHAS<br>PINQMFIDAAAQNGLPQV<br>QDYNGATQEGAFLYQVT<br>HKNGERCSAAKGYLTPHL<br>GRPNLHVVTNAVSSRILLE<br>EGRAAGVEYHHDGQLKQ<br>VRARAEVVLSGGAFGSPQ<br>LLMLSGIGPAAQLQALGIP<br>VVRDLPGVGRNLQDHLD<br>HVQGWKTRSDTETFGLSP<br>RGVGKIAAAMMEWRSKR<br>TGMITSPYAEAGAFLRSAP<br>DVQVPDLQLVFVLALVD<br>DHARKMHLGHGISCHVD<br>LLRPHSRGEVTLQSRDAR<br>DAPVIDPRFLSDERDVETL<br>YRGVCLQQKIMESAPLDP<br>VRGKMLYPVRRGDRAATI<br>ADIRNRADTQYHPVGTCK<br>MGPRDDALAVVDAQLRV<br>RGVTGLRVADASIMPTIVS<br>GNTNAPSIMIGEKCAELLK<br>G |
| pZE-PcAAO<br>(WP_15804018<br>0.1) | ATGACGAACCAACAGGCCTTTGATTATATTATTGCGGGCGGTGGAACAGCA<br>GGGAGTGTGCTGGCGGGCAGATTAAGCGAGGATCCCTCAGTCAGAGTACTG<br>CTGTTAGAAGCAGGTCGCTCGGACCGCCATCCATTTATTCATGTTCCTGCAG<br>GGTTTGCGAAACTTACTGCCGGCCCGTTTCAGTGGGGATACAGCTCAGTGCC<br>ACAGGAACACGGCAACGGCAGGAGAATACCGCTGGCCCAGGGCCGGGTACT<br>TGGAGGAGGTGGCAGTATAAATGCACAAGTCTTCACGCGTGGTGTGGATGG<br>TGACTATGATGGTTGGGCAAACGAATATGGTGCCGAGGGCTGGAGTGCAGC<br>CGATGTTCGTCCGTATTTTGTACGCAGTGAACGTAATAACCGACTTGCAGGT<br>CCTCGGCATGGGACGGAAGGTCCGCTTGGCGTTAGCGATCTTGCATCGCCTC<br>ACCCGTTATCCCGGGATTTTGTGAAAGCAGGCCAGGAATTTGGTCTGCCGTA<br>TAGTAATGATTCCAATGGTGACCGCCAGGAAGGAATTGGCTTCTATCAAACT<br>ACGACGTATAATGGGCGTCGTTGCAGTGCAGCGGTTGCATACTTGGGGGGC<br>GGTGCCCGCAAACGACCTAATCTTGTTGTACGTACGCACGTAACGGTGTCCA<br>GGATAATTGTAACCGACGGGCGGGCTACTGGCGTAGAAGTTATCGAAAACG<br>GTACGCGCCATCGTTGCCTGGCACGTCGCGAGGTACTGGTGGCGTCGGGGG<br>CTTTTGGATCTCCGAAATTGCTGCAGCTGAGCGGCATTGGGGACCCCGCTGA<br>TTTAGCAGCGGCGGGCGTTGAAACGGTGCATGCGCTCCCCGGAGTTGGGAA<br>AAATCTCCAGGATCACTGTGATCTGGATATCATAGACGAGTTGAAAGACTAT<br>GCGTCTTTGGATAGGCTGAACCGAGTGCGACCAGCCACGGCGATTGCGGGA<br>CTTGAGTATCTGGCGTTTCGGTCGGGCCCGCTGGCTAGTACTGTCGTGGAAG<br>CCGGTGGGTTCAGCTTTGGGGATCCGAATGAAGCTACCCCGGATCTGCAGTT<br>TCATTTTTTACCTGCGGCCGGTGTTGAGGCCGGGATAGGAAGTGTGCGGCCA<br>GGATATGGGGCTACACTAAACTCTTGCTTCGTTCGTCCGCGGAGTCGTGGTA<br>CCGTTCGCATCGCATCCGCTGATCCTACCGTGGCTCCACTGATTGATCCAAA<br>TTATCTGGCAGATGAGCGCGATTTGGAAATGGCAGTTGTGGGAGTTGAGCA<br>GTCACGAGAAATCATGGCCCAACCTTCTATGGCTCGTAATATCAAAAAAGA<br>GCACATTGGTGATGGCAGTCCAGTTCGTACTAAAGACGAACTGGTCAGGTTC<br>GTGCGGAATTTCGGTAGAACCAGTTATCACCCAGTAGGTACGTGCGCCATGG<br>GTGTCAGCGATGATAGTGTGGTTTCACCCCGCTTGCAGGTGCATGGGTTGGA<br>AGGTCTGCGAGTTATCGATTCCAGTATTATGCCTAGGATCGTTAGCTCAAAT<br>ACTCAGGCACCAACTGTAATGATTGCGGAAAAAGCGGTTGACATGATAAGA<br>GAAGATGCGCGA | MTNQQAFDYIIAGGGTAG<br>SVLAGRLSEDPSVRVLLLE<br>AGRSDRHPFIHVPAGFAK<br>LTAGPFQWGYSSVPQEHG<br>NGRRIPLAQGRVLGGGGS<br>INAQVFTRGVDGDYDGW<br>ANEYGAEGWSAADVRPY<br>FVRSERNNRLAGPRHGTE<br>GPLGVSDLASPHPLSRDFV<br>KAGQEFGLPYSNDSNGDR<br>QEGIGFYQTTTYNGRRCS<br>AAVAYLGGGARKRPNLV<br>VRTHVTVSRIIVTDGRATG<br>VEVIENGTRHRCLARREV<br>LVASGAFGSPKLLQLSGIG<br>DPADLAAAGVETVHALP<br>GVGKNLQDHCDLDIIDEL<br>KDYASLDRLNRVRPATAI<br>AGLEYLAFRSGPLASTVV<br>EAGGFSFGDPNEATPDLQ<br>FHFLPAAGVEAGIGSVRP<br>GYGATLNSCFVRPRSRGT<br>VRIASADPTVAPLIDPNYL<br>ADERDLEMAVVGVEQSR<br>EIMAQPSMARNIKKEHIG<br>DGSPVRTKDELVRFVRNF<br>GRTSYHPVGTCAMGVSD<br>DSVVSPRLQVHGLEGLRV<br>IDSSIMPRIVSSNTQAPTV<br>MIAEKAVDMIREDAR |
| pZE-CoAAO<br>(MNZ24256.1) | ATGACAACAGATCCTATTAACAGCACCTTTGATTATCTGGTAGTGGGGGGTG<br>GTAGTGCTGGGTGTGTGATTGCCAGCCGTCTTTCGGAAGACCCTTCAATAAG<br>CGTGTGCCTGATTGAAGCCGGAGGCAGTGATTCCTCTGTGCTGATTCATGCG<br>CCTGTGGGCATGGTCGCGATGCTACCGACGAGGATTAACAACTGGGCCTTTC<br>AGACAGAGGTGCAACCTGGCTTAAACGGTCGCCGGGGCTACCAGCCCCGCG<br>GTAAAACACTTGGCGGCAGCTCTTCAATTAACGCCATGTTATATGTTCGGGG<br>GCACCGTCATGATTACGATCATTGGGCGGCCCAAGGTAATCAGGGGTGGAG<br>TTATGCTGATGTACTGCCGTATTTTATAAAAAGCGAGCAAAATCACTCCTTG<br>CACGACGATTATCACGGCCAGCACGGACCGTTGTCGGTTACAGAAGTCTCAT<br>GTCCGTCTGCTTTGAACGCTGCCTTCATACAAGCAGCACAATTGAATGGTAT | MTTDPINSTFDYLVVGGG<br>SAGCVIASRLSEDPSISVCL<br>IEAGGSDSSVLIHAPVGM<br>VAMLPTRINNWAFQTEVQ<br>PGLNGRRGYQPRGKTLGG<br>SSSINAMLYVRGHRHDYD<br>HWAAQGNQGWSYADVL<br>PYFIKSEQNHSLHDDYHG<br>QHGPLSVTEVSCPSALNA<br>AFIQAAQLNGIQRTADYN |

| | | |
|---|---|---|
| | CCAGCGGACAGCCGATTATAATGGCGCAGAACAGGAGGGTGCATTCATGTA<br>TCAGGTCACTCAGCGTAATGGGGAACGTTGCTCAGCGGCCAAAGCCTACCT<br>GACGCCGAATTTACAACGACGCAATCTTAAAGTGCTGACTGGTGCCACGGT<br>GGAGCGGGTCCTGATTGAGTCTGCGAGAGCGGTGGGCGTACAGTTGCGTCA<br>TGGAGGTCAACGCGTTGTGTTAAAAGCGAGGCATGAAGTGATTCTGAGTGC<br>CGGCAGCTTCGGTTCGCCCCAGGTGCTTATGCTGTCAGGTATCGGCCCTCAT<br>CAGGAATTACATAAACATGGCATCCGTATTACTCACGAGTTACCTGGCGTGG<br>GCCAGAACCTTCAAGATCATATTGATTACGTTTTCACTTACCGGTCTTCCAAC<br>AACACCCAGACATTCGGTTTTCGCCGACTTTCAGCTGGCAAATGCTTAAAG<br>CAATGGCCCAATGGAAAAGGGAGCGAAAAGGTTTGGTAACCTCACCTTTCG<br>CGGAGTCGGGAGCTTTTTTAAGAGTCAGCCCGGTTTATCGATGCCGGACCT<br>CCAGCTGATATTTGTTCCTGCAATTGTTGATGATCATGCACGTAAATTACGTC<br>CTTGGCATGGTTTCTCCTGCCATTTAACTCTGCTGCGCCCGAAATCTCGCGGG<br>ACAGTAAAACTTGCCTCCTCCGACCCTTCCGCAGCCCCATTAATCGATCCTC<br>GTTTCTTCTCGGAACCGCAAGACTTGGAGGTGCTAATGCGCGGTGCAGATAT<br>TCAGCGTGCAATCCTGGAAGCTTCTCCTCTCGCACCCTACAGAGGGAAACCG<br>CTTTATCCGCTGGACCAGCATGATCGTCAAGCCCTTGAGCAGGACATTCGCA<br>ACCGTGCGGACACCCAGTATCACCCAGTGGGTTCATGTAAAATGGGTCACG<br>ATCCACTGGCAGTAGTCGATGATCAGCTGCGCGTGCATGGAATCGCTGGTTT<br>AAGAGTCGCGGATGCCAGCATTATGCCCACACTTATCGGCGGTAACACTAAT<br>GCCCCTTCAATCATGATTGGAGAGAAAGCCGCCGATATGCTGTTAAGATCTC<br>GTCAGGAAGGT | GAEQEGAFMYQVTQRNG<br>ERCSAAKAYLTPNLQRRN<br>LKVLTGATVERVLIESAR<br>AVGVQLRHGGQRVVLKA<br>RHEVILSAGSFGSPQVLML<br>SGIGPHQELHKHGIRITHE<br>LPGVGQNLQDHIDYVFTY<br>RSSNNTQTFGFSPTFSWQ<br>MLKAMAQWKRERKGLV<br>TSPFAESGAFFKSQPGLSM<br>PDLQLIFVPAIVDDHARKL<br>RPWHGFSCHLTLLRPKSR<br>GTVKLASSDPSAAPLIDPR<br>FFSEPQDLEVLMRGADIQ<br>RAILEASPLAPYRGKPLYP<br>LDQHDRQALEQDIRNRAD<br>TQYHPVGSCKMGHDPLA<br>VVDDQLRVHGIAGLRVA<br>DASIMPTLIGGNTNAPSIM<br>IGEKAADMLLRSRQEG |
| pZE-VmAAO<br>(WP_09644139<br>3.1) | ATGGATAGTTACGATTTTATAATCGTTGGCGGGGGTTCGGCCGGCTGCGTAC<br>TCGCAAGCCGCCTGTCCGAAGATCCGACCGTAAATGTTTGTCTTCTGGAGGC<br>TGGTGGCAAGGACACTTCTCCGTTTATTCATACCCCGGTTGGATGTGTCGTT<br>ATGATGCCCACCAAAATAAATAACTGGGGGTTTGAGACTGTGCCGCAGCCT<br>GGTCTGAACGGCCGTAAAGGATATCAGCCACGTGGCAAAACTCTCGGGGGA<br>TCATCGTCGATCAATGCGATGATGTACGCTCGTGGCCATCGTTATGATTACG<br>ATCTTTGGGCGAGCCTTGGTAACGAAGGTTGGTCCTATGATGAGTGCCTACC<br>ATATTTTAAAAAAGCGGAAAATAATGAAGTCCACCATGATGAGTTTCATGGT<br>CAAGGCGGGCCATTGAACGTTGCAGATCTGCGTTCACCATCTCCTATGGTCG<br>AACGTTACCTGAGTGCGTGCGAGTCCATAGGTGTGCCGACGAACCATGATGT<br>GAATGGCGCAGAACAATTCGGCGCCATGCAGACACAGGTAACCCAGTTGAA<br>CGGAGAAAGGTGTTCCGCGGCGAAAGCGTATTTGACCCCTAATCTTAATCGT<br>CCTAATCTTACCGTCTTAACAAAGGCGACGACCCATAAAGTCCTTTTTGATG<br>GAAAACGTGCGATTGGCGTTGAATATGGGATGAAAGGTCAGAGATTCCAAA<br>TATACTGCAACAAAGAAGTTATCTTGAGCGCAGGTGCATTCGGGACTCCTCA<br>AGTGCTGCTACTGAGCGGTGTCGGCCCGAAACAAGAATTAGACAAACATGG<br>GATCGATCAGGTCCACGAATTAGCCGGTGTGGGTAAGAATTTGCAGGATCA<br>TATAGATTTAGTTCATTCCTACAGAACTACCGCCAAACGCGATACGTTTGGC<br>GTGTCCCTGAAAATGGCCTCTGAGGCAAGCAAAGCCGTTCCTCAGTGGTTTA<br>AACAGAGGCAGGGCAAGTTGTCATCCAACTTTGCAGAGGGGATAGGGTTTT<br>TATATAGCGATGACGATGTCGACGTACCGGATCTGGAGTTCGTCTTTGTTGT<br>CGCCGTGGTCGACGATCACGCGCGTAAGATTCATATGTCTCATGGCTTTAGC<br>AGCCATGTTACTTTGCTACGCCCGAAGTCCACGGGAACAGTCACCCTGAACA<br>GTGCAGATCCGTATGATGTGCCTTCAATTGATCCCGCTTTTTTTCAGGATCCT<br>GACGATATGCGTGTAATGATTAAAGGGTGGAAAAAACAGTACCAAATGCTA<br>CAGTCCGAAGCTTTTGACGATGTGCGCGGGGCATCGTTTTACCCAGTCGATC<br>CCGATGACGACGCCGCTATCGAACAGGATATTCGTAACAGAGCTGACACTC<br>AGTACCACCCGGTTGGCACATGTAAAATGGGGACGTCCGACGATCTGGAAG<br>CTGTTGTTGATAGTGAACTTTCCGTTTATGGCATGGATAACTTGCGGGTTGTT<br>GATGCCTCCGTAATGCCGACTCTAGTAGGAGGTAATACCAACGCCCCTACCA<br>TAATGATTGCTGAAAAAATTGCTGATGTTATAAAAGCGAAATATTCCATATG<br>CCATCCAGCAGATGCCTCCAAAGTGTCT | MDSYDFIIVGGGSAGCVL<br>ASRLSEDPTVNVCLLEAG<br>GKDTSPFIHTPVGCVVMM<br>PTKINNWGFETVPQPGLN<br>GRKGYQPRGKTLGGSSSI<br>NAMMYARGHRYDYDLW<br>ASLGNEGWSYDECLPYFK<br>KAENNEVHHDEFHGQGG<br>PLNVADLRSPSPMVERYL<br>SACESIGVPTNHDVNGAE<br>QFGAMQTQVTQLNGERC<br>SAAKAYLTPNLNRPNLTV<br>LTKATTHKVLFDGKRAIG<br>VEYGMKGQRFQIYCNKE<br>VILSAGAFGTPQVLLLSGV<br>GPKQELDKHGIDQVHELA<br>GVGKNLQDHIDLVHSYRT<br>TAKRDTFGVSLKMASEAS<br>KAVPQWFKQRQGKLSSN<br>FAEGIGFLYSDDDVDVPD<br>LEFVFVVAVVDDHARKIH<br>MSHGFSSHVTLLRPKSTG<br>TVTLNSADPYDVPSIDPAF<br>FQDPDDMRVMIKGWKKQ<br>YQMLQSEAFDDVRGASF<br>YPVDPDDDAAIEQDIRNR<br>ADTQYHPVGTCKMGTSD<br>DLEAVVDSELSVYGMDN<br>LRVVDASVMPTLVGGNT<br>NAPTIMIAEKIADVIKAKY<br>SICHPADASKVS |
| pZE-NnAAO<br>(WP_07238075<br>4.1) | ATGTCCCAAGGTGAGTTTGATTATATTGTTATTGGAGCCGGGTCCGCGGGGG<br>CCGTCCTGGCAGCTCGTTTGTCCGAGGATCCGCACATGAGGGTTCTGCTGTT<br>GGAAGCGGGCGGCCCAAATACGTCTGTTTTGGTAAGAATGCCCGCCGGCGT<br>CGGAACATTAATTAAACAAAAATCTAAACATAATTGGGGATTTTGGAGCGA<br>ACCCGAGCCCCACATGGACGGCCGTCGCATGTGGCATCCTCGGGGTAAAGG<br>CCTGGGAGGTTCTTCCGCAATTAACGGTATGGTGTACGTCCGTGGGCATGCC<br>ACAGACTATGATCAATGGCGCCAGTTAGGCCTTGAAGGCTGGTCCTATGCCG<br>ATGTGCTGCCCTACTTTCGTAAAGCGGAAGATAATTGTCTTGGTGCAAGCGA<br>ATTTCATGGCACCGGGGGTCCCTTAAAAGTTAACTGGGGTGAACATTCAGAA<br>CATCCATTGTATCAGTCATTCCTTGCTGCGGGCCAGCAAGCAGGCCATCCAT<br>TTACTGAGGACTTCAACGGCGCCAGCCAAGAAGGGTTTGGCCGTTATCAACT<br>GACGATTAACGATGGTCAGCGTTGGTCCGCAGCACGAGGTTATTTGACGCCT<br>ATTATGGGCACACGTCGTAATCTTGCGGTGGTCACAGGTGCCCGTGTGCACA<br>GGATTGTGATCGCCAATGGTCGAGCGGCAGGTGTTGAATATAGCCTCGGAC<br>CGGGAAAGGCCGTACAGACTGCGCAGAGTTCAAGAGAGGTTTTGCTGTGTG<br>CAGGGGCATTCCAGAGTCCGCAGATTCTTCAACTATCTGGCATTGGGGACCC | MSQGEFDYIVIGAGSAGA<br>VLAARLSEDPHMRVLLLE<br>AGGPNTSVLVRMPAGVG<br>TLIKQKSKHNWGFWSEPE<br>PHMDGRRMWHPRGKGL<br>GGSSAINGMVYVRGHAT<br>DYDQWRQLGLEGWSYAD<br>VLPYFRKAEDNCLGASEF<br>HGTGGPLKVNWGEHSEH<br>PLYQSFLAAGQQAGHPFT<br>EDFNGASQEGFGRYQLTI<br>NDGQRWSAARGYLTPIM<br>GTRRNLAVVTGARVHRIV<br>IANGRAAGVEYSLGPGKA<br>VQTAQSSREVLLCAGAFQ<br>SPQILQLSGIGDPEKLKAA |

| | | |
|---|---|---|
| | GGAAAAACTGAAAGCAGCCGGGGTGGCGCCGGTCCATGCCCTGCCGGGCGT CGGAGCTAATCTGCAAGATCATCTGGACGTAACACTAAACTGGGCAGCTAC CCAGCCAGTCAGTCTGTATCATCAAATTCGGGGATGGCGACAGTATAAAGTC GGTCTTCACTATCTACTTACTGGCAAAGGCGGAGGCCGCCAGAATGGTCTGG AAGCAGGCGCGTTCCTGAAATCACGTCCGGATCTCGACCGCCCTGACTTGCA GCTCCATTTCGTAATCGCTATAATGCAGGAACATGCAAAAGTACGTGTTGAG CGTGACGGCTTCACAGTGCACGTGTGCCAGTTACGCCCTGAGTCACGCGGTA CTGTTGGACTGCGGTCCCGTGACCCCTATGATGATCCTGCCATTACTGCGAA CTTTCTGGCAACCGAAGAGGACCGGAGAGTCGTTAGAGAGGGTATCCGGAT TGCCCGAGAGGTGGTGGCACAGGACGCATTTGCTCCTTATCGTGGCGAAGA GATTTGGCCGGGTGCTGAAGTGCAATCTGACGCTGCGATTGATGCCTGGGTT AGAGCCCGAGGGGAAACCATTTACCATCCCGTTGGCACTGCAAAGATGGGA ACAGGCACGGATCCCATGGCTGTCGTGGATAAGGATTGCAAGGTGATAGGT CTTGCTGGCCTGCGTGTCGTAGATGCAAGCGTTATTCCTCTGTTGGTTGGCG GTAACACGAATGCACCTACAATTATGATCGCTGAAAAAATCGCCGATGTGA TTCGCGGCCCGAGTGCCTTAGCTGCTGCAGCT | GVAPVHALPGVGANLQD HLDVTLNWAATQPVSLY HQIRGWRQYKVGLHYLL TGKGGGRQNGLEAGAFL KSRPDLDRPDLQLHFVIAI MQEHAKVRVERDGFTVH VCQLRPESRGTVGLRSRD PYDDPAITANFLATEEDRR VVREGIRIAREVVAQDAF APYRGEEIWPGAEVQSDA AIDAWVRARGETIYHPVG TAKMGTGTDPMAVVDKD CKVIGLAGLRVVDASVIPL LVGGNTNAPTIMIAEKIAD VIRGPSALAAAA |
| pZE-PclAAO (WP_045732460.1) | ATGCACTATGATAATATAGATGATCTTTCAGATCGAGCATTTGACTACGTTG TGATTGGTGGGGGATCAGCGGGAGCCGCAGTAGCAGCTCGGCTCAGCGAGG ACCCAGATGTGACTGTTGCCTTAGTAGAGGCTGGTCCGGATGACCGCAATAT TCCCGAGATCCTGCAGCTGGACCGTTGGATGGAGCTGCTGGAGAGCGGCTA TGATTGGGACTATCCTATTGAGCCCCAGGAAAATGGAAATTCATTTATGCGT CATGCGAGGGCGAAAGTGATGGGGGGTTGTAGTTCACATAATAGCTGCATC GCTTTTTGGGCACCGAGGGAGGACATTGATGAGTGGGAGTCTAAATATGGC GCTGCGGGCTGGAACGCAGCTAATGCATGGCCGTTATACAAGCGTCTTGAA ACGAACGAAGATGCAGGCCCGGACGCGCCTCATCATGGGGACTCCGGGCCC GTTCACTTAATGAATGTACCGCCGGCGGATCCTTCAGGTGTGGCGCTGCTGG ATGCTTGCGAGCAGGCTGGTATACCTCGTGCAAAGTTTAACACTGGAGCGAC CGTAGTTAATGGGGCCAATTTCTTCCAGATCAATCGCCGTGCGGATGGCACA AGGTCGAGCAGCTCAGTATCTTATATACATCCGATAGTTGACCGGCCCAACT TTACTCTACTGACAGGTCTGAGAGCGAGACAGTTAGTAATCGACGCGGATC GACGGTGTACTGGCGTGGACGTAGTAGACGGCGCGTTCGGAAGAACGCATC GGATTACTGCACATCGGGAGGTAATAGTAAGTACTGGAGCCATTGACTCTCC AAAATTACTCATGTTGTCGGGGATAGGCCCTGCTGATCATTTGGCGGCACAC GGAATTGAAGTCCTGGTAGATAGTCCGGGTGTTGGTGAGAATCTGCAAGAC CACCCGGAAGGTGTCGTTCAATTTGAGGCCAAACAACCTATGGTAGAGACA AGTACACAATGGTGGGAAATAGGCATATTTACCCCTACAGAAGATGGCCTT GATAGACCGGATTTAATGATGCATTATGGATCCGTGCCATTTGATATGAACA CATTGCGTTATGGCTATCCGACCACAGAAAATGGATTTTCTCTCACACCTAA CGTCACCCACGCCCGTTCACGCGGCACCGTTCGCCTGCGCTCACGTGACTTT CGAGATAAGCCAATGGTCGATCCAAGATATTTTACAGACCCCGAGGGACAC GACATGCGGGTGATGGTGGCGGGGATCCGGAAGGCGCGCGAAATTGCCGCA CAACCGGCAATGGCGGAGTGGACCGGCCGCGAACTTAGCCCAGGTGTGGGA GCGCAGACTGATGAAGAACTGCAGGATTATATTCGTAAAACACATAATACT GTGTATCATCCTGTAGGAACCGTTCGCATGGGAGCGGACGATGATGGTATGT CTCCCTTAGATGCAAGACTGCGTGTCAAAGGTGTCACAGGTCTGCGAGTGGC CGATGCTTCAGTTATGCCGGAGCACGTAACCGTCAATCCGAACATTACAGTG ATGATGATCGGTGAACGTTGCGCAGATCTGATCAAAGCGGATTACGCAGGC GCCGATGCTCTGGAAGAAAAAGAATTGACCACGAGTTTTGCG | MHYDNIDDLSDRAFDYV VIGGGSAGAAVAARLSED PDVTVALVEAGPDDRNIP EILQLDRWMELLESGYD WDYPIEPQENGNSFMRHA RAKVMGGCSSHNSCIAFW APREDIDEWESKYGAAG WNAANAWPLYKRLETNE DAGPDAPHHGDSGPVHL MNVPPADPSGVALLDACE QAGIPRAKFNTGATVVNG ANFFQINRRADGTRSSSSV SYIHPIVDRPNFTLLTGLR ARQLVIDADRRCTGVDVV DGAFGRTHRITAHREVIVS TGAIDSPKLLMLSGIGPAD HLAAHGIEVLVDSPGVGE NLQDHPEGVVQFEAKQP MVETSTQWWEIGIFTPTE DGLDRPDLMMHYGSVPF DMNTLRYGYPTTENGFSL TPNVTHARSRGTVRLRSR DFRDKPMVDPRYFTDPEG HDMRVMVAGIRKAREIA AQPAMAEWTGRELSPGV GAQTDEELQDYIRKTHNT VYHPVGTVRMGADDDG MSPLDARLRVKGVTGLR VADASVMPEHVTVNPNIT VMMIGERCADLIKADYA GADALEEKELTTSFA |
| pZE-PrAAO (WP_065947574.1) | ATGAAAAAAACGTTCGATTTTATCGTTGTAGGTGGTGGGTCCGCGGGCTGTG TGGCCGCAGGGCGTCTTTCGGAAGACCCTGATACCAGCGTATGTCTGCTGGA GGCAGGGGGCGAAGGTCGAAGCTCACTTGTTAGAATTCCAGCAGCTACCGT GGCAATGGTCCCCACGAAAGTAAATAACTGGGCCTTTGATACAGTGGCCCA AGCTGCTCTGCTGGGTCGCACGGGTTATCAGCCTAGAGGAAAAACCCTGGG TGGCTCATCTAGCATTAATGCGATGATATACGTAAGAGGTCATCAGTGGGAT TATGACCACTGGGCGTCACTTGGAAACCCAGGTTGGGGTTACAAAGATGTCT TGCCCTATTTTCTGCGCTCTGAACATAATGAACGTATAGATGACGCGTGGCA TGGCAGAGACGGCCCGCTGTGGGTTAGCGATCTGAGATCGGATAACCCTTTT CAGCAACGTTTCCTAGAAGCAGCAAGAGAAACCGGGCTCCCTCTTAATGAC GACTTTAATGGAGCAGAACAGGAAGGCGTAGGTGCGTACCAAGTTACCCAG AAACATGGTGAAAGATACTCCGCCGCACGTGCTTATCTGCTGCCACATATAG GTGTTAGAGATAATTTATCTGTCGAAACCCGGGCGCAGGTGCAGCGTATTTT ATTTGAGGGTACACGGGCAGTGGGTGTTGAGGTGCTTCAACATGGGCAAGT CTATGTTCTAAGAGCACGGCGCGAGGTAATCCTGGCCGCGGGTGCGTTCCAG ACACCGCAGCTGCTCATGTTGAGTGGCGTCGGGCCGAAGGTTGAACTTCAG AGACATGGTATCCCTCTACTACATGAATTGCCTGGCGTCGGCCAAAATCTCC AGGATCATCCGGATTTCGTATTTGTATACAAAACTAACTCTCTTGATGCAAT GGGGGTTAGTCTGGGAGGATGTCTGAAAATTTTAAAAGAAATTTGGAGATT CCGCCAGGAACGCCGGGGTATGCTTACATCGAATTTTGCAGAAGGGGGCGC GTTCCTAAAAACTTGCGATACATTAGATAAACCGGACATTCAGCTTCATTTC GTTGTTGCGCCAGTCGAAGACCATGCGCGTACTTTACGTATGGGGCACGGCC | MKKTFDFIVVGGGSAGCV AAGRLSEDPDTSVCLLEA GGEGRSSLVRIPAATVAM VPTKVNNWAFDTVAQAA LLGRTGYQPRGKTLGGSS SINAMIYVRGHQWDYDH WASLGNPGWGYKDVLPY FLRSEHNERIDDAWHGRD GPLWVSDLRSDNPFQQRF LEAARETGLPLNDDFNGA EQEGVGAYQVTQKHGER YSAARAYLLPHIGVRDNL SVETRAQVQRILFEGTRA VGVEVLQHGQVYVLRAR REVILAAGAFQTPQLLML SGVGPKVELQRHGIPLLH ELPGVGQNLQDHPDFVFV YKTNSLDAMGVSLGGCL KILKEIWRFRQERRGMLT SNFAEGGAFLKTCDTLDK PDIQLHFVVAPVEDHART LRMGHGLSCHVCLLRPRS |

| | | |
|---|---|---|
| | TATCGTGCCATGTATGCCTCTTGCGTCCACGTTCTCGAGGTTCAGTTACTCTTGCCTCTAATGATCCACAGGCGGCCCCATTGATCGATCCCGCATTCCTGAAAGATCCACAGGACCTGGAAGATATGGTTGCAGCGTTTAAACTTACGCGTCGCCTCATGCAGGCGCCGAGTCTGGCCAAATGGATCACCCGTACTCTTTATACTGAAGGAGTTGAAACGGATGAGCAGATCAGGACACTACTGCGCGAAAGAACCGATTCGGTTTATCATCCGGTGGGAACTTGCAGAATGGGCGACGACCCGCTGGCAGTTGTTGACGCCCAGCTTCGGGTTCATGGTCTGCAAGCACTTCGTATTGTCGACGCGAGCATAATGCCGACTCTCATCGGCGGTAACACGAATGCGCCGACCATCATGATAGCCGAGAAGGCGGTCGACCTGATACGTGGCTTGCCACAAACTCATTCAAGTGCACGTTTTGAGAATGCGGTTGCTGCCAAGGAAGTTCCGCACGTTACCGCA | RGSVTLASNDPQAAPLIDPAFLKDPQDLEDMVAAFKLTRRLMQAPSLAKWITRTLYTEGVETDEQIRTLLRERTDSVYHPVGTCRMGDDPLAVVDAQLRVHGLQALRIVDASIMPTLIGGNTNAPTIMIAEKAVDLIRGLPQTHSSARFENAVAAKEVPHVTA |
| pZE-SpAAO (WP_114226814.1) | ATGGCGGAAGCGGAACGAACGGCTGATTACGTTATTGTGGGTGCGGGTAGTGCCGGTTGCGTTCTGGCTGCACGTCTGTCAGAAGATCCGGCGGTGAGCGTTGTGTTGCTGGAAGCCGGCGGTGATGACCGACCACTGAAAGAGCCTAGTCAGCTGATTTCAAATATGTGGATCCATGTTCCGCTCGGTTATTCTCAGACCTTAAAAGATCCTAAAGTGAATTGGCTCTACGTGACCGAACCAGATCCGGGCACCCATGGTCGCCAACATGTGTGGCCAAGAGGTAAAGTGTTAGGTGGTAGTTCCAGCATTAATGCAATGCTCTACGTTCGTGGTCAGCGCAGTGATTATGATCATTGGCGCCAGCTTGGTTGTGAAGGGTGGGGTTGGGATGAAGTGTTTCCCTATTTCCGTAAAGCACAACATCAGGAACGAGGCCCATGTGATGATCATGCCGTAGGGGGTCCACTTAACGTCTCTGATCCACGCGATGGCCACGCAGTTTCAGAAGCCGCTATCGCAGCGTGTGTCGCCCTAGGGTTGCCGGAACGTGACCTGAATGGCCCAGAACAGGAGGGTGTGGGCTGGTTTCAATTAAATATCAAGGATGGCAAGCGGTGTTCAGCGGCTGTGGCCTATTTGCACCCAGCAATGGGGCGTGCCAATCTCACCGTTGAAACTCGTGCTCTGGCGTCGTTAGTTCGCTTTGAGGGCAAACGGGCGACCGGCGTCGAATTTACTCAGCGGGGTGTGCGCCGTGTTGTACATGCTAGACGGGAAGTAATTCTTGCTGGAGGGGCTATTAACAGTCCGCAGCTCCTGGAACTGTCCGGTGTTGGCCAAGCAGAGCGTCTGAGAGGGTTAGGTATCGAAGTGGTACATGAACTTGCTGGTGTTGGTGAAAACCTGCAAGATCATTATGTGGTTGGGGAAACGTACAGACTTAAGGCAGGTACTATTTCCGTAAATGAATTAGCTAGAGGACCGAGACTCCTGCGTGAAGTTGTACGTTATTTAGAGAAAAACGCGGTCTTCTGACCCTGAGTGCGGCTCACGTGGGGGTTTTCTGTAAAAGTCGGCCAGACCTAGCCGGTCCCGATATCCAGTTCCACATCCTGCCGGCAACCATGGATGCAGAAAAATTGGCAAATGAGCAGAAAATGGAGCTCGAAGGTGAACCCGGACTGACCATCGCGCCCTGTCAGCTGCGACCGGAGTCGAGGGGTTCTGTGCATTTGCGATCCGGTGATCCGGCGGAACATCCAGCAATCGCACCGAATTATCTCTCTAATCCCCTGGATGAAGAAGTGGTTGTAGCAGCTTTAAAATGGGGTCGCCGTATCGCGGCCACAGCCCCTCTGGCTGGTTACATCGATCATGAAACGAACCCTGGCGCCGCAGTCGCTTCTGACGCCGCGCTCCTTGACTACGCCCGCGAGCGGGGAACGACCATCTATCATCCCGTTGGAACATGCGCCATGGGGCGTCATGCCGGCGCTGTTGTTGATCCAGAGTTACGCGTTCATGGTCTGGACGGTCTAAGAGTGGTGGACGCTTCGATAATGCCTCGGCTTGTGTCTGGTAATACCAATGCGCCGACAATTATGATTGCGGAGAAAGCAGCCGATATGATCAAGGCCTCGGCGAAAGCTAGAGTCGCCGCG | MAEAERTADYVIVGAGSAGCVLAARLSEDPAVSVVLLEAGGDDRPLKEPSQLISNMWIHVPLGYSQTLKDPKVNWLYVTEPDPGTHGRQHVWPRGKVLGGSSSINAMLYVRGQRSDYDHWRQLGCEGWGWDEVFPYFRKAQHQERGPCDDHAVGGPLNVSDPRDGHAVSEAAIAACVALGLPERDLNGPEQEGVGWFQLNIKDGKRCSAAVAYLHPAMGRANLTVETRALASLVRFEGKRATGVEFTQRGVRRVVHARREVILAGGAINSPQLLELSGVGQAERLRGLGIEVVHELAGVGENLQDHYVVGETYRLKAGTISVNELARGPRLLREVVRYFREKRGLLTLSAAHVGVFCKSRPDLAGPDIQFHILPATMDAEKLANEQKMELEGEPGLTIAPCQLRPESRGSVHLRSGDPAEHPAIAPNYLSNPLDEEVVVAALKWGRRIAATAPLAGYIDHETNPGAAVASDAALLDYARERGTTIYHPVGTCAMGRHAGAVVDPELRVHGLDGLRVVDASIMPRLVSGNTNAPTIMIAEKAADMIKASAKARVAA |
| pZE-BuAAO (WP_057055115.1) | ATGAATTTCGACTACATTATCGTGGGGGGGGGTTCTGCCGGCGCAACCTTGGCATCACGTTTGAGTGAAGATTCATCTGTCAGTGTGTGCCTGCTGGAGGCCGGCGGTCAGGGTGATTCTCTGGTGGTACGAATGCCTGCAGGAACCGTCGCGATGATGCCTGGACGGGGAAAAATTAATAACTGGGCTCTCAACACCGTTCCACAGCCTAACCTAAATGGCCGTCGCGGGTATCAACCCCGTGGGCGTGCCCTGGGCGGGTCATCTGCAATTAATGCCATGTTGTATGTTAGAGGCCAACGTCAAGATTATGACCATTGGGCGGACCTGGGGTGTCAGGGATGGGATTGGGATAGTGTTCTTCCTTACTTTCTGAAAGCTGAGAATAATGAACGCGGTGCCAGCAAATTGCACGGTGGTAACGGCCCATTGCACGTTAGCGAGCAAAGAAGCCCCCGACCGGTGACACTGGCTTTCATCGAGGCAGCGAAACAACGAGGAATTCCTTTTAGGGCGGATTTCAACGACGGGGACAACGAAGGCGTGGGATTATATCAAGTAACCCAGTTCCACGATACCGCACGGCGTGGTGAACGCTGTAGTGCGGCGGCAGCCTATTTACATCCCCTCCGTGGGCAGAGGCCAAATCTGACCATTAAGACTAATACCCGTGCGTCTCGGATAATTTTCGAAGGTAAACGTGCAGTTGGTGTGGCCTATCATGCCTCGGGCGCAGAACACGAGGCACGTGCAAAACGAGAAGTCATACTGGCCGCGGGGGCGTTCGGGACCCCTCAGATATTACAGTTAAGTGGAGTCGGACGCCCGGAAGACATAACCCACCACGGCATACGTATGGTGCACGAACTCCCCGGTGTGGGTCAGAACCTCCAGGATCATTTAGACTTTATTCTCGCATACACGACCCGGGATGCAGACAACTTCGGAATAGGACTACGCGGGAGTGTAAATATGCTCGGCCATATTGCGTCATGGAGAAAAGATGGTTCGGGTATGCTTGCCACACCGTTTGCTGAAGGTGGAGCATTTTTCAAATCTGCCCCTGACGTTGATCGGCCGGATCTGCAGTTACATTTTGTTATTTCCATCGTGGATGACCATGCCCGCAAATTACATGTGGGGTATGGTTATTCCTGTCATGTTTGTAATCTCAGACCATATTCGCGCGGCGAAGTATTTCTGCAGTCCCCAAATCCTTTGGATGACCCCGGCATCGACCCGCGATTTTTAAGTGATGAACGTGACGTTAAGCTCTTAGTACAGGGCGCCAAAGCAACGCGCGAAATTATGTTAGCGTCACCGCTTGCGAAATACCGTGGACGC | MNFDYIIVGGGSAGATLASRLSEDSSVSVCLLEAGGQGDSLVVRMPAGTVAMMPGRGKINNWALNTVPQPNLNGRRGYQPRGRALGGSSAINAMLYVRGQRQDYDHWADLGCQGWDWDSVLPYFLKAENNERGASKLHGGNGPLHVSEQRSPRPVTLAFIEAAKQRGIPFRADFNDGDNEGVGLYQVTQFHDTARRGERCSAAAAYLHPLRGQRPNLTIKTNTRASRIIFEGKRAVGVAYHASGAEHEARAKREVILAAGAFGTPQILQLSGVGRPEDITHHGIRMVHELPGVGQNLQDHLDFILAYTTRDADNFGIGLRGSVNMLGHIASWRKDGSGMLATPFAEGGAFFKSAPDVDRPDLQLHFVISIVDDHARKLHVGYGYSCHVCNLRPYSRGEVFLQSPNPLDDPGIDPRFLSDERDVKLLVQGAKATREIMLASPLAKYRGRELFGVTDGMSDAQWGQL |

| | | |
|---|---|---|
| | GAATTATTTGGAGTGACCGATGGCATGAGTGACGCACAGTGGGGACAACTT<br>ATACGGGATCGCTCTGACACGATTTATCATCCGGTTGGTACCTGTAAAATGG<br>GGGTCGATCCGATGGCGGTCGTTGACGCGGAATTAAAAGTACACGGCTTGC<br>AGGGGCTCAGAGTGGTAGACGCCAGCATTATGCCGACCTTAATAAGTGGTA<br>ATACCAATGCGCCTACCATCATGATCGCTGAACGTGCTGCGGATCTGATTAA<br>GCGTGCTCACGGTCATCCGGCGCAGATAAGTAGAAAAAATGAACGACAAGC<br>TTACGAATCAGCCCGTCTTCGCGGTTAA | IRDRSDTIYHPVGTCKMG<br>VDPMAVVDAELKVHGLQ<br>GLRVVDASIMPTLISGNTN<br>APTIMIAERAADLIKRAHG<br>HPAQISRKNERQAYESAR<br>LR |
| pZE-AtAAO (GFF15729.1) | ATGGCGGGCCCTGCCAGTCAGTTGGCAGCAAATGCTATCTGTAGTCTTGAAG<br>AATTCCGTCGTGTGAGTTATGACTATATTATAGTAGGTGGTGGCACAGCCGG<br>ACTGACGTTAGCGGCTCGACTGAGCGAGGACCCGAACGTAAATGTTGGGGT<br>TCTGGAAGCTGGAAAAGATCAGACCAAAAATGAACTGGTTCGCACCCCGGC<br>TCTGTTCCCCCAAATGCTCACGAATCCCGAGTACGATTGGCTTATGTATACA<br>GTTCCGCAGAAAGGTAACCATAACAAAATTCACCATCAGACTCGAGGTAAG<br>ATGCTCGGCGGTTGTTCGGCCACGAATGGAATGATGTATGTAAGAGGTTCGA<br>AACAAGATTTTGATGACTGGGGTGCATTCGGCAAAGGCTGGAGTTGGTCCTC<br>CATCGCACCATACTTCCGTAAACACGAACGTATGGACGACACACGCGTCGG<br>CTTACCGGGAGATAACAAATTTTTACAATTCCAGAAAAAATCCCATGGTCAG<br>CACGGCCCTATCGAAACAAGCTTTAATAACTGGAGGAACCCACTGGAAAGA<br>TACTTCCTGCAGGCAGCAAAAGAAGCATCAGGTATGGCAGCGAGCCCGGTC<br>GACCCGTGGGGTGGGGATCATTTGGGATTTTTTTCCTCTCTGGCGACAGTTG<br>ATAGACGCGGTGATAAGGGTACCAGATCTTATGCTGCAACAGGTTATCTTCT<br>CCCAAACCTGACGCGTCCAAATTTGAAGGTGTTAACTGAAGCATTAGCCGTT<br>TGTGTGACGCTCGAAGGGAACTCAGCGAGCGGTGTTCGCTTTATGCATTCGG<br>GAACCACCTATGATGTACGTGCAGCGAAAGAGGTCATTATATCGGGGGGAG<br>TGTATAAATCTCCCCAGGTGCTGGAGCTTAGCGGAATTGGTGATCCAAGCGT<br>GTTGAAAGCTGCTGGTGTGCAGTGTAAAGTTCCCCTACCGGGCGTGGGCGCC<br>AATCTTCAGGATCATGTGCTCTCTGGTGCCGTTTATGAGTTGAAAGACGGGG<br>TAATGTCCTTCGATGCGCTTCGTAACCCTTCTGTCGCCCAGGAACATATGGA<br>TATGTATGCCAGGGACCGGACCGGTATTCTCGCAGCGGCGACTAGTTGCATG<br>GGGTTCTTACCTTATTCTAGTCTGGTGAGCAAGGAAGAGCTGGAAGCCACCT<br>GTCAGAAGGTTCTAAGCGTTCCGGCAGAAACCGCATTTCAGCAGAAACAGC<br>GGGAGCAAATTGTAAGCCATCTCCGCTCACCGAGCAGCGCCAATATTCAGT<br>ACATCCTTTCGCAGGGTACAGTAGACTTAGAAAATGCCCCTGGAGATCAGA<br>GCAAATTTGCGAAGGCGCTGTCTCCCGAGGATCCTAATGGGTTTACGATTGT<br>TACATGTCTGCAGTACCCCAGTTCTCGTGGTACTGTTCATATTACCAGTAGC<br>GATCCGCATCAGAATCCGGCCATTGACCCTGCCTATCTCACAAATCCCGCGG<br>ATGTCGATATTCTGGCCGCAGGGCTAGAGTTTTGTGATAAAATTGCCTCGGC<br>GCCGGGGTTGAAAGACAAGGTCGTTCGTCGGGCGCTACCAAGCCCAAGCGT<br>CTCCCTCCAGAGCCGATCACAGGCCGCCGAAGCCGTGCGTGAAAACTGCAT<br>GACGGAATATCACCCATGCGGTACTTGCGCAATTGGTCAGGTTGTCGATGAG<br>CGACTTCGCGTACTGGGGGTTAAACGGTTGCGGGTTGTCGATGCCTCGGTTT<br>TTCCTGGGAACGTCTCCGGTAACATATTATCAAGCGTGTACGCGGTGGCGGA<br>GAAAGCGGCAGACATGATTAAAGAAGATGCTAAACATGTGCGTGCCAGCCT<br>G | MAGPASQLAANAICSLEE<br>FRRVSYDYIIVGGGTAGLT<br>LAARLSEDPNVNVGVLEA<br>GKDQTKNELVRTPALFPQ<br>MLTNPEYDWLMYTVPQK<br>GNHNKIHHQTRGKMLGG<br>CSATNGMMYVRGSKQDF<br>DDWGAFGKGWSWSSIAP<br>YFRKHERMDDTRVGLPG<br>DNKFLQFQKKSHGQHGPI<br>ETSFNNWRNPLERYFLQA<br>AKEASGMAASPVDPWGG<br>DHLGFFSSLATVDRRGDK<br>GTRSYAATGYLLPNLTRP<br>NLKVLTEALAVCVTLEGN<br>SASGVRFMHSGTTYDVRA<br>AKEVIISGGVYKSPQVLEL<br>SGIGDPSVLKAAGVQCKV<br>PLPGVGANLQDHVLSGAV<br>YELKDGVMSFDALRNPSV<br>AQEHMDMYARDRTGILA<br>AATSCMGFLPYSSLVSKE<br>ELEATCQKVLSVPAETAF<br>QQKQREQIVSHLRSPSSAN<br>IQYILSQGTVDLENAPGDQ<br>SKFAKALSPEDPNGFTIVT<br>CLQYPSSRGTVHITSSDPH<br>QNPAIDPAYLTNPADVDIL<br>AAGLEFCDKIASAPGLKD<br>KVVRRALPSPSVSLQSRSQ<br>AAEAVRENCMTEYHPCG<br>TCAIGQVVDERLRVLGVK<br>RLRVVDASVFPGNVSGNI<br>LSSVYAVAEKAADMIKED<br>AKHVRASL |
| pZE-OaAAO (WP_083562132.1) | ATGTTCGACTATATTGTCGTGGGTGGTGGCTCTGCGGGAGCTGTGGTTGCCA<br>GTCGTCTATCTGAAGATCCGAATGTTAGTGTTTGTCTGCTCGAAAGTGGACC<br>GGTTGATAAATCAGTGTTGATTCACGCTCCTGCCGGCGTGGTTGCCATGATG<br>CCGAGAAAAAACAAACTTAACTATGCATTTGAGACGGTGCCACAGAAGGGG<br>TTAAATGGTCGTCGCGGCTATCAACCGCGTGGACGGTGTCTGGGCGGGAGTT<br>CATCTGTGAATGCAATGCTGTATGTGAGAGGTCATCGGGCTGACTATGACCA<br>CTGGTCAGCACTGGGCAACGCCGGATGGTCTTACGCAGAAGTTTTACCTTAT<br>TTTAAACGATCCGAACATAACGAGAACATTCACGACGAATTCCACGGCCAG<br>GGCGGACCGCTTAACGTGATGAATCTTCGCAGTCCAGGTGCATTAAACGCA<br>GAGTTCATTAAGGCGTGTGAGGAGCAGGGAATCGCAGCCACAGCGGATTGT<br>AACGGGGCGGAGCAAGATGGTGCACTGGAATATCAGGTCACACATATCAAT<br>GGGGAACGTTGTAGCGCGGCCAAAGCATATCTCACACCTAACTTGTCTCGCC<br>CGAATCTGGAAGTTATGACTCAGACGCCAGTGCTACAGATATTGCTGGACG<br>GTAAGACCGCGACTGGCGTTCGTGTGAAACGCGAAGGACAAACTCTGGATC<br>TCCGTGCACGTCGTGAGGTAATAGTTTGTGGCGGTGCATTCAATTCCCCCCA<br>GTTACTGATGCTGAGTGGAATTGGCCCTGGTGCTCATCTGCAAGATCGTGGC<br>ATAGCCGTGGCACATGATCTGCCGGGCGTAGGTGAAAACCTGCAGGACCAT<br>ATCGATATTGTACATTCATACCGCGCTCCGAATTCTGACACTTTCGGTCTTTC<br>GTTTGGATTCATGCCCAAAATTCTGGCCGCCATGTTACAGTGGCGCCGGCAA<br>CGCTCCGGACTTATTACATCGCCTTACGCAGAAGCAGGCGCCTTTTTCGCT<br>CATCTGAAGCTGACGGTCGGCCGGATTTACAGCTGATCCTGGTTCGAGCGTT<br>AGTGGACGACCATGGTCGGAAGATGCACCTGGGCCACGGATTTTCATCTCAC<br>TGCACGCTGTTAAACCCGCAAGCTCGTGGCTCCGTCCGTTTAGCCTCTAATG<br>ATCCTCTGGCAGACCCCTTGATCGATCCTTGCTTTTTGGATAACGAGCGAGA<br>TATGAGAGTTTTTAAAGAAGGCGCCAGAAAACAACTTGCAGTGTTAAATGC<br>CGCAGCATTATCGCCGTGGCGGGGTGATATGTTATACCCGCTGAATCCGGAT | MFDYIVVGGGSAGAVVA<br>SRLSEDPNVSVCLLESGPV<br>DKSVLIHAPAGVVAMMP<br>RKNKLNYAFETVPQKGLN<br>GRRGYQPRGRCLGGSSSV<br>NAMLYVRGHRADYDHW<br>SALGNAGWSYAEVLPYF<br>KRSEHNENIHDEFHGQGG<br>PLNVMNLRSPGALNAEFI<br>KACEEQGIAATADCNGAE<br>QDGALEYQVTHINGERCS<br>AAKAYLTPNLSRPNLEVM<br>TQTPVLQILLDGKTATGV<br>RVKREGQTLDLRARREVI<br>VCGGAFNSPQLLMLSGIG<br>PGAHLQDRGIAVAHDLPG<br>VGENLQDHIDIVHSYRAP<br>NSDTFGLSFGFMPKILAA<br>MLQWRRQRSGLITSPYAE<br>AGAFFRSSEADGRPDLQLI<br>LVRALVDDHGRKMHLGH<br>GFSSHCTLLNPQARGSVR<br>LASNDPLADPLIDPCFLDN<br>ERDMRVFKEGARKQLAV<br>LNAAALSPWRGDMLYPL<br>NPDDDAALEEDLRNRADT |

| | | |
|---|---|---|
| | GATGATGCTGCTCTTGAAGAAGATCTACGCAACCGGGCTGACACCCAATATC<br>ATCCCGTGGGTACGTGCAAAATGGGGCAGGATGACGGCGCAGTTGTGGACC<br>ATCGCCTTCGCGTTCATGGTATTGCAGGTCTGAGGGTGGCTGATGCAAGTAT<br>AATGCCGACTCTGGTCGCGGGAAACACGAATGCCCCAACTATCATGATTGGT<br>GAGAAAGCAGCTGATATGATCCGTGAAGACGCCGCGTAA | QYHPVGTCKMGQDDGAV<br>VDHRLRVHGIAGLRVAD<br>ASIMPTLVAGNTNAPTIMI<br>GEKAADMIREDAA |
| pZE-PaAAO<br>(WP_21330559<br>9.1) | ATGCAAACCAGCTATGACTACATAGTCATCGGTGGTGGCTCTGCCGGTTCCA<br>CCTTGGCCGGACGCCTGTCTGAGGATCCGCGGATAACGGTCTGTCTGCTGGA<br>GGCCGGGGATTCCGATAAAAGCACGCTGATCCAGATGCCGGCCGCGGTAGT<br>TGCGGTGATGCCGACCCGTCTTCACAACTGGGCTTTTAAAACTGTCCCGCAG<br>CCGGGGCTCAACGGCCGTCGTGGTTATCAGCCCAGAGGTAAAACCCTAGGC<br>GGTTCTTCTTCCATCAATGCGATGCTTTACGTGCGGGGTGATCGCTTAGATTA<br>TGACCATTGGGCCGCCCAGGGAAATCCTGGCTGGTCGTTTGAAGACGTTTTA<br>CCGTATTTCAAGAAAGCAGAACACAACGAAACCCTGGACAATGCCTTTCAC<br>GGAAAAGGCGGCCCTCTGAATGTGGCCGAAATCCAAACGCCTTCTCCCATA<br>CACCAACGGTTTCTGCAGGCGGCGGCTCATTGCGGGTTACCGGGAGTAGCA<br>GATTACAACGGGTTAACCCGCGATGGCGCTTTCATGTATCAGGTTACTCAAA<br>AAAATGGTGAGCGCTGTTCTGCCGCAAAAGCGTATTTGACGCCGAACTTAA<br>GTAGAAAAAACCTTACGGTTCTGACGGGAGCGCAGACGGAGCGTGTTCTGC<br>TGGCGGGTAAGCGTGCGGTTGGAGTACGCGTCAATATAGGCGCCAAAAGCC<br>AGGATCTGCGCGCGTCGCGTGAAGTTATCCTGTCTGCGGGCGCCTTTGGGAG<br>TCCGCAGATTTTAATGCTGAGTGGCATCGGTTCTGGCGCTCAGCTCCAGGCC<br>ATGGGTATAGCCGTGGTGCATGATTTACCGGGCGTTGGAGAAAACCTGCAG<br>GACCACATTGATTATGTATATTCCTACAGCACCCCGAGTAGCCCGGAAACAT<br>TCGGTGTTTCTCTGGGCGGCGGAGCTCGTGTGGTGAAAGGTATTGCGCAGTG<br>GCGTCGCGAGAGAAAAGGTCTCATGACGACGCCGTACGCGGGTGCAGGTGC<br>TTTCGTTCGCTCTAGTCCGGATGTGCCGGCTCCAGACTTACAACTAATCTTCG<br>TCGAAGCCATTGTCGATGATCATGCTCGCAAGCTTCACGTCTCACACGGATT<br>TAGCTGCCATGTAGACGTGCTTAGACCAAAAAGCCGTGGTACAGTGAAGCT<br>GGCCACCACCAACCCTTTTGATGCTCCGCTCATCGACCCCCGCTTCTTCTCTG<br>ACCCAGCCGATATGGATGTTATGCTTAAAGGTGCAAACCTCCAGCGCCGGAT<br>CATGGAAGCTCCGCCGTTAAAAGCACTGGGCGCAAAACCGCTCTATCCGGT<br>GGATCAGTCAGATCAGCGGGCCGTTGAACAAGACATCCGCCGTCGCGCCGA<br>CACCCAGTATCATCCAGTTGGTACCTGCAAAATGGGGTCGGACCCTCTGGCG<br>GTTGTAGATGCGCAATTGCGTGTCCACGGGATAGAGGGTCTGCGCGTGGTTG<br>ATGCCTCTATTATGCCAACCGTTGTAAGCGGTAACACTAACGCGCCGACTAT<br>CATGATTGCCGAAAAAGCAGTCGATATGATTAGACAAGCCGCCAAGTAA | MQTSYDYIVIGGGSAGST<br>LAGRLSEDPRITVCLLEAG<br>DSDKSTLIQMPAAVVAV<br>MPTRLHNWAFKTVPQPG<br>LNGRRGYQPRGKTLGGSS<br>SINAMLYVRGDRLDYDH<br>WAAQGNPGWSFEDVLPY<br>FKKAEHNETLDNAFHGK<br>GGPLNVAEIQTPSPIHQRF<br>LQAAAHCGLPGVADYNG<br>LTRDGAFMYQVTQKNGE<br>RCSAAKAYLTPNLSRKNL<br>TVLTGAQTERVLLAGKRA<br>VGVRVNIGAKSQDLRASR<br>EVILSAGAFGSPQILMLSGI<br>GSGAQLQAMGIAVVHDL<br>PGVGENLQDHIDYVYSYS<br>TPSSPETFGVSLGGGARV<br>VKGIAQWRRERKGLMTT<br>PYAGAGAFVRSSPDVPAP<br>DLQLIFVEAIVDDHARKL<br>HVSHGFSCHVDVLRPKSR<br>GTVKLATTNPFDAPLIDPR<br>FFSDPADMDVMLKGANL<br>QRRIMEAPPLKALGAKPL<br>YPVDQSDQRAVEQDIRRR<br>ADTQYHPVGTCKMGSDP<br>LAVVDAQLRVHGIEGLRV<br>VDASIMPTVVSGNTNAPTI<br>MIAEKAVDMIRQAAK |
| pZE-ScAAO<br>(WP_16956084<br>9.1) | ATGTACGATTATATCGTCGTTGGCGGTGGGTCTGGGGGATCCGTTGTTGCGT<br>CAAGACTGAGTGAAGATCCAAATATTAAGGTCTGTTTGTTGGAAGCAGGTG<br>GGCCTGATAAAAGCATACTGATCCATGCTCCAATGGGTACTGCGGCAATGCT<br>GCCCCGTAAAATTAATAATTGGGCCTTTGAGACTGTACCACAACCGGGCCTT<br>AATAACCGAAAGGGTTATCAACCGCGAGGTAAAACGCTGGGCGGTAGTAGT<br>TCGATCAATGCGATGCTGTATGTGAGAGGTCATCGTTGGGATTACGACCATT<br>GGGCAAGTTTAGGCAATACCGGATGGTCGTACGCCGATGTCCTGCCCTACTT<br>TAAAAAAGCGGAGAACAATGAACGTGGCGCCGACGATTTTCACGGTACAGG<br>CGGCCCACTGAATGTGGCCGACTTACGATCCCCGATGAAGATTGGCGATGTA<br>TTCATGAAGGCGGCGCGCGAATTACAGCTGCCGATGAACAAAGACTTCAAT<br>GGAGCTGAACAAGAGGGCGTGGGCTATTTTCAAACCACCCAAAAAAACGGT<br>GAACGTTGTTCCGCGGCGAAAGGCTATCTGACCCCGAATCTGGATCGCCCCA<br>ATCTGGACATTTTGACGCATGCCACCGCCTCACGTATACTGTTTGAAGGTAA<br>AAAAGCAGTGGGGGTGCAGTACGTGCAAGACGGCCAGACAAAAACACTGA<br>AAGCACGCCAAGAGATTATTCTGAGTGGGGGAGCCTTTGCCACCCCTCAACT<br>GTTACTCCTGAGTGGTGTCGGCGCTCCGGAAAATATAACACCGTTCGGCATT<br>GATATGGTTCACGAGCTGCCAGGAGTAGGAGAAAACTTGCAAGATCATATT<br>GATTATGTATCGGCCTATCGCTCTTCATCAAGAGATTCCATGGGTATTTCGCT<br>GTCTGGCGGCGTTGATCTGATTAAAGCCATCTTTGAATGGCGAAAACAGCGG<br>ACAGGGATGGTGACTACTACGTTTGCGGAGATTGGGGCGTTCTTGAAGACTT<br>CAAACGATATGGAAGTACCGGATTATCAACTTCACTTTGTCATAGGGATGCT<br>GGATGATCACGCCCGTAAAACAAACCTGGGTTACGGGTTCAGCTGTCACTGT<br>TGTGTCCTGCGGCCAAAGTCACGTGGTACAGTAACACTTAACTCCGCAAGAG<br>CAGAGGACGCGCCTCGGATCGATCCCAGATTCTTGGATAATGATGACGATGT<br>TCAAACCCTGCTTACGGGTGTTAAAGAAATGGAACGTATCTTACATGCCCTC<br>GCATTTGAAGGTGTCCGGGGTAAGCGTCTGTACCCTGTTGATTTGAATTCTG<br>ATGCAGCAATGATTGCCGAAATTCGAAATCGAGCTGATACGGTGTATCATCC<br>TGTTGGCACCTGTAAGATGGGAACCGATGACATGGCAGTCGTTGACCCGCA<br>GCTGAAAGTACATGGCCTTGAAGGACTGCGAATTGTTGACGCGTCGATTATG<br>CCTACGCTGGTCGGTGGTAACACAAATGCTCCCACGATAATGATTGGCGAA<br>AAAGCGGCTGATATGATTAAGGCCGACGCCAAAACCCCAGCAATCGCCTAA | MYDYIVVGGGSGGSVVA<br>SRLSEDPNIKVCLLEAGGP<br>DKSILIHAPMGTAAMLPR<br>KINNWAFETVPQPGLNNR<br>KGYQPRGKTLGGSSSINA<br>MLYVRGHRWDYDHWAS<br>LGNTGWSYADVLPYFKK<br>AENNERGADDFHGTGGPL<br>NVADLRSPMKIGDVFMK<br>AARELQLPMNKDFNGAE<br>QEGVGYFQTTQKNGERCS<br>AAKGYLTPNLDRPNLDIL<br>THATASRILFEGKKAVGV<br>QYVQDGQTKTLKARQEII<br>LSGGAFATPQLLLLSGVG<br>APENITPFGIDMVHELPGV<br>GENLQDHIDYVSAYRSSS<br>RDSMGISLSGGVDLIKAIF<br>EWRKQRTGMVTTTFAEIG<br>AFLKTSNDMEVPDYQLHF<br>VIGMLDDHARKTNLGYG<br>FSCHCCVLRPKSRGTVTL<br>NSARAEDAPRIDPRFLDN<br>DDDVQTLLTGVKEMERIL<br>HALAFEGVRGKRLYPVDL<br>NSDAAMIAEIRNRADTVY<br>HPVGTCKMGTDDMAVVD<br>PQLKVHGLEGLRIVDASI<br>MPTLVGGNTNAPTIMIGE<br>KAADMIKADAKTPAIA |
| pZE-RsAAO<br>(WP_24146385<br>5.1) | ATGGCGGAGATGGAATTCGATTACGTCGTGGTTGGAGCCGGTTCAGCGGGA<br>TGCGTTCTGGCCAGTCGGCTGTCTGAGGATCCAAACGTCACTGTATGTCTGC<br>TTGAAGCCGGTGGTCCGGATTCATCAGTATTAATTCATGCGCCGGCGGGCGT | MAEMEFDYVVVGAGSAG<br>CVLASRLSEDPNVTVCLL<br>EAGGPDSSVLIHAPAGVV |

| | | |
|---|---|---|
| | TGTAGCTATGGTGCCGACGAAAATCAACAACTACGGTTATGAAACCGTTCCG<br>CAGGCAGGGCTGTATGGTCGTCGTGGTTACCAGCCTCGCGGTAAGACGCTG<br>GGGGGAAGTTCTTCAATTAATGCAATGCTGTACGTTCGTGGTAATCGGTGGG<br>ATTATGATCATTGGGCGGGTCTTGGTAATCCAGGCTGGTCATACGATGATGT<br>CTTGCCGCTATTTAAAAGAAGCGAAAATAACGAGCAGTTCCAGGATAGCTTT<br>CATGGCCAAGGAGGGCCGCTGAATGTCACGTATCCCCGTTTTGATAGTCCAT<br>TGTCCCAGATGTTTTTGGATGCAGCTGCCCAGAACGGAATGGCGCTGAATCC<br>GGATTATAACGGCGCGCAGCAGGATGGAGCATTTCTGTATCAGGTCACACA<br>TAAGAATGGCGAGCGTTGTTCAGCAGCAAAAGGGTATCTCACGCCTCACCTC<br>AACCGACCGAATCTGAAGGTGATCACGAGAGCAGTTACCGCTAAAATCAAT<br>GTGCAAGATCGCCGTGCAACAGGCGTGCAGTTTTATCAGAGCAACCAACTG<br>CAGACCGTCCGGGCAAGGCGGGAGGTTGTCGTTTCGGCCGGGGCGTTTGGC<br>TCCCCCCAACTGTTGCAGCTTTCGGGGATAGGCGCAGCCCAAGATTTGCAAA<br>AGCTTGGGATTGCCGTGACACATAACCTTCCGGGTGTCGGGCAGAACCTACA<br>AGACCATATTGATTATGTTCAGAGCTGGTCAGTCCCTTCTAATACGGCGAGT<br>TTTGGTGTTTCACTGCGTGGTACCGGAAAGGCATTGGGAGCAATCATGGAGT<br>GGCGCAAACAGAGAAGCGGTATGGTGACTAGCGCTATTGCCTCAGCTGGTG<br>CATTCTTCAAGTCTCATCCAGATGTGGCAGTCCCTGACTTACAGTTAGTGTTC<br>GTGTTAGCATCGGTGGATGATCATGCGCGCAAACCACATCTGGGTCATGGTA<br>TTAGCTGCCATGTGGATGTCCTGCGGCCGTATAGCAGAGGAACGGTCGGCCT<br>ACAGAGCACAGATCCCAGAGTGGCGCCCGTTATAGACCCGCGTTTTTGGCT<br>GATGAACGTGACATGCAGCTGCTGTTAAAGGGCGGGCAAGTTCAACAGCGG<br>TTGATTGAAAGCTCCGCCTTCGCCCCTGTGCGTGGTAAAATGATGTACCCAG<br>TTCGTGTTGACGATATTCAAGCGATGGAACAGGATATTCGTAATCGTGCAGA<br>TACACAGTACCATCCCGTCGGTACGTGTAAGATGGGGCCGGATAGTGATGC<br>AATGGCTGTAGTCGATGCCCAATTGAGGGTGAAAGGTATTGCGGGACTGCG<br>GGTAGCCGATGCGTCAATTATGCCAACGCTTATTGGGGGAAATACGAATGCT<br>CCTTCCATTATGATAGGCGAGAAAGCAGCAGACTTGATTCGTGCCGCCCACT<br>GA | AMVPTKINNYGYETVPQA<br>GLYGRRGYQPRGKTLGGS<br>SSINAMLYVRGNRWDYD<br>HWAGLGNPGWSYDDVLP<br>LFKRSENNEQFQDSFHGQ<br>GGPLNVTYPRFDSPLSQM<br>FLDAAAQNGMALNPDYN<br>GAQQDGAFLYQVTHKNG<br>ERCSAAKGYLTPHLNRPN<br>LKVITRAVTAKINVQDRR<br>ATGVQFYQSNQLQTVRA<br>RREVVVSAGAFGSPQLLQ<br>LSGIGAAQDLQKLGIAVT<br>HNLPGVGQNLQDHIDYV<br>QSWSVPSNTASFGVSLRG<br>TGKALGAIMEWRKQRSG<br>MVTSAIASAGAFFKSHPD<br>VAVPDLQLVFVLASVDDH<br>ARKPHLGHGISCHVDVLR<br>PYSRGTVGLQSTDPRVAP<br>VIDPRFLADERDMQLLLK<br>GGQVQQRLIESSAFAPVR<br>GKMMYPVRVDDIQAMEQ<br>DIRNRADTQYHPVGTCK<br>MGPDSDAMAVVDAQLRV<br>KGIAGLRVADASIMPTLIG<br>GNTNAPSIMIGEKAADLIR<br>AAH |
| pZE-BaAAO<br>(WP_25876749<br>9.1) | ATGAGCCACACAGAGTTCGATTATATCGTAGTGGGAGCTGGTAGTGCGGGTT<br>GCGTATTAGCCTCCCGCCTTACTGAAGACCCTAATGTGACTGTCTGTCTTTTA<br>GAGGCAGGCGGTCATGATACGGGCGTTCTAATCCGCGCACCCGCGGGCTAT<br>GTGGCCATGGTCCCTACTAGAATTAACAATTATGCATATCAGACGGTCCCAC<br>AAGCGGGCCTGAGTGACCGTCGCGGTTATCAGCCACGCGGTAAAACCTTGG<br>GGGGTTCTTCCTCAATCAATGCGATGTTTTATGTGCGTGGAAACCGCTGGGA<br>CTATGATCATTGGGCCAGCCTCGGAAATCCGGGCTGGAGTTATGATGATGTG<br>CTTCCTCTTTTCAAACGGGCAGAAAATAATGAACAGTTCAGGGATGATTTTC<br>ATGGACAGGGAGGACCACTGAACGTTACGTATCCTAGACACCAATCGCCGC<br>TAACCAAACTCTTTCTTGAGGCAGCCGCTCTGCATCAAATTAGACTGAATCC<br>GGACTACAATGGCGCCGATCAGGAGGGCGCTTTTGAATACCAAGTAACACA<br>AAAAGGGGGAGAACGTTGCTCTGCAGCCAACGCGTACTTAACACCTAACCT<br>GTCGCGCAATAACTTAGAAGTTCTGACGTCCGCTATCAGCGCCACTGTGTTG<br>CTCGAAGATCGACGGGCGTATGGAGTCTCGTACTATCAGGAAGGCGAACTG<br>AAGGAAGTGAGGGCCCGAAGAGAGGTCATTCTGGCAAGCGGCGTTTTTGGC<br>AGCCCCCAACTGCTGCAGTTGTCAGGCATTGGTCCTGGAGAAGAACTTAAG<br>AAATTCGATATTCCGGTAGTGAACGAACTGCCGGGCGTAGGTAAAAATCTC<br>CAAGATCATATTGACTACATTCAATCTTGGTACGTACCCTCTGACACAGAAA<br>CGGTGGGCATCTCTTTACGGGGGGGACTGAAACTGGCGAGGGCTGCTTTTGA<br>ATGGAGCAAAAACCGTACGGGTCTTATAACAACCACATACGGAACAGCCGG<br>TGCCTTTTTACGCTCCTCGCCGGAAGTGCAAATTCCTGATCTGCAGCTTATAT<br>TTGTAATCGCCCTGGTCGACGATCACGCTCGTAAACTGCATTTGGGCCACGG<br>CATCTCCTGTCATGTCGACGTTCTCCGTCCGTATTCCCGAGGTACGGTCGGCT<br>TAGACAGTTCCGATCCATGGGACGCCCCGCGTATTGATCCGAATTATTTCAG<br>TGATGAGCGTGATTTGAAACTCCTGGTACGCGGAGCTCTGGTTCAACAGCGA<br>ATTATAGAATCCCCGCCGTTTGGCCGACTGCGTGGGAAGATGTTATACCCGA<br>CAAGGCTTGATGATCTTGCGGGCATCGAAAAAGACATTCGCTGCAGGGCAG<br>ATACCCAGTACCATCCGGTGGGCACATGTAAAATGGGCCCCAATGGGGATC<br>CGATGGCAGTGGTTGACGCAAAATTGCGCGTTCGGGGGGTCGATGCTTTGCG<br>CGTTGTAGATGCGTCGATAATGCCTACGATTGTTGGCGGGAATACCAATGCT<br>CCAACGATTATGATTGCTGAGAAAGCAACGGATCTTATCCGCAATAGTGTTT<br>AA | MSHTEFDYIVVGAGSAGC<br>VLASRLTEDPNVTVCLLE<br>AGGHDTGVLIRAPAGYVA<br>MVPTRINNYAYQTVPQAG<br>LSDRRGYQPRGKTLGGSS<br>SINAMFYVRGNRWDYDH<br>WASLGNPGWSYDDVLPL<br>FKRAENNEQFRDDFHGQG<br>GPLNVTYPRHQSPLTKLFL<br>EAAALHQIRLNPDYNGAD<br>QEGAFEYQVTQKGGERCS<br>AANAYLTPNLSRNNLEVL<br>TSAISATVLLEDRRAYGVS<br>YYQEGELKEVRARREVIL<br>ASGVFGSPQLLQLSGIGPG<br>EELKKFDIPVVNELPGVG<br>KNLQDHIDYIQSWYVPSD<br>TETVGISLRGGLKLARAA<br>FEWSKNRTGLITTTYGTA<br>GAFLRSSPEVQIPDLQLIFV<br>IALVDDHARKLHLGHGIS<br>CHVDVLRPYSRGTVGLDS<br>SDPWDAPRIDPNYFSDER<br>DLKLLVRGALVQQRIIESP<br>PFGRLRGKMLYPTRLDDL<br>AGIEKDIRCRADTQYHPV<br>GTCKMGPNGDPMAVVDA<br>KLRVRGVDALRVVDASI<br>MPTIVGGNTNAPTIMIAEK<br>ATDLIRNSV |
| pZE-MlAAO<br>(WP_02377925<br>2.1) | ATGACTTTCGATTATGTAATTGCCGGTGGGGGCAGTGCCGGGTGTACTCTTG<br>CCGCCAGATTAAGTGAAGACCCGTCTAAGACCGTTTGCTTAGTAGAGGCCG<br>GAGGTGAAGGAAAAAACCTGTTTATTCGAGCCCCTGCAGGGGTGATCGCGT<br>TACTTCCGGGGCGGCCGAAGATCCATAACTGGGCATTCGAGACAGTACCAC<br>AGCAGGGACTGGGTGGTCGTAAAGGGTATCAGCCCCGCGGGAAGGCCTTGG<br>GTGGAAGTAGTGCGATCAACGCTATGCTGTATGTTAGAGGACATCGCAGTG<br>ATTACGACGAGTGGGCTAATTTTGGTTGTGACGGATGGTCGTGGGATGAAGT<br>TTTGCCTTATTTCCGGAGGGCTGAAGGCAATCAACGGGGATCCGATGCATTA<br>CATGGGGATGACGGTCCCCTTAGGGTGGCAGAACAGCAAGAGCCGAGAGCG | MTFDYVIAGGGSAGCTLA<br>ARLSEDPSKTVCLVEAGG<br>EGKNLFIRAPAGVIALLPG<br>RPKIHNWAFETVPQQGLG<br>GRKGYQPRGKALGGSSAI<br>NAMLYVRGHRSDYDEWA<br>NFGCDGWSWDEVLPYFR<br>RAEGNQRGSDALHGDDG<br>PLRVAEQQEPRALSRAFV |

| | | |
|---|---|---|
| | TTAAGTAGAGCGTTCGTTGAAGCATGTGGTGAAAATCAGATTCGCCGTAATG<br>ATGATTTTAACGGCCCAGAGCAGGAAGGTGCGGGACTGTACCAGGTAACAC<br>AGTTTTGGGGGGGCGACCGTAATGGGGAACGCTGCTCTGCCGCAGCTGCTTA<br>TCTGCACCCCGCAATTGGCCGTCCCAACCTGACCGTGATTACAGGGGCACAT<br>GCGACCGGTATTGTGCTTGATGGGAAAAGGGCAACAGGTGTAAGGTACCGC<br>GAGGGTAATAGCGAAGCGGTTGCTCGCGCCCGGCGAGAGGTAATCGTGTGC<br>GGCGGTGCATTTGGAAGTCCCCAGTTGCTCCTGCTGAGCGGTGTAGGTCCTG<br>CGGTGGAATTGGCCGCTCACGGTATCCGAATGGTGCATGAACTTCCGGGCGT<br>TGGCAAGAACCTGCAAGATCATCTAGATTTTATAATGGGATGGACTAGCCGT<br>GATGCAGATATGATGGGAATTGGTCTGCGGGGCTTGCCAGGTTTGTTACGCC<br>ATATGCTACGTTGGCGTAAGGACGGGGGTGGTATGATCGCGACACCATATG<br>CGGAGGGAGGGGCGTTCCTGAAAAGCGATCCGGAGATAGATCGCCCGGATT<br>TGCAACTGCACTTTTGCATTGCCATAGTCGACGATCATGGTAGGAAACTGCA<br>CATGGGTTATGGTTTTTCCTGTCACGTATGCGTCCTCCGGCCGCACAGTCGC<br>GGTGAGGTTGGCCTGTCGACCCCAGATCCCCTGGCTCCGCCGCGTATTGACC<br>CGCGCTTTCTTAGTGATGAAAGGGATGCCCATCTATTATTGAAAGGCGTGCG<br>CACGATGCGAGGCATCTTAGAAGCCCCCGCGTTGGCGAAGTACCGGGCCAA<br>AGAGATTTACACGGCAGGTGCATCGAGTGACGCGGAGCTGATGGCTCACAT<br>TCGAGCCCGTGCAGACACCGTCTATCATCCTGTGGGTACTTGCCGTATGGGT<br>GTGGACGACATGGCGGTCGTGGACCCACAGCTGAGGGTGCGAGGCATGCAG<br>GCGTTACGAGTTGTGGATGCATCCGTGATGCCGACGCTTATTGGTGGCAACA<br>CCAACGCACCGACCATCATGATCGCCGAAAAGGCAGCGGACATGATTAAAA<br>CGGCTGCTTAA | EACGENQIRRNDDFNGPE<br>QEGAGLYQVTQFWGGDR<br>NGERCSAAAAYLHPAIGR<br>PNLTVITGAHATGIVLDG<br>KRATGVRYREGNSEAVA<br>RARREVIVCGGAFGSPQL<br>LLLSGVGPAVELAAHGIR<br>MVHELPGVGKNLQDHLD<br>FIMGWTSRDADMMGIGL<br>RGLPGLLRHMLRWRKDG<br>GGMIATPYAEGGAFLKSD<br>PEIDRPDLQLHFCIAIVDD<br>HGRKLHMGYGFSCHVCV<br>LRPHSRGEVGLSTPDPLAP<br>PRIDPRFLSDERDAHLLLK<br>GVRTMRGILEAPALAKYR<br>AKEIYTAGASSDAELMAH<br>IRARADTVYHPVGTCRMG<br>VDDMAVVDPQLRVRGM<br>QALRVVDASVMPTLIGGN<br>TNAPTIMIAEKAADMIKT<br>AA |
| pZE-CcAAO<br>(WP_10123142<br>8.1) | ATGAAGAATACCCTGGATAAGTTTGACTATATCATTGTTGGCGCCGGATCAG<br>CGGGATGTGTCGTGGCGGCGAGACTGTCAGAAGATCCCAATGTATCAGTCT<br>GTCTCCTGGAAGCGGGTGGCCCCGACAAGAGCGTATTTATCCATGCGCCGAT<br>AGGGCTGGCCGCTATGTTGCCCACGAAGTTAAACAATTGGGCTTTTGAAACA<br>ATCCCGCAAGCCGGGTTAAATGGGCGCAAAGGTTACCAGCCTCGTGGCAAG<br>ACACTAGGGGGTTCGTCCAGCACAAACGCCATGCTGTACGTGAGAGGTAAT<br>AAGTGGGATTACGATAATTGGGCCGCGCTGGGCAACGAAGGCTGGTCCTAT<br>GAAGATATCCTTCCTTATTTCAAAAAGTCAGAGGCGAATGAAGTTTTCAACG<br>ACAAGTACCATAATGTAGATGGTCCTTTAGGTGTCTCTAGTGCGTCCCATGC<br>TTCGGATCTAAACCAAATGTTTATTGATTCGTGCGTCCAGCAGGGTATTAAA<br>CATACTGATGATTGTAACGGTGCAGAACAAGAAGGTGTGTTTTTATACCAGA<br>GAACCATCAAAAATGGCGAACGATGTTCCGCGGCCAAAGCTTATCTAACAC<br>CGAATAAGGCACGAAAGAATCTGACAATAATAACACATGCATTAACGGAAA<br>AAGTGTTGTTTGAAGACAAGACCGCTGTGGGTGTTCGTTACAAGAAAAACA<br>ATAAAACTATTGAGATATTATGTAATAAAGAGGTGATTTTGTCGTCTGGAGC<br>CTTTGGAAGTCCGCAGATTCTGATGTTATCAGGTGTGGGTGCCAAACAGCAT<br>CTGAAGGATAAAGGTATTACTTCTGTCCATGATCTGCCGGGGGTGGGGCAA<br>AATCTGCAAGACCATATTGACTACGTTCAAACGTTTAAGGTTGATAGCCGTC<br>ATGATACGTTTGGTTTTTCCGTGCGCGGAAGCTTCCGTGTGCTAAAATGGAT<br>CTCAGAATGGCGGAAAACTCGGACAGGTAAAGTGACGAGCTCTCTCGCTGA<br>GTCCGGGGCTTTCTTTTCGACAGAATCGGATTTGGTCGCGCCAGACGCACAA<br>CTGATCTTTGTTCCAGCAATTGCAGATAACCATGCACGTACAGTGAACTTTG<br>GGCACGGATACTCTTGTCACATAACCCTTCTGCGTCCCGATTCCGTAGGTGA<br>GGTTAAATTGAATTCATCGAATCCCGAAGATAGTCTGGCGATAGACCCGAAT<br>TTTTTCACCAAGATAAAGATATGGAGATTATTAAACGGGCCGCCAAAAAA<br>ATGCAGGACATTCTTGAAGGAAAGCCCTTTTCAAGCATACGGAAAAAAATG<br>CTGTATTTCGTTGAGAACGGTAATGATGAACAGCTTGAAAAAGATATAAGG<br>AATCGTGCGGATACACAGTATCATCCATGCGGCACTTGTAAAATGGGGCCTG<br>TAGCTGACGAGATGGCTGTGGTAGATAGTCAGCTGCGGGTCCACGGCATGC<br>AGAATCTTCGTGTTGTTGATGCCTCTATCATGCCCAAAATTATTACTGGCAA<br>CACAAATGCACCAACCATAATGATTGGTGAAAAGGCTGCAGATATGATTCT<br>CGCGAACTAA | MKNTLDKFDYIIVGAGSA<br>GCVVAARLSEDPNVSVCL<br>LEAGGPDKSVFIHAPIGLA<br>AMLPTKLNNWAFETIPQA<br>GLNGRKGYQPRGKTLGG<br>SSSTNAMLYVRGNKWDY<br>DNWAALGNEGWSYEDIL<br>PYFKKSEANEVFNDKYHN<br>VDGPLGVSSASHASDLNQ<br>MFIDSCVQQGIKHTDDCN<br>GAEQEGVFLYQRTIKNGE<br>RCSAAKAYLTPNKARKNL<br>TIITHALTEKVLFEDKTAV<br>GVRYKKNNKTIEILCNKE<br>VILSSGAFGSPQILMLSGV<br>GAKQHLKDKGITSVHDLP<br>GVGQNLQDHIDYVQTFK<br>VDSRHDTFGFSVRGSFRV<br>LKWISEWRKTRTGKVTSS<br>LAESGAFFSTESDLVAPDA<br>QLIFVPAIADNHARTVNFG<br>HGYSCHITLLRPDSVGEV<br>KLNSSNPEDSLAIDPNFFH<br>QDKDMEIIKRAAKKMQDI<br>LEGKPFSSIRKKMLYFVEN<br>GNDEQLEKDIRNRADTQY<br>HPCGTCKMGPVADEMAV<br>VDSQLRVHGMQNLRVVD<br>ASIMPKIITGNTNAPTIMIG<br>EKAADMILAN |
| pZE-LeAAO<br>(WP_25141942<br>7.1) | ATGCAGGCTGACTATGTGATTATCGGGGGAGGCTCTGCGGGGGCCTCATTAG<br>CGGCCCGTCTGAGCGAAGATCCGGCGACAACAGTCTGTCTCCTGGAGGCAG<br>GCGGGGGAGGCAAGTCTATATTTGTCCGTGCGCCGGCCGGCACTGTTGCAAT<br>GTTACCGGGCTGGGGTAAAATTAATAATTGGGCTTTTAAAACCGCCCCTCAG<br>GGAGGACTTAATGGTCGAAGAGGTTACCAGCCCCGCGGCCGCGCTTTAGGT<br>GGATCTTCGGCTATCAACGCTATGCTGTATGTTCGCGGACATCGCGCCGATT<br>ACGACCAGTGGGCTGCGCTGGGTTGCGAAGGCTGGGATTGGGACAGCGTCC<br>TACCGTATTTTAAGCGCTCTGAGGGGAATGAAAGGGGGGCGGACGCGGCTC<br>ATGGCGCAGACGGCCCCCTGCAAGTGAGCGATCAAAACGACCCGCGCCCAA<br>TTAGTCGCGCGTTCATAGAGGCCGCAAAGCAGATGCAGATAAGAGAACGTC<br>AAGATTTCAATACCGGCGATAATGAAGGTGTGGGGTTATATCAGGTAACTC<br>AGTTCCATGATGAAAAACGTCGCGGAGAACGCTGCAGTGCCGCTGCCGCAT<br>ATCTCCACCCTGTGATGGAGAAAAGGAAAAATTTAATCGTTCTGACTCACGC<br>TCGCGCAACCCGCATCCTGTTTGACGGGAAACGGGCAGTTGGTGTGGCATTC<br>CGGCAGGGTCGTCGGGCACAGACCGTCACAGCACGGCGTGAGGTCATCCTG | MQADYVIIGGGSAGASLA<br>ARLSEDPATTVCLLEAGG<br>GGKSIFVRAPAGTVAMLP<br>GWGKINNWAFKTAPQGG<br>LNGRRGYQPRGRALGGSS<br>AINAMLYVRGHRADYDQ<br>WAALGCEGWDWDSVLP<br>YFKRSEGNERGADAAHG<br>ADGPLQVSDQNDPRPISR<br>AFIEAAKQMQIRERQDFN<br>TGDNEGVGLYQVTQFHD<br>EKRRGERCSAAAAYLHPV<br>MEKRKNLIVLTHARATRI<br>LFDGKRAVGVAFRQGRR<br>AQTVTARREVILCGGAFN |

| | | |
|---|---|---|
| | TGTGGTGGAGCATTCAACTCTCCCCAGATCCTGCAGTTAAGCGGGATCGGGG<br>CTGCCGATGATCTGAGACCGCATGGCATTGAACCACTCCATGAATTACCCGG<br>AGTTGGTAAAAATTTGCAAGATCACTTAGATTTTACCTTAGCATGGAAAAGT<br>AAGGATACCGATAATTTCGGCATCGGTCTTACCGGCGGTGCGAATCTGCTGC<br>GGCATATGTTGAGGTGGAGAAAGGACGGGGGCGGTATGATCGCTAGCCCGT<br>TCGCCGAAGGAGCGGCGTTTTTAAAGACCGCTCCAACTCTGGATCTCCCAGA<br>TATACAGCTTCATTTTGTGATCTCGATTGTAGATGATCATGCAAGACGCTTA<br>CATCCGGGCTATGGCTTTAGCGTTCATGTTTGCGTGCTACGCCCGAGGTCTC<br>GCGGTGTTGTAGGTCTGGAAAGTGCAGATCCGCTCGCGGCTCCGCGGATTGA<br>CCCACGGTATCTTTCAGATCGAAGGGATCTCGATACGTTAATTGCCGGAGCC<br>AAGCTGACAAGACAGATAGTTATGCAGGAGCCCATGGCTCGCTATAGACAT<br>AAAGAGATGTTTGGACTGCATGACGGTCTGTCCGATGCCGAATGGGCGGCG<br>CACATACGAGCCAGGGCTGATACGATATATCATCCCGTTGGTAGTTGTCGTA<br>TGGGAGTGGATGATATGGCTGTAGTGGGTCCAGATCTCCGCGTTCACGGGTT<br>AGAAGGTGTTCGGGTAGTGGACGCCTCAGTTATGCCTACCCTCATCGGCGGT<br>AATACAAACGCTCCGACGATCATGATCGCAGAAAAGGCCGCAGATATGATT<br>TGTGACGTTGCTTAA | SPQILQLSGIGAADDLRPH<br>GIEPLHELPGVGKNLQDH<br>LDFTLAWKSKDTDNFGIG<br>LTGGANLLRHMLRWRKD<br>GGGMIASPFAEGAAFLKT<br>APTLDLPDIQLHFVISIVDD<br>HARRLHPGYGFSVHVCVL<br>RPRSRGVVGLESADPLAA<br>PRIDPRYLSDRRDLDTLIA<br>GAKLTRQIVMQEPMARY<br>RHKEMFGLHDGLSDAEW<br>AAHIRARADTIYHPVGSC<br>RMGVDDMAVVGPDLRV<br>HGLEGVRVVDASVMPTLI<br>GGNTNAPTIMIAEKAADM<br>ICDVA |
| pZE-RbAAO (MDJ0612750.1) | ATGGAATTTGACTATGTTATTGTAGGCGGGGGTTCCGCAGGTTCCACCCTTG<br>CTAGTCGTTTAACCGAAGATCCGGCCATTCGCGTATGTCTTCTTGAAGCGGG<br>CGGTGATGGGAAGGACCTCTTAATTCGTACCCCTTTGGCGGTGGTAGCGATG<br>TTACCAGGACGCCCGAAAATCAATAATTGGGCTTATGAAACTGTCCCACAA<br>GATGGGCTTAATGGTCGTAAAGGATACCAGCCTCGGGGCAAAGCTCTTGGA<br>GGAAGTTCAGCCATTAATGCGATGTTATATGTCCGTGGACACCCAACAGACT<br>ATGATGATTGGGCCAATTCTGGATGCGAGGGTTGGTCCTGGGAAGAGGTAC<br>TTCCGTATTTTCAGAAATCTGAAAACAATCAGCGCGGTGCGGATACTTTACA<br>TGGCTCCGGCGGGCCTCTTGAAGTATCTGATCAAAAAGCTCCCCGGGCGATC<br>ACAGTTGCTTTTATTGATGCTGCAGCCGAACTTCAGCATAGACGTAATGACG<br>ATTTTAATGGTCCGGAACAGGAGGGTGTAGGAAAATATCAGGTAACTCAGT<br>TTCATCGCAAAGACAAAAATGGCGAACGCTGTTCGACAGCGGCAGCGTATC<br>TGCACCCGGTAATGAGTAGACCTAACCTGACCGTTGTGACAAAGGCCACGG<br>CCTCACGCATCCTGATGGAAGGCAAACGCGCTATCGGAGTCGAATATTTTCA<br>AGGTAAAACAGAAAAACAGGTGATGGCCAAGCGTGAAGTCCTTCTGTGTGG<br>CGGTAGCTTCAATTCACCGCAGCTCTTGCAGCTGTCAGGTATTGGGCTTCCT<br>GAGGACATCCAGCCCCATGGGATCGAAATGATTCATGAGCTCGGTGGTGTC<br>GGCCAGAACCTGCAGGACCATCTTGATTTTGTTCTTGCATTCAAGTCCAAAG<br>ACAAGGATAACGTGGGCCTTTCGTTGGGAGGCGCTGCCGGGCTCGTTTCACA<br>TATCCTAAAGTGGCGCAAAGATGGTAATTCAATGGCGGCCACGCCATTCGC<br>AGAAGGTGGTGCCTTTCTGAAGACCGATAATTCACTTAGCCGCCCGGATATT<br>CAGCTGCATTTTGTGATTGGCATTGTTGATGATCATGCCCGTAAGCTCCATCT<br>GGGCCACGGTTTCTCCTGTCACGTTTGTCAGCTGAGGCCTCACTCACGGGGT<br>TCAGTGAGTCTTCTCGACTCAAACCCGATGAGCGCCCCGTTGATAGACCCTC<br>AATATTTAAGCGATGAACGCGATCTGCCGGTGCTGTTAAAAGGCGCAAAAA<br>TGAGCAGAGATATTCTCATGGCGCCAGCCTTGGAAAATTATCGTCACAAAG<br>AGCTGTTCGGTATTCGTGACGGTTTGTCGGACGGCGAATGGGAGGCGCATAT<br>TCGTAGCCGCGCTGATACAATTTACCACCCCGTCGGTACGTGTAAAATGGGG<br>ATGGACGATAATGCGGTAGTAGATCCGGAGTTGAAAGTTAGAGGCCTCGAA<br>GGCCTGAGAGTGGTGGACGCGAGTGTCATGCCGACCTTAATAGGAGGCAAT<br>ACCAATGCACCGACGATCATGATAGCGGAGAAAGCCGCGGATTTGATTAAA<br>GCTGAGCTGCTGTAA | MEFDYVIVGGGSAGSTLA<br>SRLTEDPAIRVCLLEAGGD<br>GKDLLIRTPLAVVAMLPG<br>RPKINNWAYETVPQDGLN<br>GRKGYQPRGKALGGSSAI<br>NAMLYVRGHPTDYDDW<br>ANSGCEGWSWEEVLPYF<br>QKSENNQRGADTLHGSG<br>GPLEVSDQKAPRAITVAFI<br>DAAAELQHRRNDDFNGP<br>EQEGVGKYQVTQFHRKD<br>KNGERCSTAAAYLHPVM<br>SRPNLTVVTKATASRILM<br>EGKRAIGVEYFQGKTEKQ<br>VMAKREVLLCGGSFNSPQ<br>LLQLSGIGLPEDIQPHGIE<br>MIHELGGVGQNLQDHLDF<br>VLAFKSKDKDNVGLSLGG<br>AAGLVSHILKWRKDGNS<br>MAATPFAEGGAFLKTDNS<br>LSRPDIQLHFVIGIVDDHA<br>RKLHLGHGFSCHVCQLRP<br>HSRGSVSLLDSNPMSAPLI<br>DPQYLSDERDLPVLLKGA<br>KMSRDILMAPALENYRHK<br>ELFGIRDGLSDGEWEAHIR<br>SRADTIYHPVGTCKMGM<br>DDNAVVDPELKVRGLEG<br>LRVVDASVMPTLIGGNTN<br>APTIMIAEKAADLIKAELL |
| pZE-VjAAO (WP_286617962.1) | ATGAGTATCGAGGAATTTGATTACGTGGTGGTTGGGGGTGGAAGTGCCGGTT<br>GCGTTGTTGCTAGTCGACTGAGCGAGGATCCGCGTGTGACAGTGTGCCTGCT<br>GGAGGCCGGGTCAGCTGATTCGAGCGTATTTATCCATGCACCTGCTGGGGTT<br>GTGGCAATGTTACCCATTCCGTATAAAAATTGGGCCTTTAAAACCGTTCCGC<br>AAAAAGGTCTGAATGGCCGCCGAGGGTATCAGCCACGCGGTAAAGTACTAG<br>GAGGCTCATCCTCAACTAATGCTATGCTATATGTGCGCGGCAATCGGTGGGA<br>CTATGATCACTGGGCGTCACTGGGAAACAGAGGTTGGTCGTATGAAGATGTT<br>CTTCCGTATTTTAAAAGAGCTGAGGCAAATGAGACGCACGGTAATTGTTCTT<br>ACCATGGTAGTTCAGGTCCTCTCAACGTAGCTGAATTACGCTCCCCAAGCGC<br>ACTGAATAAAGCATTTCTGGGCGCTGCGGCCATGAAAGGTGTGCCCAATGTT<br>GCGGATTACAACGGCGCCGAACAGTTTGGAAGCTTTATGTACCAAGTCACCC<br>AAAAAAATGGTGAACGCTGTTCAGCAGCCAAGGGCTACATTACACCGCATC<br>TGGCACGTCCCAACCTGTGCGTGAAAACTAATGCGCTGTCGAGCCGTATTGA<br>TTTTGAAGGACGTCGCGCCTGCGGTGTGACCTATACGGCCGCCGGTGCGGAA<br>AAACAAGTCAGAGCGCGCCGAGAGGTGATCCTTTCCAGTGGTGCCTTTGGCT<br>CGCCACAGCTTTTAATGCTGAGCGGGATCGGGCCGGGGGCCGCCTTACAGC<br>GGATGGGCATTCCGGTTATCGGAGATCTTCAGGGCGTAGGCGAAAATCTTCA<br>AGATCATATTGATCATGTTCAATCATACCGCGCTCGGTCAGACAGCCCAACC<br>GTCGGCTTAAGCCTACGGGGTGGCCTGAAAGTGGCGCGGGCCATACCAGAA<br>TGGAAAAAACACCGAACGGGTCTGGTCACCACGAATTATGCGGAGTCAGGG<br>GCGTTCGTGCGATCCTCTCCATCTGTCAGTGTGCCAGATCTCCAGCTGGTTTT | MSIEEFDYVVVGGGSAGC<br>VVASRLSEDPRVTVCLLE<br>AGSADSSVFIHAPAGVVA<br>MLPIPYKNWAFKTVPQKG<br>LNGRRGYQPRGKVLGGSS<br>STNAMLYVRGNRWDYDH<br>WASLGNRGWSYEDVLPY<br>FKRAEANETHGNCSYHGS<br>SGPLNVAELRSPSALNKA<br>FLGAAAMKGVPNVADYN<br>GAEQFGSFMYQVTQKNG<br>ERCSAAKGYITPHLARPN<br>LCVKTNALSSRIDFEGRRA<br>CGVTYTAAGAEKQVRAR<br>REVILSSGAFGSPQLLMLS<br>GIGPGAALQRMGIPVIGDL<br>QGVGENLQDHIDHVQSYR<br>ARSDSPTVGLSLRGGLKV<br>ARAIPEWKKHRTGLVTTN<br>YAESGAFVRSSPSVSVPDL<br>QLVFVVALVDDHSRKVH |

| | | |
|---|---|---|
| | CGTTGTGGCACTGGTGGATGACCACTCACGTAAAGTACATCTGGGCCACGGT<br>TATTCCTGTCATATAGAAGTATTGCGCCCCCACTCCAGAGGCAATGTTCGCC<br>TCGCTTCACGGGATCCTAGAGCCGCACCCCTTATTGATCCGAAGTTTCTCGA<br>TGATCAACGGGATTTGGACTTGTTGGTGAAGGGTGTCCAATTACAAATGGAT<br>ATCCTGGAGGCTTCGCCCTTCGATCCGTATAGAGGTAAAATGTTATATCCGG<br>TGAATCGTAATGATTCTGCGGCTATCGCCGAAGACATCCGGAATAGGGCCG<br>ACACTCAATACCACCCTGTCGGAACATGTAAAATGGGGGTGGCAAGCGATC<br>CAGCTGCGGTGGTAGATGAAAGGCTGCGAGTTCATGGGGTAGAAGGTCTGC<br>GCGTTGTGGACGCTAGTATCATGCCGACACTGTGTGGCGGGAACACTAATGC<br>GCCGACTATAATGATTGGTGAAAAGGCAGCTGATATGATTCGCGCAGATAT<br>GCGCGCCTGA | LGHGYSCHIEVLRPHSRG<br>NVRLASRDPRAAPLIDPKF<br>LDDQRDLDLLVKGVQLQ<br>MDILEASPFDPYRGKMLY<br>PVNRNDSAAIAEDIRNRA<br>DTQYHPVGTCKMGVASD<br>PAAVVDERLRVHGVEGL<br>RVVDASIMPTLCGGNTNA<br>PTIMIGEKAADMIRADMR<br>A |
| pZE-RhsAAO<br>(TAM24039.1) | ATGGAATTCGATTACGTCATCGTCGGGGCCGGCTCTGCCGGCTGCGTGCTCG<br>CCAGTCGCCTGTCTGAAGATCCTGGTATAACAGTTTGCCTCCTGGAAGCCGG<br>AGGTCCGGACAAGTCAGTACTGATTCACGCACCCGCGGGCGCTGCTGCAAT<br>AATTCCTAGTAAAATGAACAACTGGGCTTTTGAAACCGTTCCACAAAAAGG<br>GCTTAATGGTCGTATGGGTTATCAGCCACGCGGTAAAACGTTGGGCGGTTCG<br>TCTTCAATAAATGCTATGGTATATGTACGGGGTAATCGTTGGGATTATGATC<br>ACTGGGCAGCCCTGGGCAATCCGGGCTGGTCTTACGATGATGTGCTCCCGTT<br>TTTCAAGAGGGCCGAAAATAATGAGCAGTTGCGTAACGAATGGCACGGACA<br>GGGTGGTCCGTTAAATGTAACGTACCCCCGCCATAATTCCCCCCTAAATCGG<br>ATGTTCTTGGATGCCGCAGCCATGAACGGCCTGCCCCTGAACCCTGATTATA<br>ACGGTGCCGGTCAACACGGCGCGTTTATGTACCAGGTCACACAAATTAACG<br>GTGAACGTTGCTCAGCTGCCAAAGCATACCTCACGCCGCAGTTGGCGCGCG<br>GCAATCTGTGCGTAATCACACATGCCGTCAGTAGTCGAATCTTGTTAGAAGG<br>CCAGCGAGCAACCGGAATCGAGTATCTTTTAGGCAACGAAACACGTCAGGT<br>TAAAGCTAGACGTGAAGTTATTCTAAGTGCCGGTGCCTTTGGATCACCTCAG<br>TTACTGCAGTTGTCAGGTATAGGCCACGCTGAGGAATTACAGGCTCTCGGTA<br>TTCGAGCCGCCGTAAACCTCCCGGGGGTAGGCAAAAATCTGCAGGACCATA<br>TAGATTACGTTCAAACCTGGAGGACACCGTCTGATTCAGACACGTTCGGGGT<br>AAGTCTGCGGGGTACAGCACGCCTGACTTCTGCGATATTTGAATGGAGAAAT<br>AAGAGATCCGGTATGATAACATCGAATTTTACTGAAGCGGGAGCCTTTCTGT<br>GCAGCAGTCCGGAAGTTACCGTACCCGATCTGCAGCTGCATTTTGTGATCGG<br>TATAGTTGACGATCACGCGCGCAAGTTACACCTTGGCCATGGTATGAGTTGC<br>CATGTTTCTGTCATGCGCCCGTTTTCGCGTGGGACCGTTGGGCTGCGGAACA<br>CAGATCCGAGGGCCGCGCCGCTGATTGATCCGCGTTTTTTCGATGATGAGCG<br>TGATTTCCAGCTGTTGTTGAAGGGGGGACAACTCCAGCAACGAATCTTCGAA<br>AGTCGGCCGTTTGATGGCGTCCGTGGAAAAATGCTATATCCAGTCGACATCA<br>CAGATACAGCACGTATGGCTCAGGATATTCGCGCGAGGGCAGATACCCAAT<br>ATCATCCTTCATGTACTTGTAAGATGGGTCCGTCCACCGATGCTCAGGCGGT<br>CGTCGATGCACGCCTGAGAGTTCATGGTGTGGCCGGTTTACGGGTGGCGGAT<br>GCCAGCATCTTCCCTACTGTAACGGGTGGCAATACAAATGCACCGACTATCA<br>TGATCGGCGAAAAACTTTCACAGATGTTAAGGGAAGAAACAGTTACGAGAA<br>GTACTACGCGATAA | MEFDYVIVGAGSAGCVLA<br>SRLSEDPGITVCLLEAGGP<br>DKSVLIHAPAGAAAIIPSK<br>MNNWAFETVPQKGLNGR<br>MGYQPRGKTLGGSSSINA<br>MVYVRGNRWDYDHWAA<br>LGNPGWSYDDVLPFFKRA<br>ENNEQLRNEWHGQGGPL<br>NVTYPRHNSPLNRMFLDA<br>AAMNGLPLNPDYNGAGQ<br>HGAFMYQVTQINGERCSA<br>AKAYLTPQLARGNLCVIT<br>HAVSSRILLEGQRATGIEY<br>LLGNETRQVKARREVILS<br>AGAFGSPQLLQLSGIGHA<br>EELQALGIRAAVNLPGVG<br>KNLQDHIDYVQTWRTPSD<br>SDTFGVSLRGTARLTSAIF<br>EWRNKRSGMITSNFTEAG<br>AFLCSSPEVTVPDLQLHFV<br>IGIVDDHARKLHLGHGMS<br>CHVSVMRPFSRGTVGLRN<br>TDPRAAPLIDPRFFDDERD<br>FQLLLKGGQLQQRIFESRP<br>FDGVRGKMLYPVDITDTA<br>RMAQDIRARADTQYHPSC<br>TCKMGPSTDAQAVVDAR<br>LRVHGVAGLRVADASIFP<br>TVTGGNTNAPTIMIGEKLS<br>QMLREETVTRSTTR |
| pZE-RhoAAO<br>(MCB1395394.<br>1) | ATGTCGTACGATTTTGTGATCGTTGGCGGAGGGTCCGCAGGTGCCACCCTGG<br>CGGCCCGTCTTACCGAAGACCCACGACTAAGAGTTTGCCTGCTGGAAGCAG<br>GAGGCGCTGGCAAAGATATCTTGATCAGGGCTCCCCTGGGTGTAGTCGCAAT<br>GCTTCCGGGTTATGGTAAAATAAATAATTGGGCCTTTAAGACAGTTCCACAG<br>CCGGGTCTGAATGGTCGTCGCGGCTATCAACCTCGCGGTAGAGCGCTGGGG<br>GGCTCCAGCGCGATCAATGCGATGCTCTACCTTCGGGGTCAACGGCAAGATT<br>ACGATAGCTGGGCCGATTTAGGCTGTACGGGATGGGGTTGGGACGATGTCTT<br>GCCGTATTTTCGGAAGGCTGAAAATAACGTCAGGGGTGCCAACGCGGCACA<br>TGGCGATAGCGGACCGTTACAGGTATCAAACCAAAAAGCACCCCGACCCAT<br>TTCGGAAGCATTCATAGAAGCCTCGGGCGAGATGCAGATCCGCCGTACCGA<br>AGATTTTAATGCTGGAGATAATGAGGGTGCCGGGTACTTTCAATGTACACAG<br>TTCCATTCTCCTGATAAAAATGGTGAGCGCTGTAGCGCGGCGGCAGGCTATT<br>TACATCCGGTAATGGATCAGCGCCCGAATCTGACGGTAATAACAAAGGCGC<br>GTGCCACGCGGATCCTATTTGAAGGAAAACGGGCCGTTGGTGTGGAATACC<br>GGCAGGGAGGCGCCGTTAAACAGGTTAGAGCGGGCAAGGAGGTAATACTCA<br>GTGCGGGCGCTTTCCAGTCGCCGCAAATCCTTATGCTAAGCGGTGTCGGCTC<br>TGCTGAAGCACTCCGCCCACATGGTATAGCCCAGCTGCACGAACTGCCGGG<br>CGTCGGACAAAATCTGCAGGACCATATTGACTTCATCATGGGCTACAAAACC<br>CGAGATACAGATACAATCGGCTTAGGTCTGCGTGCGGGCATTAAACTCCTGG<br>GTGAGATGCTCAAGTGGAGAAAAGACGGAAATAGTATGGTAGCTTCTACAA<br>TCGCAGAAACAGGGTCTTTTTTTCGCACGGATCCTTCCCTGGACCGTCCAGA<br>TGTCCAAACTCATTTTGTCATCTCCGTTGTAGATGACCACGCGCGTAAGCTG<br>CATTATGGACATGGCTATAGTTGTCATGTTTGCGTGCTGCGTCCGCACTCCA<br>GGGGGGAAGTTTTCTTGCAATCTGCCGACCCAATGGCTGACCCCGGGATTGA<br>CCCGCGCTTCCTAAGCGACGAAAGGGATTTAAAAACCCTGATACGTGGCGC<br>TCGCATGACCCGTGATGTACTGGAAGCTCCCGCGCTGGCCAAATATCGACAC<br>AAGGAACTGTTTGGTATTCGGGACGGCATGTCTGACGCCGAATGGGAACAA | MSYDFVIVGGGSAGATLA<br>ARLTEDPRLRVCLLEAGG<br>AGKDILIRAPLGVVAMLP<br>GYGKINNWAFKTVPQPGL<br>NGRRGYQPRGRALGGSSA<br>INAMLYLRGQRQDYDSW<br>ADLGCTGWGWDDVLPYF<br>RKAENNVRGANAAHGDS<br>GPLQVSNQKAPRPISEAFI<br>EASGEMQIRRTEDFNAGD<br>NEGAGYFQCTQFHSPDKN<br>GERCSAAAGYLHPVMDQ<br>RPNLTVITKARATRILFEG<br>KRAVGVEYRQGGAVKQV<br>RAGKEVILSAGAFQSPQIL<br>MLSGVGSAEALRPHGIAQ<br>LHELPGVGQNLQDHIDFI<br>MGYKTRDTDTIGLGLRAG<br>IKLLGEMLKWRKDGNSM<br>VASTIAETGSFFRTDPSLD<br>RPDVQTHFVISVVDDHAR<br>KLHYGHGYSCHVCVLRP<br>HSRGEVFLQSADPMADPG<br>IDPRFLSDERDLKTLIRGA<br>RMTRDVLEAPALAKYRH<br>KELFGIRDGMSDAEWEQV<br>IRNRADTVYHPVGTCKM |

| | | |
|---|---|---|
| | GTCATTCGTAACCGTGCAGATACTGTTTACCACCCTGTGGGGACGTGTAAAATGGGAACAGATGACATGGCCGTGGTTACTCCGGAACTCAAGGTCCGCGGTTTAGAGGGTCTTCGTGTTGCAGATGCATCGATTATGCCGTTACTTGTTTCGGGAAATACCAACGCTCCGACCATTATGATTGGTGAGAAGTGTGCTGATATGATTAAAGCAGACCATGCCGCTTAG | GTDDMAVVTPELKVRGLEGLRVADASIMPLLVSGNTNAPTIMIGEKCADMIKADHAA |
| pZE-RiAAO (WP_149777140.1) | ATGCAGTTTGATTATGTCATCGTGGGTGGGGGGAGTGCTGGCTGCGTTTTGGCAAATCGCCTCTCAGCCAACCCAGGCGCCAGGGTTTGTCTGCTCGAAGCAGGCGGAGGTGGTAATGGCATTTTAGTACGCATGCCGGCAGGCGTCGTTGCCATGTTGCCGGGGCGCCCAAAGATCAACAATTGGGCATTTGAAACGGTTCCCCAGCCTGGCCTGAACGGCAGGAAAGGTTATCAACCTAGAGGTCGCGCATTGGGTGGTTCCTCAGCCATTAACGCCATGTTATATGTTCGCGGACAGCGCCAGGATTATGATGGCTGGGCTGATTTAGGGTGTGAAGGTTGGGACTGGGATTCTGTGCTGCCGTATTTCAAACGTTCAGAGAATAATGAAAGAGGCGCTGATGATTTGCATGGAGCCGATGGTCCGCTGCAGGTCTCGGATCAAAAGGAGGAACGTCCGATAACTCGGGCATTTGTTGAAGCAGCTGCCCAGCTGCAGCATAAGGTGACAGATGATTTCAATCGTGGTGACAATGAGGGTGCAGGGTTGTACCAAGTGACCCAGTTTCATGATCCGGCAAAAAACGGCGAACGCTGTAGCGCTGCAGCCGCGTATCTGTTCCCAGTAATGGACCGTCCAAATCTTACTGTAATTACAGGCGCTCAGGCGAGAGAAATTACTTTCGATGGCCATCGTGCGACGGGCGTCATCTACCGTCGCGGTGGTAAAGGGGCCGATCTGACAGTGACCGCTGCACGCGAGGTCCTTGTCTGCTCCGGGGCGTTAAAATCACCCCAGCTTCTCCAGATGTCAGGTATTGGAGATCCAGAAGATCTGACCCCCCATGGCATTGCAGTACGTCATGCGCTGCCAGGGGTCGGTAAGAACCTGCAGGACCATCTGGATTTTATATTAGCCTATAAAACAAAAGACACGGATAACTTCGGAATCGGTGCAGCAGGCACTGTTGGCCTAATAAAACACCTCCTAAGATGGCGGAAAACGGGCGTTTCTATGGCGGCTACTCCATTCGCGGAAGGCGCTGCATTTTTAAAAACGAGCCCAGATCTGGATCGTCCTGATGTGCAACTTCATTTTACGATAGCATTAGTTGATGATCATGCGCGTAAACTCCACCTGGGCTATGGCTTTTCTTGTCATATCTGCAAACTGAGACCCGAATCGCGTGGCACTGTAAGCCTGCATTCAGCTGATCCTTTCGCAGCACCGGCCATTGATCCGGCTTTCCTAAGCGATCCGCGTGATCTAGATACAATGATCAAAGGCGCTCGTATGACTCGCGAAATTTTGGAAGCCCCGGCGCTAGCGAAGTACAGACACAAGGAGATGTTTGGCACTGATACGGCGCGAACCGATGCGGATTGGGAACGACATATACGTGCACGTGCGGATACAATTTACCATCCCGTTGGTACTTGTAAAATGGGCGTTGATGATATGGCAGTAGTCGACCCGCAACTGCGTGTGCGTGGTTTGCAGGGACTCCGTGTGGTTGATGCAAGCGTTATGCCTACCCTGGTGTCCGGCAACACCAACGCCCCGACCATAATGATCGCCGAAAAAGCCGCGGACATGATCTTGGCTGATGTCGCGTAG | MQFDYVIVGGGSAGCVLANRLSANPGARVCLLEAGGGGNGILVRMPAGVVAMLPGRPKINNWAFETVPQPGLNGRKGYQPRGRALGGSSAINAMLYVRGQRQDYDGWADLGCEGWDWDSVLPYFKRSENNERGADDLHGADGPLQVSDQKEERPITRAFVEAAAQLQHKVTDDFNRGDNEGAGLYQVTQFHDPAKNGERCSAAAAYLFPVMDRPNLTVITGAQAREITFDGHRATGVIYRRGGKGADLTVTAAREVLVCSGALKSPQLLQMSGIGDPEDLTPHGIAVRHALPGVGKNLQDHLDFILAYKTKDTDNFGIGAAGTVGLIKHLLRWRKTGVSMAATPFAEGAAFLKTSPDLDRPDVQLHFTIALVDDHARKLHLGYGFSCHICKLRPESRGTVSLHSADPFAAPAIDPAFLSDPRDLDTMIKGARMTREILEAPALAKYRHKEMFGTDTARTDADWERHIRARADTIYHPVGTCKMGVDDMAVVDPQLRVRGLQGLRVVDASVMPTLVSGNTNAPTIMIAEKAADMILADVA |
| pZE-RuAAO (WP_209940099.1) | ATGCAGTTCGATTATGTAATTGTTGGAGGTGGCTCTGCGGGATGTGTGCTGGCAAATCGCCTGTCGGAAGATCCGGCTACCAGGGTGTGCCTCTTGGAGGCGGGTGGATCCGGGGATGACGTGGTGGTCAGGATGCCCAGCGCTGCAGTGGCCACGCTACCCGGGCGCCCGCGTATACATAACTGGGCCTTTGAGACAGTTCCACAAACGGGCCTAAACGGTCGGAAGGGTTATCAACCCCGAGGAAAGACCCTGGGTGGAAGTTCGGCTATCAATGCAATGTTATACGTAAGAGGACAAAGGCAGGACTATGACGATTGGGCCGATATGGGTTGTGAGGGCTGGGCCTGGAATGACGTACTCCCGTATTTCAAACGCAGTGAGAACAACGAGCAGGGCGCAGACGATATGCATGGTGGCTGTGGTCCACTACATGTGTCTCACCAGAAAGAGCCTCGCCCTATCACGCGTGCTTTCGTGGAGGCCGCAGGCCAATTGCAACATCGTCGCACCTCCGACTTTAATCGCGGTGATAATGAAGGCGTCGGTCTGTATCAGGTAACTCAATTTCATGGTCCGGCAAAGAATGGGGAGCGTTGTTCAGCAGCCTCCGCATTTCTCATACCTGCTATGGATCGACCGAACCTCACCGTAATCACTCACGCAGTTGTCAAAGAATTGACGTTTGAAGGTAAACGGGCCTCTGGTGTGACCTATCGTACCAAAGGAAAAGGCGCCGATATAAGTGTTAAAGCAACCCGTGAAGTGCTGCTATGTGCTGGTGCTCTTAAATCTCCGCAGCTGCTTCAACTGTCGGGCATTGGAGCGGGATCAGATCTGAATCCGCACGGGATCGCCATCCGACATGAACTGCCGGGTGTTGGGCAGAACCTTCAGGACCATCTTGATTTCATTCTGACGTATAGAACAAAAGATACGAATACCTTTGGTATTGGTCCAGTTGGGGCTGTTCGTCTTCTGGGGCACATCCAACGTTGGTACAAAACGGGCGCATCCATGGCAGCTACTCCGTTCGCTGAAGGCGCGGCTTTTTTAAAAACTAGTCCAGATCTCGAACGTCCTGACATTCAGTTGCACTTTACGATCGCAATGGTCGATGATCACGCGCGTAAGTTGCATTATGGTTACGGTTATGGTTGTCATGTTTGCAAATTAAGACCGGATTCTCGCGGCGCCGTAGGACTGCGCAGTGCCGACCCAATGGATACCCCTGCCATTGATCCGGCTTTTCTGTCAGACGCCCGTGATCTTGAGACTATGATTAAGGGGGCGCGGATGACTCGGGATATTTTGGAAGCCCCGGCATTGGCGAAATATAGACATAAGGAAATGTTTGGAACCGATACCGCTCAAACCGACGCTGACTGGGAAAAACACATTCGGGCCCGGGCAGATACCATTTACCACCCAGTTGGCACTTGTAAAATGGGGCGCGATCCAATGGCCGTGGTAGACCCGCAATTGCGTGTCCACGGCCTGGAAGGTCTCAGAGTAGTGGATGCCTCTATCATGCCGAAACTTGTAAGTGGTAACACGAACGCTCCCACTATCATGATTGCCGAAAAAGCTGCTGACATGATCCGCAGTGCGGCCGGCTGA | MQFDYVIVGGGSAGCVLANRLSEDPATRVCLLEAGGSGDDVVVRMPSAAVATLPGRPRIHNWAFETVPQTGLNGRKGYQPRGKTLGGSSAINAMLYVRGQRQDYDDWADMGCEGWAWNDVLPYFKRSENNEQGADDMHGGCGPLHVSHQKEPRPITRAFVEAAGQLQHRRTSDFNRGDNEGVGLYQVTQFHGPAKNGERCSAASAFLIPAMDRPNLTVITHAVVKELTFEGKRASGVTYRTKGKGADISVKATREVLLCAGALKSPQLLQLSGIGAGSDLNPHGIAIRHELPGVGQNLQDHLDFILTYRTKDTNTFGIGPVGAVRLLGHIQRWYKTGASMAATPFAEGAAFLKTSPDLERPDIQLHFTIAMVDDHARKLHYGYGYGCHVCKLRPDSRGAVGLRSADPMDTPAIDPAFLSDARDLETMIKGARMTRDILEAPALAKYRHKEMFGTDTAQTDADWEKHIRARADTIYHPVGTCKMGRDPMAVVDPQLRVHGLEGLRVVDASIMPKLVSGNTNAPTIMIAEKAADMIRSAAG |

| | | |
|---|---|---|
| pZE-HsAAO<br>(WP_21666600<br>1.1) | ATGTTTGATTACGTGATTGTTGGCGGTGGATCTGCGGGTGCCACTTTAGCTG<br>CTCGTCTGTCAGAAGATCCGCGCATTTCAGTTTGCCTGCTTGAGGCCGGTGG<br>ACAGGGAGACTCGCTTTTAGTGCGCACTCCTGCAGCTGTAGTCGCAATGCTT<br>CCAGGTTATGGGAAATTAAACAATTGGGCGTTACAAACTACGCCGCAACCT<br>GGACTAAACGGGCGTCGGGGATACCAGCCACGAGGCCGTGCATTGGGAGGT<br>AGCTCTGCTATAAATGCAATGCTGTATGTACGGGGACAGCGACAAGACTAT<br>GACGGTTGGGCGCAAGCTGGATGCCCTGGTTGGGATTGGGAGTCGGTCCTTC<br>CGTATTTCAAAAGAGCGGAAAATAATGTTCGGGGCGCGGATGCGTGGCACG<br>GGGCGAGTGGCCCGCTTCAAGTGAGTGAACAGAACCGGCCACGCCCTATCA<br>CCAGAGCTTTTATTGAGGCGGGCCAAGCGCGTGGTCATCGGCTGTGTCAAGA<br>CTTTAATACCGGTGACAATGAAGGTGTTGGTTTGTATCAAGTAACCCAGTTT<br>CACACACCGGCACATCGTGGCGAGCGTTGTAGTGCGGCAGCTGCCTATTTGC<br>ATCCGGTGATGGGCCAACGCCCCAACCTGTCAGTATTACGGCAGGTACGGG<br>CCCAGCGCCTCTTGTTTGAGGGAAAACGTGCCATCGGTGTGGCTTATCGTCA<br>ACAACAGGGTGATGCGCAGGTTCGCGCTGCACGTGAAGTCATTATTGCCGC<br>AGGGGCCTTCGGATCACCGCAGCTTCTCCAGCTGTCTGGTGTTGGGCGCTCG<br>GATGACATCTTACCTCATGGCATTGCACTGCATCATCCACTCCCGGGTGTCG<br>GCCAGAATCTGCAAGATCATCTAGATTTCACTCAGGGTTGGACCACCCGGGA<br>TACCGACAATTTTGGGCTTGGTGTTGTGGGTGGCCTGCGCCTGTTGGGACAG<br>CTGCTGCCTTGGAAGCGCCACGGTGAGGGTCTCATTGCAACCCCTTTCGCCG<br>AAGGCGCAGCTTTTCTGAAGACGCGTCCGGAACTAGACCGCCCCGACATCC<br>AATTACACTTCTGTATCGCAATAGTAGATGATCACGCACGCAAGTTGCACGC<br>CGGCTATGGTTTCTCATTACATATGTGTATGCTGCGCCCGCACTCCCGTGGCC<br>GAGTTGGCTTACAGTCAGCTGATCCTATGGCTGATCCCCTTATCGATCCAGG<br>GTATCTGAGTGATCCGCGCGATCTAGCTACTATGATTGATGGCGCAAGAATG<br>GCTCGCCAGATAGTTATGACAGAACCCTTGCGCCACTACTGTAGACGTGAAC<br>TGTTCGGTGGTAGGGATGACATGGACGATGCCCAATGGGAATCCATGATTC<br>GTCATCGCGCCGATACCATTTATCATCCAGCAGGTACTTGTAGGATGGGCGA<br>AGACGCGATGGCGGTGGTTGACGCACAGTTGCGCGTTCACGGCCTGCAGGG<br>TCTGCGCGTTGTTGACGCAAGCGTGATGCCTACACTGGTATCAGGCAACACC<br>AATGCGCCAACTATTATGATCGCCGAAAAGGCCGCGGATATGATCCGGGCC<br>TCTCACGAGCAGGCGCGCGCCGCCTGA | MFDYVIVGGGSAGATLA<br>ARLSEDPRISVCLLEAGGQ<br>GDSLLVRTPAAVVAMLPG<br>YGKLNNWALQTTPQPGL<br>NGRRGYQPRGRALGGSSA<br>INAMLYVRGQRQDYDGW<br>AQAGCPGWDWESVLPYF<br>KRAENNVRGADAWHGAS<br>GPLQVSEQNRPRPITRAFI<br>EAGQARGHRLCQDFNTG<br>DNEGVGLYQVTQFHTPA<br>HRGERCSAAAAYLHPVM<br>GQRPNLSVLRQVRAQRLL<br>FEGKRAIGVAYRQQQGD<br>AQVRAAREVIIAAGAFGS<br>PQLLQLSGVGRSDDILPHG<br>IALHHPLPGVGQNLQDHL<br>DFTQGWTTRDTDNFGLG<br>VVGGLRLLGQLLPWKRH<br>GEGLIATPFAEGAAFLKTR<br>PELDRPDIQLHFCIAIVDD<br>HARKLHAGYGFSLHMCM<br>LRPHSRGRVGLQSADPMA<br>DPLIDPGYLSDPRDLATMI<br>DGARMARQIVMTEPLRH<br>YCRRELFGGRDDMDDAQ<br>WESMIRHRADTIYHPAGT<br>CRMGEDAMAVVDAQLR<br>VHGLQGLRVVDASVMPT<br>LVSGNTNAPTIMIAEKAA<br>DMIRASHEQARAA |
| pZE-JtAAO<br>(WP_10400743<br>5.1) | ATGAGTTATGATTTTGTAATAGTTGGAGGTGGTTCCGCGGGAGCTACCCTGG<br>CAGCCCGGCTCTCAGAAGATCCTGCAGTTAAGGTCTGCCTCCTGGAGGCTGG<br>AGGCGGTGGTCGAGACATTCTGATCCGTGCACCGATTGGGGTGGTGGCGAT<br>GCTGCCGGGACATGGTAAAATTAACAACTGGGCTTTTGAGACCGTGCCGCA<br>GCCAGGCTTGAATGGTCGCAAGGGCTATCAGCCGAGAGGCCGTGCACTGGG<br>TGGTTCGTCTGCCATTAATGCAATGCTGTATGTTCGGGGCCAACGCCAGGAT<br>TATGATGGCTGGGCTGAAGCGGGCTGCGATGGGTGGTCGTGGGACGAAGTA<br>CTACCATATTTCCGTAAGGCTGAGAACAATGTCAGAGGTGAGAATGAATTCC<br>ATGGAGCCAGCGGGCCTTTGCATGTTTCCGACCAGAAAGCTCCGCGACCCAT<br>TAGTGAAGCATTTCTTGAAGCTAATGCGCAGATGCAAATTCGTAGAGTTGAT<br>GATTTTAATACGGGGGATAACGAAGGAGCATCACTCTACCAGTGCACACAG<br>TTTCATGATCCGGAACGTAATGGAGAAAGATGCTCCGCAGCAGCGGCATAC<br>TTGTTCCCGGTCATGGATAAACGTCCCAATTTAACTGTTATTACTAAGGCAC<br>GTACTACCAGAATTTTATTCGAGGGAAAAAGGGTTGTGGGGGTGGAATATA<br>GAGTCGGCAAACAAACACAGCGTGCGATGGCGGCACGCGAAGTTATTCTCA<br>GCGCAGGCGCATTTCAATCACCGCAGATTCTGATGCTGTCTGGCGTTGGTAG<br>AGAGCAAGATATACGACCGCATGGCATAGAAATGGTACATGAATTGCCGGG<br>AGTCGGGCAGAATTTACAAGACCATATTGATTTTGTCCTAGCGTGGAAAACC<br>AAAGATACTGACAACATTGGAATTGGTTTACGCGCTACCGCGAAACTCACCT<br>CCGAAATTTTCAAATGGCGTAAACACGGGAATTCAATGGTGGCCTCAACGA<br>TTGCCGAAGCGGGTAGTTTTTTCAAGACCGATCCCAACCTGGACCGACCTGA<br>TGTGCAGACCCATTTTGTTATTAGTATTGTAGATGATCACGCGCGTAAGCTC<br>CACTTGGGACACGGCTATTCATGCCATGTATGCGTCCTGCGTCCACACAGCC<br>GTGGGGAAGTGTTCTTGCAGAGTGCTGACCCTATGGCTGCACCGGGTATCGA<br>CCCAAAATTTTTGAGCGACGAACGCGACCTCAAGACGCTTGTTAAAGGGGC<br>CAAAATGACCAGACAGATCATGACCGCGCCACCGATGCGTCCTTATATCCAC<br>AAAGAATTATTTGGTGTTCACGACGATATGACAGATGCTGAATGGGAGGCT<br>CATATTCGCGCACGGGCAGATACCGTTTATCATCCGGTGGGCACGTGCAAGA<br>TGGGAGTCGATGATATGGCAGTAGTTGACCCGGCGTTGAAGGTTCGTGGGCT<br>TGAAGGATTACGGGTGGTGGATGCATCGGTGATGCCGACTTTGGTATCTGGT<br>AATACCAATGCGCCGACAATTATGATCGCGGAGAAAGCAAGTGATTTGATC<br>CGCGCGGAATACGGCGCCCAGTCAGCGGCTGCAGAATAA | MSYDFVIVGGGSAGATLA<br>ARLSEDPAVKVCLLEAGG<br>GGRDILIRAPIGVVAMLPG<br>HGKINNWAFETVPQPGLN<br>GRKGYQPRGRALGGSSAI<br>NAMLYVRGQRQDYDGW<br>AEAGCDGWSWDEVLPYF<br>RKAENNVRGENEFHGAS<br>GPLHVSDQKAPRPISEAFL<br>EANAQMQIRRVDDFNTG<br>DNEGASLYQCTQFHDPER<br>NGERCSAAAAYLFPVMD<br>KRPNLTVITKARTTRILFE<br>GKRVVGVEYRVGKQTQR<br>AMAAREVILSAGAFQSPQI<br>LMLSGVGREQDIRPHGIE<br>MVHELPGVGQNLQDHIDF<br>VLAWKTKDTDNIGIGLRA<br>TAKLTSEIFKWRKHGNSM<br>VASTIAEAGSFFKTDPNLD<br>RPDVQTHFVISIVDDHAR<br>KLHLGHGYSCHVCVLRPH<br>SRGEVFLQSADPMAAPGI<br>DPKFLSDERDLKTLVKGA<br>KMTRQIMTAPPMRPYIHK<br>ELFGVHDDMTDAEWEAH<br>IRARADTVYHPVGTCKM<br>GVDDMAVVDPALKVRGL<br>EGLRVVDASVMPTLVSGN<br>TNAPTIMIAEKASDLIRAE<br>YGAQSAAAE |
| pZE-Bb1AAO<br>(MDG1108522.<br>1) | ATGAAATTCGATTTTGTCATTGTAGGTGGGGGATCAGCGGGCTCTACTCTGG<br>CCGCGCGTCTCACAGAGGATCCGGATGTAACCGTCTGTCTGCTTGAGGCCGG<br>CGGTGGAGCCGACCCCATTCTGCTGAGAGCTCCTGCAGCCGTTGTGGCTCTG<br>TTACCGGGTTATGGCAAACTGTATAACTGGGCTTTTCAGACAGTTCCTCAAC<br>CAGGGTTAAATGGTCGGCGCGGCTATCAACCCCGGGGACGCGCACTGGGGG<br>GGAGCTCAAGCATCAACGCGATGCTCTATGTCCGTGGTCAGAAATCTGATTA | MKFDFVIVGGGSAGSTLA<br>ARLTEDPDVTVCLLEAGG<br>GADPILLRAPAAVVALLP<br>GYGKLYNWAFQTVPQPG<br>LNGRRGYQPRGRALGGSS<br>SINAMLYVRGQKSDYDG |

| | | |
|---|---|---|
| | TGATGGCTGGGCGGATATGGGTTGCAATGGCTGGTCTTGGGACGACTGTCTC<br>CCGTATTTTATTCGGAGTGAAAACAATGCGCGGGGGGCTTCTGCGTGGCATG<br>GCGATAGTGGACCCTTGCACGTCTCCGATCAGCAATCGCCGCGGCCTATTTC<br>CAAAGCTTTCGTGGAAGCTGCAACCCAGATGGGGCATGCCGCCGTTGATGA<br>TTTCAATACCGGTGAAAATGATGGTGCCGGTTTGTTTCAGGTGACCCAGTTT<br>CACGACGAAGCGAAAAATGGTGAACGCTGTTCTGCAGCCCTGGGTTACTTGT<br>ACCCAGCCCTGTCACGCCCCAATCTGACCGTTATTACTAAAGCTCGTGCGAC<br>GAAAATCCTGTTCGAGGGCCAACGCGCGGTTGGGGTTAATTATCGGTCAGG<br>CGGCGAGGATAAATCAGCCTACGCAGCGAAAGAAGTTATCCTATGCGGCGG<br>CGCTTTTCAAAGCCCGCAGCTCCTGCAGTTAAGCGGAGTTGGTCGCAGCGCA<br>GACATTACGCGATTTGGCATCGAGATGGTGCATGAACTGCCGGGTGTAGGTC<br>AAAACTTACAAGATCATCTGGATTTTATCTTAGCTTGTAAAACCGACGACAA<br>AGATAACTTTGGCATCGGTGTTGCTGCGTCCATCCAACTGGTGAAACATATA<br>CTGCAGTGGAGACAAGATGGCACAGGGATGGTTGCAACTCCGTTTGCAGAG<br>GGTGGTGCGTTTCTAAAATCTGATCCTTCCGTGGAAAGGCCAGACATCCAAT<br>TACATTTTGTGATAGCTGTTGTAGATAACCATGGAAAAAATCTGCACCGTGG<br>ATACGGTTATAGTTGCCATGTGTGCCCGGTGCGCCCACATAGTCGTGGCGAA<br>GTATTCTTAACGAGTGGTGACCCTTTTGCTGATCCCGGCATTGATCCGAAGT<br>TTCTCAGCGATCCGCGTGATCTCGCCGTTATGGTTAAAGGTGCAAAAATAGC<br>TCGGGATATTCTTGAAGCTCCTGCCATGAAACTCTACAAAAAACATGATTTA<br>TTTGGCGTTACCGATAACTTGTCTGATGCTGCCTGGGAAGAACACATCCGGG<br>CGCGGAGCGACACCATATACCACCCGGTTGGCACGTGCAAAATGGGCGTTG<br>ATGAAATGGCGGTCGTTGATCCTGAACTGCGTGTTCATGGTTTGAGCGGTTT<br>ACGTGTAGTTGACGCGAGCGTAATGCCTACAGTTGTTTCGGGCAATACAAAT<br>GCGCCGACTATCATGATTGCCGAACGTGCGGCGGACATGATTCGCAGTGCAT<br>ACGCTAAAGATAATGCCATATTAGAAAGCTCAAGTTAA | WADMGCNGWSWDDCLP<br>YFIRSENNARGASAWHGD<br>SGPLHVSDQQSPRPISKAF<br>VEAATQMGHAAVDDFNT<br>GENDGAGLFQVTQFHDE<br>AKNGERCSAALGYLYPAL<br>SRPNLTVITKARATKILFE<br>GQRAVGVNYRSGGEDKS<br>AYAAKEVILCGGAFQSPQ<br>LLQLSGVGRSADITRFGIE<br>MVHELPGVGQNLQDHLD<br>FILACKTDDKDNFGIGVA<br>ASIQLVKHILQWRQDGTG<br>MVATPFAEGGAFLKSDPS<br>VERPDIQLHFVIAVVDNH<br>GKNLHRGYGYSCHVCPV<br>RPHSRGEVFLTSGDPFADP<br>GIDPKFLSDPRDLAVMVK<br>GAKIARDILEAPAMKLYK<br>KHDLFGVTDNLSDAAWE<br>EHIRARSDTIYHPVGTCK<br>MGVDEMAVVDPELRVHG<br>LSGLRVVDASVMPTVVSG<br>NTNAPTIMIAERAADMIRS<br>AYAKDNAILESSS |
| pZE-OgAAO (WP_00742780 9.1) | ATGGAATTTGATTATGTCATTGTTGGCGGAGGTAGTGCGGGAGCCGTGCTCG<br>CTGCGCGTCTGTCTGAAGATCCGGCAACCTCCGTTTGCTTACTGGAGGCCGG<br>GGGTGAAGGCCGCCATCTCCTGATTCGCGCCCCTGCGGCAGTCGTCGCAATG<br>ATGCCCGGCCATGGTCGTATAAGCAATTGGGCGTTTAAAACTGTTCCTCAGC<br>CGGGCTTGAACGGTCGCCGCGGATATCAGCCTCGGGGTAAGGGGTTGGGTG<br>GGTCGAGCGCGATTAACGCTATGTTGTATATTCGAGGCCACAGAAGCGATTA<br>TGATGATTGGGCAGAATCAGGCTTAGACGGATGGGGTTGGGATGACGTTCT<br>GCCATACTTTATACGTTCGGAAGGCAATGCGAGCGGTGCTGACGATGCGCAT<br>GGGGCCGATGGACCCCTCCAGGTGCGAGATCAGCCCCACCCACGTGCAATC<br>AGCCGGGCATTCGTGGAGGCAGGCACACAGCTGCAGCACCGAGCGGTAGCC<br>GACTTCAATCGTGGCGACAATGAAGGTATCGGCCTGTATCAGGTGACACAA<br>TTTCATTCCGGTCCCCGGCGAGGAGAACGTTGTTCTGCAGCAGCGGCGTATC<br>TGCATCCTGTAATGGGTAGGCGAGCCAACCTGCATGTCGAAACAAGGGCGC<br>ATGCCACCCGCATTGTACTGGCTGGACGGCGTGCAACGGGCGTTGCATGGC<br>GCAAAGGCCGCCGTGGAGAAGAAAGGGTTGTTCGCGCGCGTCGTGAGGTCA<br>TACTTTCAGCGGGCGCTTTCGGCAGCCCGCAATTACTGCAGCTGTCCGGTAT<br>CGGTAGAGGTCAAGATATTCGGGCTCATGGCATAGCTGTTGCACATGATTTA<br>CCTGGTGTCGGACAAAATTTGCAAGACCACCTGGATTTTATTTTGGCTTGTA<br>AAACTCGTGATACCGATAATTTCGGTATTGGAGCGCGGGCAACCTGGGGAC<br>TGATGAGGCATGCTTGGGCCTGGCGTCGCGATGGTGGAGGCATGATCGCAA<br>CTCCTTTTGCTGAAGGCGCCGCCTTTTTAAAGACCCGGCCGGGGCTGGACCG<br>CCCGGACGTTCAACTGCATTTTGTTATCAGTATTGTGGATGATCATGCAAGA<br>AAACTGCATTTAGGCCACGGATATAGTTGCCACGTTTGTGCTCTCCGTCCCC<br>ATTCTCGCGGGCAAGTTGGTCTTCGCAGCGCAGATCCAATGGCCCCCCCCGC<br>AATTGACCCAAACTACTTATCGGATCCTCGGGACCTGGAGACAACGATTCGG<br>GGCGCCAAAATGACCAGAGCGATACTGCAAGCACCGGCACTGGCTAGATAT<br>TGTCGTACCGAATTGTTTGGGATTAGGGATGGGATGTCTGATGCCGACTGGG<br>AGGGGCACGTACGTGCGCGCGCGGATACAATCTATCACCCGGTTGGGACCT<br>GTCGGATGGGCCCCGCCGGGGATGCAGGTGCAGTAGTAGACGCAGCCCTGC<br>GGGTTAGAGGTATGGAAGGCCTCCGTGTGGTCGATGCAAGCGTAATGCCTA<br>CCCTGATTGGGGGTAACACGAACGCTCCTACAATTATGATCGCCGAGAAAG<br>CAGCAGATACCATTCGAGGTAGAGCTCCGCAAGGAGTCGCGGCAGAGTAA | MEFDYVIVGGGSAGAVL<br>AARLSEDPATSVCLLEAG<br>GEGRHLLIRAPAAVVAM<br>MPGHGRISNWAFKTVPQP<br>GLNGRRGYQPRGKGLGG<br>SSAINAMLYIRGHRSDYD<br>DWAESGLDGWGWDDVL<br>PYFIRSEGNASGADDAHG<br>ADGPLQVRDQPHPRAISR<br>AFVEAGTQLQHRAVADF<br>NRGDNEGIGLYQVTQFHS<br>GPRRGERCSAAAAYLHPV<br>MGRRANLHVETRAHATRI<br>VLAGRRATGVAWRKGRR<br>GEERVVRARREVILSAGA<br>FGSPQLLQLSGIGRGQDIR<br>AHGIAVAHDLPGVGQNL<br>QDHLDFILACKTRDTDNF<br>GIGARATWGLMRHAWA<br>WRRDGGGMIATPFAEGA<br>AFLKTRPGLDRPDVQLHF<br>VISIVDDHARKLHLGHGY<br>SCHVCALRPHSRGQVGLR<br>SADPMAPPAIDPNYLSDPR<br>DLETTIRGAKMTRAILQAP<br>ALARYCRTELFGIRDGMS<br>DADWEGHVRARADTIYH<br>PVGTCRMGPAGDAGAVV<br>DAALRVRGMEGLRVVDA<br>SVMPTLIGGNTNAPTIMIA<br>EKAADTIRGRAPQGVAAE |
| pZE-VsAAO (WP_17130862 4.1) | ATGGACTCATATGATTTCATTGTTGTAGGAGGTGGAAGTGCAGGTTGTGTGA<br>TGGCGTCTCGCCTGAGCGAAGATCCGAACGTTACGGTTTGCTTGCTGGAGGC<br>AGGTGGTAAAGATACCTCGCCGTTTATCCATACCCCTGTTGGGGTGGTTGCC<br>ATGATGCCTACCAAACTGAATAATTGGGCATTTGAAACAGTTAAACAGCCG<br>GGCTTAAATGGTCGCAAGGGCTACCAACCGCGAGGTAAGACACTGGGAGGA<br>AGCAGTTCAATCAATGCGATGATGTATGCACGCGGGCATCGGTACGATTATG<br>ATATTTGGGAAAGTCTGGGAAATACCGGTTGGGATTATAAGTCCTGTCTCCC<br>GTATTTTAAAAAAGCCGAGAACAACGAAGTGCATCAGGATGAGTACCATGG<br>CCAGGGGGGTCCCCTCAATGTCGCGAATCTTAGAAGCCCAAGCCCGATGCT<br>GGAACGCTACTTAACAGCCTGTGAGTCCATCGGTGTACCGCGCAATGAAGA<br>TATCAACGGCGCGGCCCAGTTTGGGGCAATGCCCACCCAGGTGACTCAGAT<br>CAACGGCGAAAGATGCAGCGCCGCGAAAGCATATCTGACTCCGAATCTCTC | MDSYDFIVVGGGSAGCV<br>MASRLSEDPNVTVCLLEA<br>GGKDTSPFIHTPVGVVAM<br>MPTKLNNWAFETVKQPG<br>LNGRKGYQPRGKTLGGSS<br>SINAMMYARGHRYDYDI<br>WESLGNTGWDYKSCLPY<br>FKKAENNEVHQDEYHGQ<br>GGPLNVANLRSPSPMLER<br>YLTACESIGVPRNEDINGA<br>AQFGAMPTQVTQINGERC<br>SAAKAYLTPNLSRPNLTV |

| | | |
|---|---|---|
| | CCGTCCTAATCTGACGGTTGTGACCAAGGCAACCACCCATAAGGTCCTTTTT<br>GAAGAAAAAAAAACTGTGGGCGTTGAATACGGAAGCAATGGAAAACGATA<br>TCAGATCCGTTCAAATAAAGAGGTGATTCTGTCTGCCGGGGCATTTGGCAGC<br>CCTCAACTGTTACTTTTATCAGGTGTTGGCGCTAAGGCCGAACTGGAAGCTT<br>TGGGTATTGAACAAGTGCATGAACTGCCGGGCGTAGGTAAAAATCTGCAGG<br>ACCATATCGACCTGGTGCACTCATATAAGTGCAGCGAAAAACGCGAAACCT<br>TTGGCATTTCACTTCAGATGGCTTCCGAAATGACAAAAGCACTGCCGTTATG<br>GCGCAAAGAACGACGTGGAAAGATGAGTTCGAATTTCGCTGAGGGTATCGG<br>TTTTTTATGTTCGGACGATCACATTAGCGTCCCGGACTTAGAGTTTGTCTTTG<br>TTGTGGCTGTCGTTGACGACCATGCTCGTAAGATTCATACGTCTCACGGATT<br>TACATCGCACGTAACATTATTACGGCCGAAAAGCCACGGCTCCGTTACCCTA<br>AATTCTAATGATCCGTATGATCCTCCGAAGATCGATCCGGCATTTTTCAGTC<br>ACCCAGAAGACATGGAAATTATGATTAAGGGCTGGAAAAAACAGTATCAAA<br>TGCTGGAATCCTCAGCGTTTGACGATATCCGCGGTAACGCATTCTATCCTGT<br>TGACGCAAGTGATGATAAGGCAATCGAACAAGATATTCGTAATAGAGCCGA<br>CACTCAATATCACCCTGTCGGTACCTGTAAAATGGGTCCTAGCTCGGATTCC<br>CTGGCAGTAGTGGACAATGACCTGAAGGTGTATGGCCTGGATAATCTGAGA<br>GTTATTGATGCTTCCGTAATGCCAACTCTCATAGGGGCCAATACTAATGCAC<br>CAACCATTATGATCGCCGAGAAAGTGGCCGACCAGATTAAAGAAGAATATG<br>GTTTAGGGCAGCAGGAATCGTCTATCCAGGCGAAAGGCGAAGCGGTGGTAT<br>GA | VTKATTHKVLFEEKKTVG<br>VEYGSNGKRYQIRSNKEV<br>ILSAGAFGSPQLLLLSGVG<br>AKAELEALGIEQVHELPG<br>VGKNLQDHIDLVHSYKCS<br>EKRETFGISLQMASEMTK<br>ALPLWRKERRGKMSSNF<br>AEGIGFLCSDDHISVPDLE<br>FVFVVAVVDDHARKIHTS<br>HGFTSHVTLLRPKSHGSV<br>TLNSNDPYDPPKIDPAFFS<br>HPEDMEIMIKGWKKQYQ<br>MLESSAFDDIRGNAFYPV<br>DASDDKAIEQDIRNRADT<br>QYHPVGTCKMGPSSDSLA<br>VVDNDLKVYGLDNLRVI<br>DASVMPTLIGANTNAPTI<br>MIAEKVADQIKEEYGLGQ<br>QESSIQAKGEAVV |
| pZE-Bb2AAO<br>(MCB2021549.<br>1) | ATGAACCATGACCTGGAGTTTGATTATGTAATAGTAGGTGCCGGTTCGGCAG<br>GGTGTGCAATTGCAGCACGGCTTACCGAGGATCCTGATGTGTCCGTCTGCCT<br>GTTAGAAGCCGGCGGTCCGGATAAGTCAGTGCTGATCCACGCCCCAGTCGG<br>GGTGGTTGCAATGTTGCCTACTAAAATAAACAACTATGCGTATCAGACTGTA<br>CCTCAACCTGGATTAAATGGGCGCCGCGGTTACCAACCGCGTGGGAAAACC<br>CTGGGCGGTAGCTCTTCAATTAACGCAATGTTATATGTTAGAGGTCACCGCC<br>ATGATTACGACCATTGGGCGGCTTTAGGAAACCCGGGTTGGAGCTATGATG<br>AAGTCCTGCCGCTTTTTCGTCGGTCGGAAAATAATGAGCACTTTCGCAATGA<br>TCCATACCATGGTAGTGGTGGACCATTGAACGTCGCCGAATTACGTAGTCCA<br>AGCGAGTTGGATGAGGCATTTGTTCAAGCCGCCGTAAACCATGGTATCCCGC<br>GCACCCCAGACTATAATGGCGCTGAACAGTTTGGGTCATTTCGCTATCAGGT<br>GACGCAGAAGAATGGAGAGCGATGCTCGGCTGCGAAAGGCTATCTTACCCC<br>GAACCTGAGCCGACCCAACCTGAAAGTTATAACCCACGCAGTTTCGGCTAG<br>AGTTCTGTTAGACGGTAGACGCGCCCGTGGAGTAGCCTATTACCAGGGCAAT<br>GAGCTGACGGAGGTGCGTGCTCGCAGAGAAGTGGTGCTGGCTGGAGGAGCA<br>TTTGGTTCGCCGCAGCTGCTGATGCTGAGTGGCATTGGGCCGGCAGAAGAGT<br>TACGTAAGCACGGTATTACTGTGGCGCATGCGCTTCCAGGCGTGGGTCAGAA<br>TCTGCAAGATCATATAGATTATGTTCAGACCTACATGGCATCTAGCCGTACG<br>GAAACCTTTGGTGTTAGCGCACGCGGAGGGGTACGGATGACAAAAGGAATT<br>CTGGAGTGGCGGCGGGAAAGAACGGGGCCAGTGACCTCTCCTTTTGCCGAA<br>GCTGGAGCTTTTTTCGTTCCAGCCCGGCAGTCCCGGTTCCAGATTTACAGCT<br>GGTATTTGTAGTGGCAATTGTCGATGACCATGCACGCAAAATGCATTTGGGG<br>CACGGACTGTCTTGCCATCTGACCTTACTCCGTCCAAAAAGTCGTGGTACCG<br>TTGGAATCGCTAGCTCAGATCCAAGAGAAGCACCCGTGATTGATCCGCGCTT<br>TTTCAGCGATCCGGCCGACATGCCACTGATGCTGGATGGTGCCGCAAAAATG<br>CAGGCTATCTTAGAAGATAGAGCACTGGCCCGCTATCGCAACGGGAAGATG<br>CTTTACACAGTCAGAGCCGATGATCGCTCTGGCCTTGAGGCTGATATTCGGA<br>ATCGGGCAGACACCCAGTATCACCCGGTTGGTTCATGTAAAATGGGCCCGG<br>CGGATGATCCTCTGGCCGTGGTTGACGAGCGTCTTCGGGTTCGCGGAATAGA<br>AGGCTTACGGGTTGCCGACGCGTCGATTATGCCCACGCTGGTGGGAGGCAA<br>TACTAATGCGCCAACAATTATGATCGGTGAGAAAGCCGCCGATATGATTCGT<br>GAGGATGCAAGACCTAAAGCCCAGGTGAGAGAAGCTGTTCGCTCCCCGGCG<br>TTCGCGGCATAA | MNHDLEFDYVIVGAGSA<br>GCAIAARLTEDPDVSVCL<br>LEAGGPDKSVLIHAPVGV<br>VAMLPTKINNYAYQTVPQ<br>PGLNGRRGYQPRGKTLGG<br>SSSINAMLYVRGHRHDYD<br>HWAALGNPGWSYDEVLP<br>LFRRSENNEHFRNDPYHG<br>SGGPLNVAELRSPSELDEA<br>FVQAAVNHGIPRTPDYNG<br>AEQFGSFRYQVTQKNGER<br>CSAAKGYLTPNLSRPNLK<br>VITHAVSARVLLDGRRAR<br>GVAYYQGNELTEVRARR<br>EVVLAGGAFGSPQLLMLS<br>GIGPAEELRKHGITVAHAL<br>PGVGQNLQDHIDYVQTY<br>MASSRTETFGVSARGGVR<br>MTKGILEWRRERTGPVTS<br>PFAEAGAFFRSSPAVPVPD<br>LQLVFVVAIVDDHARKM<br>HLGHGLSCHLTLLRPKSR<br>GTVGIASSDPREAPVIDPR<br>FFSDPADMPLMLDGAAK<br>MQAILEDRALARYRNGK<br>MLYTVRADDRSGLEADIR<br>NRADTQYHPVGSCKMGP<br>ADDPLAVVDERLRVRGIE<br>GLRVADASIMPTLVGGNT<br>NAPTIMIGEKAADMIRED<br>ARPKAQVREAVRSPAFAA |
| pZE-LaAAO<br>(WP_13851627<br>4.1) | ATGGAGTTCGATTTTGTAATCGTTGGCGGAGGTTCAAGCGGTGCTACCCTGG<br>CGGCACGTCTCAGCGAGGACAGTTCTGTCACAGTGTGCCTACTTGAAGCCGG<br>TGGTCGGGGTGATAATAGCCTGATACGTACCCCGGCCGCAATGGTTGCTATG<br>GTCCCGGGACACGGGAAATTGAATAACTGGGCATTTAACACAGTCCCTCAG<br>CCCGGTCTGAACGGCCGTATAGGATATCAACCTCGCGGAAAAGCACTAGGA<br>GGTTCCTCCGCCATTAACGCAATGCTGTACATTCGTGGCCAAAGGCAAGATT<br>ATGACGGTTGGGCGAATCTGGGCTGCGATGGCTGGGATTGGGACAGCGTTC<br>TGCCCTATTTCAAGGATGCCGAGAACAATGAACGCGGTGCGGACCCATTTCA<br>TGGCGCTAGTGGTCCACTGCATGTTTCGGACCAGAATTCGCCGCGCCCTGTG<br>ACAAGGGCCTTCGTTGAAGCAGCAAAAGCTTGGGGTCTGCCTGAGCAGCAA<br>GACTTCAATACTGGTGATAACGAAGGGACCGGCTTGTATCAGGTGACTCAGT<br>TTCACGATCCTAATAAACACGGCGAACGCTGTAGTGCCGCCGCGGCCTATCT<br>ACATCCGATCATGACAGAGCGGTCAAATCTTACGGTTCTCACCAATGCCCAT<br>GCATGTAGAATCCTACTGGAAAATCAGCGAGCTAAGGGCGTATTTTACCGCC<br>ATTCAGGCAAAGAGTTTTTGGTAAAAGCTCGCCGTGAAGTTATTGTGAGTGC<br>GGGTGCCTTCGGAAGCCCACAGCTGTTACAGCTGAGCGGGGTTGGCCGTCCT | MEFDFVIVGGGSSGATLA<br>ARLSEDSSVTVCLLEAGG<br>RGDNSLIRTPAAMVAMVP<br>GHGKLNNWAFNTVPQPG<br>LNGRIGYQPRGKALGGSS<br>AINAMLYIRGQRQDYDG<br>WANLGCDGWDWDSVLP<br>YFKDAENNERGADPFHG<br>ASGPLHVSDQNSPRPVTR<br>AFVEAAKAWGLPEQQDF<br>NTGDNEGTGLYQVTQFH<br>DPNKHGERCSAAAAYLHP<br>IMTERSNLTVLTNAHACRI<br>LLENQRAKGVFYRHSGKE<br>FLVKARREVIVSAGAFGSP<br>QLLQLSGVGRPQDITPYGI |

| | | |
|---|---|---|
| | CAGGACATTACCCCTTATGGTATTTCCATGGTTCATGAGCTGGCTGGAGTGG<br>GTCAAAATATGCAGGATCACTTAGACTTCACATTAGCGTTCAAATCTCTTGA<br>CACCGATAATTTTGGTTTAGGGCTGGCCGGCGCTTTAGGATTATTCAAGCAC<br>CTGACAAGCTGGCGGCGTAACGGAACTGGCATGTTGTCAAGCCCGTTCGCTG<br>AAGGAGCAGCGTTTCTGAAGTCCTCTAAGAGTATTGACCGCGCCGACCTTCA<br>GCTTCATTTTGTAATAAGTATCGTTGAGGACCATGCCCGCAAATTGCATTCT<br>GGTTATGGATTTAGTTGCCATGTTTGTGCGCTGCGGCCTTATAGCCGTGGAG<br>AGGTCTTTCTACAGTCGGCCGATCCACTGGACGACCCCGGAATTGATCCTAA<br>ATTTTTATCCGATCATCGTGATCTGGAAACCCTGATTAAGGGCGCAAAAATC<br>ACACGGGAAATTCTAATGCAGAAACCCCTGGAAAACTATCGCCATAAAGAA<br>CTTTTCGACGTTCATGAGGGCATGTCTGACTCCCAATGGGAGAGTAAAATTC<br>GCGCTCGTGCAGATACGATATATCACCCGGTTGGCACCTGTAAGATGGGCAC<br>GGATACCATGTCTGTGGTGGACGCACAACTGCGCGTTCACGGATTACAGGG<br>CCTGCGAGTCGTTGACGCCAGTGTTATGCCGACCTTAGTGTCAGGGAACACG<br>AATGCCCCCTCAATTATGATAGCTGAAAAAGCTGCTGACATGATACTGGGCA<br>AAAATAGAATAACTAAAACCTCCACGAGTCCGCAGATCAACGAGAAAGAAA<br>TGAGTCATGTTTAA | SMVHELAGVGQNMQDHL<br>DFTLAFKSLDTDNFGLGL<br>AGALGLFKHLTSWRRNG<br>TGMLSSPFAEGAAFLKSS<br>KSIDRADLQLHFVISIVED<br>HARKLHSGYGFSCHVCAL<br>RPYSRGEVFLQSADPLDD<br>PGIDPKFLSDHRDLETLIK<br>GAKITREILMQKPLENYR<br>HKELFDVHEGMSDSQWE<br>SKIRARADTIYHPVGTCK<br>MGTDTMSVVDAQLRVHG<br>LQGLRVVDASVMPTLVSG<br>NTNAPSIMIAEKAADMIL<br>GKNRITKTSTSPQINEKEM<br>SHV |
| pZE-VbAAO (MDQ0039797.1) | ATGGAATTCGATTATGTCATCGTGGGGGGCGGGTCCGGAGGAGCAACACTG<br>GCGAGTCGTTTAAGCGAAGATCCCGGGGTAACCGTATGCTTAATCGAGGCA<br>GGTGGGGACGGTCGCGGTATTTTAGTACGTGCGCCGGCTGCAACTGTTGCGA<br>TGTTACCAGGGAGACCGCCAATCAATAATTATGCATATAAAACGGTCCCTCA<br>GCCCGGTTTGGGAGGTAGGTCTGGCTATCAGCCACGTGGTCGTGGGTTAGGA<br>GGTTCCAGCGCCATAAATGCCATGCTCTATGTCCGAGGTCATCGTGACGATT<br>ACGATGATTGGGCTCGGGCGGGATGTGAGGGATGGTCTTTTGACGAAGTAC<br>TGCCGTATTTTAAACGTGCTGAAGGCAATGAGCGCGGAGAATCAGCGCTGC<br>ACGGAGCGGGCGGACCATTACAAGTCTCCGAGCAACAGTCCCCGAGACCTA<br>TTACTGAAGACTTCATACGTGCGGCCGCCGAATGCGGTATACCGCGTAATGA<br>TGACTTCAACGGCGCCGAACAAGAGGGTGCGGGGTTATACCAGGTAACTCA<br>GTTCCACGGTGGCACGAAAAATGGAGAACGTTGCAGTGCTGCCGCGGCTTA<br>TCTGCATCCTGCGATGCATGCACGTCCAAATCTTACGGTGCTGACTGGCGCC<br>CAGGCGTTGCGTGTGGTATTAGATGGTAAGCGTGCCACTGGGGTGGAAGTC<br>CGCCGTGGTGGGACAACCGAAGTTATCCGTGCCCATCGGGAGGTGGCACTG<br>TGTGGGGGTGCATTTAATAGTCCTCAGCTACTCATGTTGTCAGGTATCGGAG<br>ATCCTGCAGAATTAGGTCGACACGGCATCGCAGTGCGACATGCACTTCCTGG<br>TGTTGGGCAAAACCTCCAGGATCATACTGACTTCATTCTTGCATACACTTCA<br>AAGGATATTGATCTCTTTGGGATTGGGGTCAAAGCGGGCCTGAAATTGATGA<br>AAGCAATTTTCGAGTGGCGAAAGAGCGGTAAGGGTCTTGTAGCAACGCCGT<br>TCGCAGAGGGCGGCGCATTCATAAAGAGTTCTCCGGAGCTGCGCCGTCCTG<br>ACCTGCAGCTTCACTTTGTCATCGCCATAACAGACGATCATGCACGCAAACT<br>ACATATGGGGTTTGGATTTAGCTGTCACGTTTGCGTACTGCGTCCCAAAGGT<br>AGGGGAGACGTGCGTCTGAACGACGCGAACCCACTATCAGCCCCCCGCATA<br>GATCCGAGGTTTCTTAGTGATGCTGAAGATATGGCTTTACTGCTGCAGGGCG<br>TGAAAAAGATGCGCGAAATATTACGTGCTCCAGCCCTGCAAAAATATCGCC<br>ACCGCGAGGTCTACACGGCTGATGCCCATACCGACGAGCAGCTAACGCAGC<br>ACATCCGAGCCCGGGCTGATACGATTTACCATCCTGTTGGCACTTGCAAAAT<br>GGGCGTAGATGCGATGGCCGTCGTGGATGCACAGCTCCGTGTTCACGGCATT<br>GAAAACTTAAGGGTTGTCGATGCTAGCGTGATGCCGACTCTGATTGGGGGC<br>AATACAAATGCCCCTACCATTATGATCGCAGAACGCGCCGCTGACTGGATGC<br>GTGGGCCTATTAAGTTTGACAGAGCGGTGGTCGACGCCAGAGCACCGGCAA<br>CAATACCTGCCCCGTCCGTGCCGCTGGAAGCCGCGGTTGGCGCAGTGCTGTC<br>TCCCGCTCCGGCAGCGGCGCGGTCAGGGATGTAA | MEFDYVIVGGGSGGATLA<br>SRLSEDPGVTVCLIEAGGD<br>GRGILVRAPAATVAMLPG<br>RPPINNYAYKTVPQPGLG<br>GRSGYQPRGRGLGGSSAI<br>NAMLYVRGHRDDYDDW<br>ARAGCEGWSFDEVLPYFK<br>RAEGNERGESALHGAGGP<br>LQVSEQQSPRPITEDFIRA<br>AAECGIPRNDDFNGAEQE<br>GAGLYQVTQFHGGTKNG<br>ERCSAAAAYLHPAMHAR<br>PNLTVLTGAQALRVVLDG<br>KRATGVEVRRGGTTEVIR<br>AHREVALCGGAFNSPQLL<br>MLSGIGDPAELGRHGIAV<br>RHALPGVGQNLQDHTDFI<br>LAYTSKDIDLFGIGVKAGL<br>KLMKAIFEWRKSGKGLV<br>ATPFAEGGAFIKSSPELRR<br>PDLQLHFVIAITDDHARKL<br>HMGFGFSCHVCVLRPKGR<br>GDVRLNDANPLSAPRIDP<br>RFLSDAEDMALLLQGVK<br>KMREILRAPALQKYRHRE<br>VYTADAHTDEQLTQHIRA<br>RADTIYHPVGTCKMGVD<br>AMAVVDAQLRVHGIENL<br>RVVDASVMPTLIGGNTNA<br>PTIMIAERAADWMRGPIK<br>FDRAVVDARAPATIPAPS<br>VPLEAAVGAVLSPAPAAA<br>RSGM |
| AAO C- term his tag | GGTAGCAGTCACCATCATCACCACCATTAA | GSSHHHHHH* |
| pACYC-CvTA-AlaDH | CvTA:<br>ATGGGCAGCAGCCATCACCATCATCACCACAGCCAGGATCCGAATTCGATG<br>CAGAAGCAGCGTACAACATCGCAATGGCGCGAACTTGACGCCGCTCATCAC<br>CTGCATCCCTTCACCGATACCGCCTCCCTTAACCAGGCCGGCGCGCGCGTGA<br>TGACACGTGGAGAAGGGGTGTATTTGTGGGACTCGGAGGGAAATAAAATCA<br>TCGACGGTATGGCTGGATTATGGTGTGTGAACGTTGGCTACGGTCGTAAGGA<br>CTTTGCCGAAGCGGCCCGTCGTCAGATGGAAGAATTACCGTTCTACAATACT<br>TTTTTCAAAACAACCCATCCTGCGGTCGTAGAGTTATCTTCATTATTGGCGG<br>AAGTCACTCCAGCAGGGTTTGACCGCGTGTTTTATACAAATAGTGGATCAGA<br>ATCGGTTGACACAATGATCCGTATGGTCCGTCGTTACTGGGACGTCCAAGGC<br>AAACCGGAGAAGAAGACGTTAATCGGCCGCTGGAATGGTTATCACGGTTCG<br>ACCATTGGAGGTGCATCTCTTGGGGGCATGAAGTATATGCATGAGCAGGGT<br>GATTTGCCTATCCCTGGCATGGCGCACATCGAACAACCGTGGTGGTATAAGC<br>ACGGTAAAGACATGACGCCGGACGAGTTTGGAGTTGTCGCTGCGCGTTGGTT<br>GGAAGAGAAGATCCTGGAAATTGGGGCGGACAAGGTAGCCGCCTTCGTAGG<br>AGAACCAATCCAAGGTGCCGGGGGAGTGATCGTCCCGCCAGCTACCTATTG<br>GCCCGAGATCGAGCGCATTTGCCGTAAATATGACGTATTGCTGGTTGCAGAT | CvTA:<br><br>MGSSHHHHHHSQDPNSM<br>QKQRTTSQWRELDAAHH<br>LHPFTDTASLNQAGARVM<br>TRGEGVYLWDSEGNKIID<br>GMAGLWCVNVGYGRKD<br>FAEAARRQMEELPFYNTF<br>FKTTHPAVVELSSLLAEVT<br>PAGFDRVFYTNSGSESVD<br>TMIRMVRRYWDVQGKPE<br>KKTLIGRWNGYHGSTIGG<br>ASLGGMKYMHEQGDLPIP<br>GMAHIEQPWWYKHGKD<br>MTPDEFGVVAARWLEEKI<br>LEIGADKVAAFVGEPIQG |

| | | |
|---|---|---|
| | GAGGTAATTTGTGGCTTCGGGCGCACCGGGGAGTGGTTCGGGCACCAACAT<br>TTCGGTTTTCAGCCGGACTTATTTACGGCGGCGAAGGGTTTAAGCTCAGGTT<br>ATTTACCGATTGGGGCTGTGTTTGTGGGCAAGCGTGTTGCCGAAGGCTTAAT<br>CGCGGGAGGCGACTTTAATCACGGATTCACATACTCTGGACACCCGGTTTGT<br>GCCGCAGTAGCTCACGCGAATGTAGCCGCATTACGTGACGAGGGAATCGTC<br>CAGCGTGTGAAGGACGATATCGGCCCTTATATGCAGAAGCGCTGGCGCGAG<br>ACTTTTTCACGTTTTGAGCACGTAGACGATGTGCGTGGCGTAGGCATGGTAC<br>AGGCCTTTACCTTAGTCAAAAATAAAGCTAAGCGCGAGTTGTTCCCAGACTT<br>TGGCGAAATCGGAACGTTGTGTCGCGATATCTTTTTTCGCAATAATCTTATC<br>ATGCGCGCTTGCGGGGATCATATTGTAAGTGCCCCGCCATTGGTGATGACTC<br>GTGCCGAGGTAGATGAGATGTTAGCAGTCGCAGAGCGCTGCCTTGAGGAGT<br>TTGAGCAAACATTAAAAGCTCGCGGACTTGCCTGA<br>AlaDH:<br>ATGGGCAGCAGCCATCACCATCATCACCACATGATCATAGGGGTTCCTAAA<br>GAGATAAAAAACAATGAAAACCGTGTCGCATTAACACCCGGGGGCGTTTCT<br>CAGCTCATTTCAAACGGCCACCGGGTGCTGGTTGAAACAGGCGCGGGCCTT<br>GGAAGCGGATTTGAAAATGAAGCCTATGAGTCAGCAGGAGCGGAAATCATT<br>GCTGATCCGAAGCAGGTCTGGGACGCCGAAATGGTCATGAAAGTAAAAGAA<br>CCGCTGCCGGAAGAATATGTTTATTTTCGCAAAGGACTTGTGCTGTTTACGT<br>ACCTTCATTTAGCAGCTGAGCCTGAGCTTGCACAGGCCTTGAAGGATAAAGG<br>AGTAACTGCCATCGCATATGAAACGGTCAGTGAAGGCCGGACATTGCCTCTT<br>CTGACGCCAATGTCAGAGGTTGCGGGCAGAATGGCAGCGCAAATCGGCGCT<br>CAATTCTTAGAAAAGCCTAAAGGCGGAAAAGGCATTCTGCTTGCCGGGGTG<br>CCTGGCGTTTCCCGCGGAAAAGTAACAATTATCGGAGGAGGCGTTGTCGGG<br>ACAAACGCGGCGAAAATGGCTGTCGGCCTCGGTGCAGATGTGACGATCATT<br>GACTTAAACGCAGACCGCTTGCGCCAGCTTGATGACATCTTCGGCCATCAGA<br>TTAAAACGTTAATTTCTAATCCGGTCAATATTGCTGATGCTGTGGCGGAAGC<br>GGATCTCCTCATTTGCGCGGTATTAATTCCGGGTGCTAAAGCTCCGACTCTT<br>GTCACTGAGGAAATGGTAAAACAAATGAAACCCGGTTCAGTTATTGTTGAT<br>GTAGCGATCGACCAAGGCGGCATCGTCGAAACTGTCGACCATATCACAACA<br>CATGATCAGCCAACATATGAAAAACACGGGGTTGTGCATTATGCTGTAGCG<br>AACATGCCAGGCGCAGTCCCTCGTACATCAACAATCGCCCTGACTAACGTTA<br>CTGTTCCATACGCGCTGCAAATCGCGAACAAAGGGGCAGTAAAAGCGCTCG<br>CAGACAATACGGCACTGAGAGCGGGTTTAAACACCGCAAACGGACACGTGA<br>CCTATGAAGCTGTAGCAAGAGATCTAGGCTATGAGTATGTTCCTGCCGAGAA<br>AGCTTTACAGGATGAATCATCTGTGGCGGGTGCTTAA | AGGVIVPPATYWPEIERIC<br>RKYDVLLVADEVICGFGR<br>TGEWFGHQHFGFQPDLFT<br>AAKGLSSGYLPIGAVFVG<br>KRVAEGLIAGGDFNHGFT<br>YSGHPVCAAVAHANVAA<br>LRDEGIVQRVKDDIGPYM<br>QKRWRETFSRFEHVDDVR<br>GVGMVQAFTLVKNKAKR<br>ELFPDFGEIGTLCRDIFFRN<br>NLIMRACGDHIVSAPPLV<br>MTRAEVDEMLAVAERCL<br>EEFEQTLKARGLA*<br><br>AlaDH:<br>MGSSHHHHHHMIIGVPKE<br>IKNNENRVALTPGGVSQLI<br>SNGHRVLVETGAGLGSGF<br>ENEAYESAGAEIIADPKQV<br>WDAEMVMKVKEPLPEEY<br>VYFRKGLVLFTYLHLAAE<br>PELAQALKDKGVTAIAYE<br>TVSEGRTLPLLTPMSEVA<br>GRMAAQIGAQFLEKPKGG<br>KGILLAGVPGVSRGKVTII<br>GGGVVGTNAAKMAVGL<br>GADVTIIDLNADRLRQLD<br>DIFGHQIKTLISNPVNIADA<br>VAEADLLICAVLIPGAKAP<br>TLVTEEMVKQMKPGSVIV<br>DVAIDQGGIVETVDHITTH<br>DQPTYEKHGVVHYAVAN<br>MPGAVPRTSTIALTNVTV<br>PYALQIANKGAVKALAD<br>NTALRAGLNTANGHVTY<br>EAVARDLGYEYVPAEKA<br>LQDESSVAGA* |

### Supplementary Figures

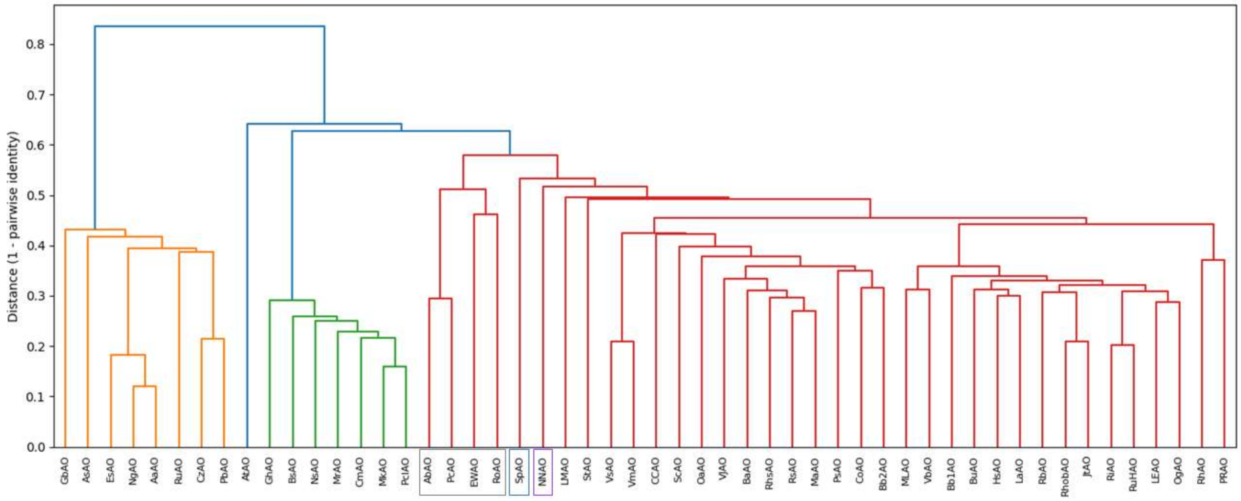


**Supplementary Figure 1**. Hierarchical clustering analysis on candidate AAOs. The coloring indicates four very distinct clusters (orange, blue, green, and red) along with the emergence of three additional small clusters (boxed in gray, blue, and purple) after specification of seven total clusters. The red cluster contains homologs with the highest mean expression level.

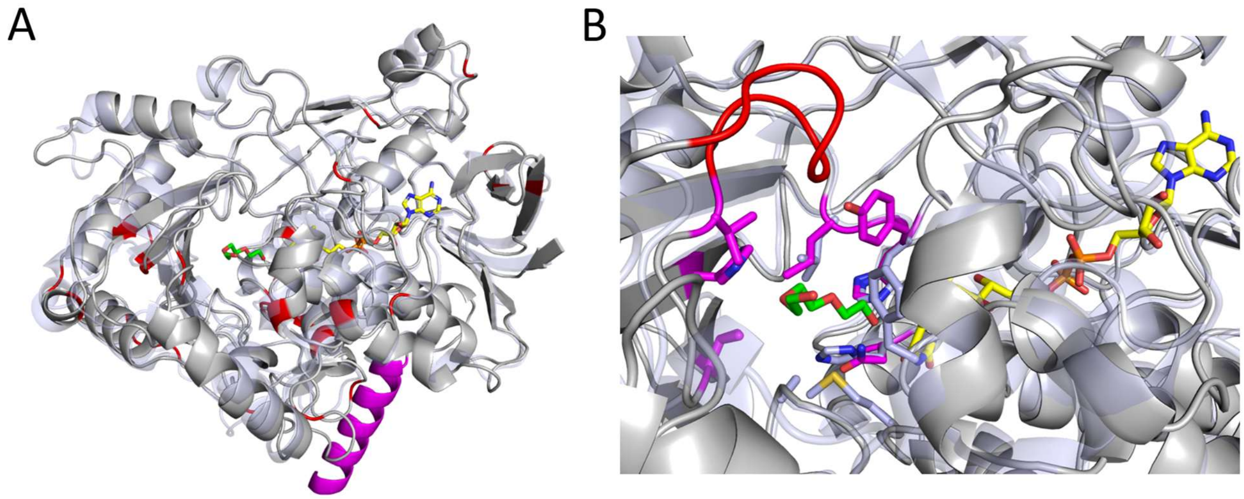


**Supplementary Figure 2.** Comparisons of the AlphaFold-predicted structure of BuAAO (dark gray) after alignment to the published crystal structure of ShAAO (PDB ID 8RPG, transparent light blue). (A) Global structural differences in Cα atom position for each alignment position using an automated analysis, followed by coloring the top 30 most divergent positions in red. The alcohol substrate bound in the crystal structure is colored green and the FAD co-factor is colored yellow. Highlighted in magenta is a C-terminal alpha-helix that is unique to BuAAO. Note that this analysis of divergent backbone positions omits loop regions. (B) View of the difference in binding pocket geometries from outside the binding pocket, where an extended and shifted loop region belonging to BuAAO (highlighted in red) appears to influence binding pocket access.
